# Organic-acid-derived feather hydrolysate drives soil microbiome succession toward copiotrophic and putatively plant-beneficial taxa

**DOI:** 10.64898/2026.09.22.753663

**Authors:** Jan Veselský, Martina Dlasková, Radim Čegan, Šárka Kobzová, Zdeněk Kubát, Lenka Vaňková, Roman Hobza, Olga Šolcová, Lucie Kajan Grodecká

**Affiliations:** Department of Plant Developmental Genetics, Institute of Biophysics of the Czech Academy of Sciences, Brno, Czech Republic; Department of Experimental Biology, Faculty of Science, Masaryk University, Brno, Czech Republic; Institute of Chemical Process Fundamentals of the Czech Academy of Sciences, Prague, Czech Republic; Institute of Experimental Botany of the Czech Academy of Sciences, Centre of the Region Haná for Biotechnological and Agricultural Research, Olomouc, Czech Republic; Department of Chemistry and Biochemistry, Mendel University in Brno, Brno, Czech Republic

**Keywords:** feather hydrolysate, microbial community dynamics, copiotrophs, rrn-copy number, PGPR, phytopathogen suppression, keratin

## Abstract

As circular agricultural inputs are increasingly sought, feather hydrolysate (FH), derived from keratin-rich poultry waste, has emerged as a promising fertilizer and biostimulant. However, how FH shapes soil microbiome succession over time, and what functional potential this restructuring carries, remains poorly understood. Here, we used multi-marker high-throughput sequencing of 16S, 18S, and ITS rDNA to track soil microbial communities over 30 days following FH amendment, with sampling at days 0, 1, 4, 14, and 30. FH induced a rapid and reproducible microbial succession characterized by an early decline in alpha diversity and a pronounced day-1 bloom of copiotrophic taxa, as inferred from predicted rrn operon copy-number profiles. This initial response was accompanied by enrichment of bacterial genera previously associated with plant growth-promoting traits, including pathogen suppression, nutrient solubilization, siderophore production, phytohormone synthesis, and ACC deaminase activity. Although these groups gradually declined after the initial peak, several remained at significantly elevated abundances throughout the 30-day experimental period. In parallel, FH-treated soils showed a sustained reduction in the cumulative relative abundance of genera annotated as putative phytopathogens. By integrating temporal community profiling, rrn-based life-strategy inference, and literature-derived functional annotation, we show that FH acts as a strong ecological filter, shifting the soil microbiome toward copiotrophic and putatively plant-beneficial taxa. These findings provide a temporal framework for understanding FH-mediated microbiome restructuring and highlight the need for future studies linking inferred microbial functional potential to measured disease incidence, nutrient fluxes, and plant performance across diverse soil contexts.

**Graphical abstract:** 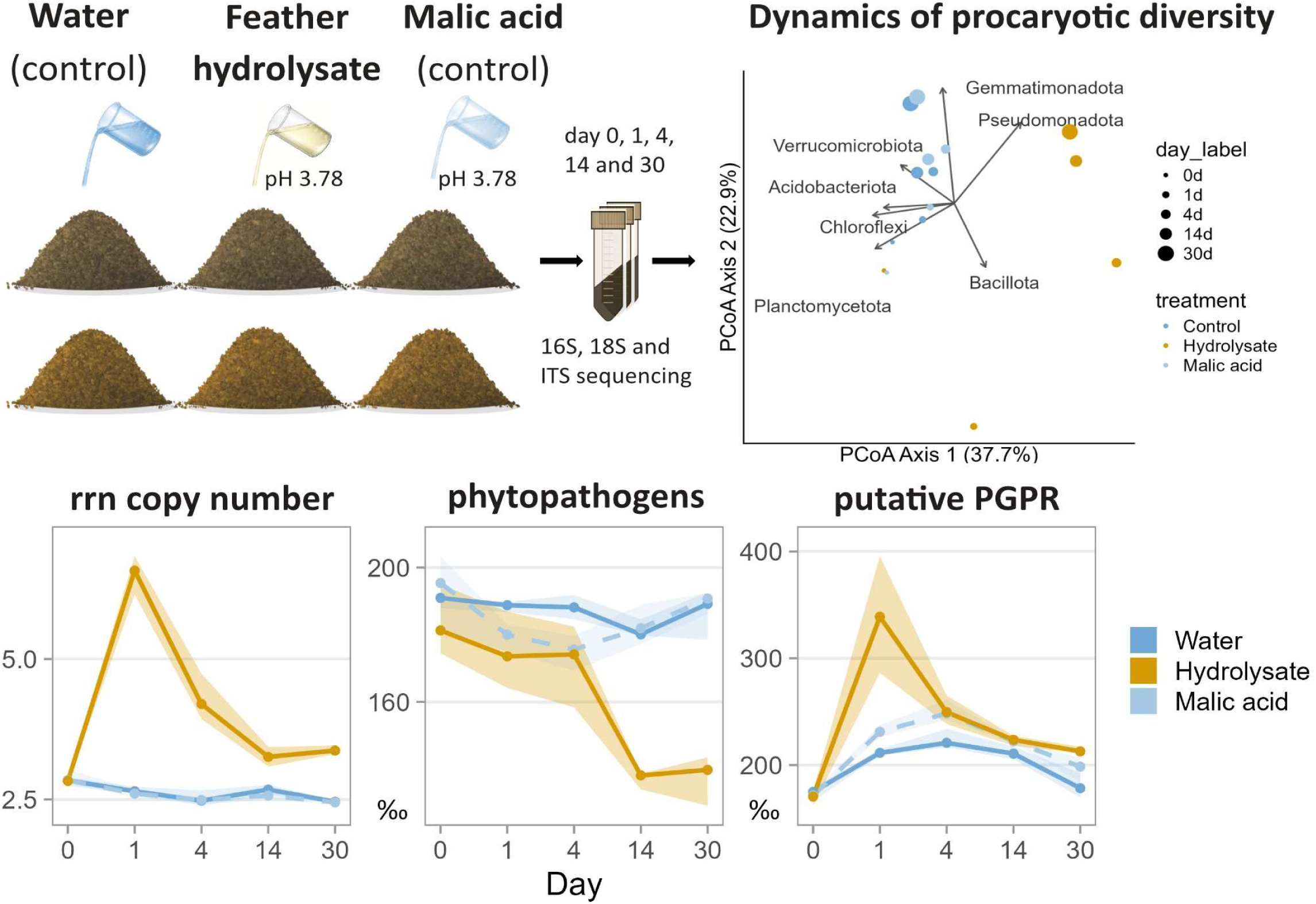

## 1. Introduction

Protein hydrolysates (PHs), promising products of organic waste, have recently gained significant attention as potential biostimulants and fertilizers (Halpern et al., 2015). Derived from protein-rich materials such as poultry feathers, fish residues, leather waste, legume biomass, etc., they are mainly composed of free amino acids, oligopeptides and polypeptides (Colla et al., 2015; Jagadeesan et al., 2023). Being rich in carbon and nitrogen, PHs serve as an energy and nutrient source for both soil microbiota and plants. However, viewing PHs merely through a nutritional lens would be a reductive oversimplification. PHs not only benefit particular microbial taxa, thereby influencing microbial community balance and the ecological processes it supports, but they can also regulate the production of secondary metabolites, including antimicrobial compounds.

To this end, PHs and other amino acid-rich substrates have been shown to suppress phytopathogen load and plant diseases through induced systemic resistance, microbe-dependent processes, or a combination thereof (Andreo-Jimenez et al., 2021; Postma and Schilder, 2015; Russ et al., 2024). Microbes can suppress pathogens either directly, through production of antimicrobial compounds, or indirectly, through the sequestration of essential nutrients (Philippot et al., 2013). In general, nitrogen availability can both stimulate and suppress antibiotic production (Romero-Rodríguez et al., 2018). Individual amino acids, such as Gly, Lys, Arg, Trp, Glu, Asn, and Gln, have been shown to promote production of antibiotics (Kim et al., 2021) or volatile organic compounds (VOCs) with antimicrobial properties (Papazlatani et al., 2025), where amino acids can serve as both precursors and regulators. While existing literature mostly focuses on a few particular pathogens, our knowledge regarding the broader impact of PHs on phytopathogen communities and their temporal dynamics remains rather limited.

PHs have been repeatedly shown to promote plant growth by enhancing germination frequency, root biomass, leaf chlorophyll content, and the secondary metabolites content in fruits (Colla et al., 2014; Gurav and Jadhav, 2013; Jagadeesan et al., 2023; Kaur et al., 2021). In these cases, beyond the direct nutritional and regulatory effects of amino acids and peptides on plants, a significant microbial contribution is expected (Philippot et al., 2013). In fact, amino acids are a crucial component of root exudates, which plants utilize to recruit and sustain beneficial microbiota, collectively known as plant growth-promoting rhizobacteria (PGPR) (Beauregard, 2015; Upadhyay et al., 2022; Upadhyay and Kumar, n.d.). These bacteria provide plants with multiple services, such as nitrogen fixation, nutrient solubilization, and the production of phytohormones or the stress-alleviating enzyme ACC deaminase, in addition to their role in phytopathogen biocontrol (Berendsen et al., 2012; Glick, 2012; Feng et al., 2023). Indeed, PH amendment has been linked to the enhancement of several PGPR-related functions in soil (Jagadeesan et al., 2023; Paul et al., 2013).

Despite these promising benefits, the efficacy of PHs can be inconsistent, often due to chemical characteristics of the PH that stem from the selected protein source and the production process (Colla et al., 2017, 2015; Costa et al., 2024). For feather keratin, a highly recalcitrant scleroprotein, traditional inorganic acid hydrolysis often leads to the irreversible degradation of essential amino acids and high salinity in the final product, limiting its agricultural utility (Ben Hamad Bouhamed and Kechaou, 2017). Conversely, organic acid hydrolysis (e.g., using malic or formic acid) offers a gentler, more sustainable alternative that preserves delicate amino acid profiles and yields salt-free products suitable for direct soil application (Hanika et al., 2021).

In this work, we aimed to decipher the dynamics of soil microbiome shifts following the amendment of a chicken feather hydrolysate (FH) prepared via mild malic-acid hydrolysis. We hypothesized that this high-quality nutrient pulse would rapidly select for copiotrophic taxa and subsequently reshape the community possibly toward plant-beneficial functions.

Through extensive literature-based annotation of the detected genera, we demonstrate that the suppression of most phytopathogens remained stable throughout the whole experimental period. Further, we show an increase in the abundance of multiple putative PGPR groups with the inferred potential for pathogen suppression, nutrient solubilization, and the production of siderophores and phytohormones. While this enhancement gradually declined after peaking at day 1, the increase remained distinct even at day 30.

## 2. Methods

### 2.1. Experimental soil

Two independent soil samples were collected from an agricultural field near Lukavec, Czechia, on August 6, 2024. The elemental composition of the soil was determined by acid digestion with aqua regia enhanced by microwave treatment followed by analysis by inductively coupled plasma optical emission spectroscopy (ICP-OES). The content of total carbon and total nitrogen was analysed using FLASH 2000 (ThermoFisher Scientific Inc., Waltham, MA, USA) organic elemental analyzer. For each measurement 1-3 mg of measured sample was placed in a soft tin container and introduced into a quartz reactor filled with copper oxide and electrolytical copper. The reactor was heated to 950 °C and a small volume of oxygen was injected along with the sample. The gases released by the combustion of the sample were measured by the machine’s in-built detector. The relative carbon and nitrogen content was determined by comparing the resulting spectra to BBOT CHNS standard (ThermoFisher Scientific). Each soil sample was analyzed in triplicate, and the results are presented as mean values (Tab. S1). The relative standard deviation among replicates did not exceed 10% for any measured element.

### 2.2. Feather hydrolysate

Poultry feather hydrolysate was prepared as was described earlier (Hanika et al., 2021). The dry matter content of the hydrolysate was 2.4 wt.%. Elemental analysis of the dry matter was performed using a scanning electron microscope (SEM, Tescan Indusem) equipped with an energy-dispersive X-ray spectrometer (EDS, SDD XFlash® 5010, Bruker; energy resolution 125 eV). Measurements of ten replicate samples were carried out at an accelerating voltage of 15 kV. Table S2 shows the average values with a (95%) margin of error of up to 5%. Minor element content in the hydrolysate form is depicted in table S3. Amino acid content was measured with HPLC/MS (Tab. S4). Total amino acid content was 1878 mg L^-1^. Total low molecular weight peptide content was 20 948 mg L^-1^.

### 2.3. Experimental design

Soil samples were air-dried and sieved to <1.25 mm. A batch experiment was conducted in triplicate using 250 g of soil in 800 mL beakers. Three treatments (60 mL total volume) were applied: (i) distilled water (control); (ii) a hydrolysate mixture (20 mL hydrolysate [pH 3.59] + 40 mL water; final pH 3.78); and (iii) a malic acid solution adjusted to pH 3.77. Solutions were added in 10 mL increments and mixed thoroughly. Beakers were sealed with Durafilm to minimize evaporation, and 5 mL of distilled water was added on day 14 to maintain moisture. Samples were collected on days 0, 1, 4, 14, and 30, immediately flash-frozen in liquid nitrogen, and stored at −20 °C. On days 20 and 30, soil pH was determined in a 0.01 M CaCl₂ suspension (1:5 w/v) after 60 min of shaking followed by 60 min of sedimentation (Tab. S5**Chyba! Nenalezen zdroj odkazů.Chyba! Nenalezen zdroj odkazů.**).

### 2.4. Soil microbiome analysis

Total DNA was extracted using the NucleoSpin® Soil kit (Macherey-Nagel). Quality and yield were verified via Qubit (ThermoFisher Scientific) and control PCR. Target regions of 16S, 18S and ITS rDNA were amplified (Tab. S6), and the resulting sequencing libraries were sequenced on the AVITI platform (2x300 bp paired-end).

Raw paired-end reads were joined using PEAR v0.9.11 (Zhang et al., 2014), demultiplexed based on specific primers and subsequently adapter-trimmed, filtered by expected amplicon length and quality-filtered using Trimmomatic v0.32 (Bolger et al., 2014). For amplicon of ITS region, unjoined reads with primer sequence present were included in the dataset. Taxonomic composition of the microbial community was performed using Kraken2 (Wood et al., 2019), individually for each amplicon, utilizing the SILVA database v138-NR99 (Quast et al., 2013).

Taxonomic composition profiles produced by Kraken2 were imported into the R environment and processed using phyloseq (McMurdie and Holmes, 2015; “R: The R Project for Statistical Computing,” n.d.). For each sample, taxonomic profiles generated from individual amplicons were merged by averaging the relative abundance of each taxonomic feature across the relevant amplicons. If a given feature was not detected in any sample for a particular amplicon dataset, that amplicon was excluded from the abundance calculation for that feature, rather than being treated as zero. This procedure resulted in one combined taxonomic composition estimate per sample. The merging was performed separately for eukaryotic and prokaryotic amplicons. This combined dataset was further used to assess the relative abundance of microbial groups and beta-diversity of the community. For analyses that required both eukaryotic and prokaryotic profiles, relative counts of both datasets were summed.

Ecological function annotation of eukaryotes was done by UNITE 9 database (Abarenkov et al., 2024). Prokaryotic community was annotated based on available literature. List of specific taxa and their assigned function is available in table S7. Furthermore, in prokaryotic community, expected number of *rrn* gene copies was assigned based on rrnDB 5.1 (Stoddard et al., 2015). Genera with a predicted rrn copy number ≤2 were classified as oligotrophic and genera with ≥5 copies as copiotrophic; genera with an intermediate copy number (3–4) were not assigned to either category. Beta diversity was determined via the Bray-Curtis dissimilarity index and subsequent Principal Coordinate Analysis ordination using phyloseq package.

To assess alpha diversity of the microbial community, separate dataset was used, where prior to the merging, reads for each amplicon have been rarefied. Specific rarefaction depths are shown in table S8. Alpha diversity metrics were calculated using phyloseq package. For data analysis, tidyverse and coin packages (Hothorn et al., 2008; Wickham et al., 2019) were used and data visualization was done using ggplot2 package (Wickham, 2016).

### 2.5. Analysis of hydrolysate microbiota

To distinguish autochthonous soil microbiota from induced strains, near-full length SSU rRNA gene sequencing was performed using MinION platform (Oxford Nanopore Technologies), with SQK-LSK114 kit (Oxford Nanopore Technologies) for sequencing library preparation. The subsequent basecalling was performed by Dorado v 1.4.0 sup model (Oxford Nanopore Technologies), and the resulting file was converted to fastq format using samtools v1.14 (Li et. al., 2009). The resulting sequences were demultiplexed, filtered for minimal Phred score of q10 using Cutadapt v5.2 (Martin, 2011) and the adapter and primer sequences were removed. Demultiplexed and filtered reads were annotated using EMU pipeline with SILVA138.2 SSURefNR99 (Quast et al., 2013) as the reference database.

### 2.6. Statistics

Alpha diversity was calculated separately for each sample at the genus level from rarefied relative abundance data us-ing three metrics: observed richness (number of taxa with non-zero relative abundance), the Shan-non diversity index (H′ = −Σ pᵢ ln pᵢ), and the inverse Simpson index (1/Σ pᵢ²), where pᵢ is the relative abundance of taxon i within a given sample after excluding absent taxa and re-normalising to unit sum. Metrics were computed both for the amplicon-combined dataset (prokaryotic and eukaryotic taxa separately, Fig. 1, Fig. S1) and separately for each amplicon (Fig. S15–S17). Beta diversity was assessed using the Bray–Curtis dissimilarity index, and community composition was visualised by principal coordinates analysis (PCoA) implemented in the phyloseq R package. To quantify between-group differences, pairwise centroid distances were calculated as Euclidean distances in the two-dimensional ordination space, with each axis scaled by the square root of its relative proportion of variance explained, so that axes explaining more variance contribute proportionally more to the distance.

**Figure 1.**
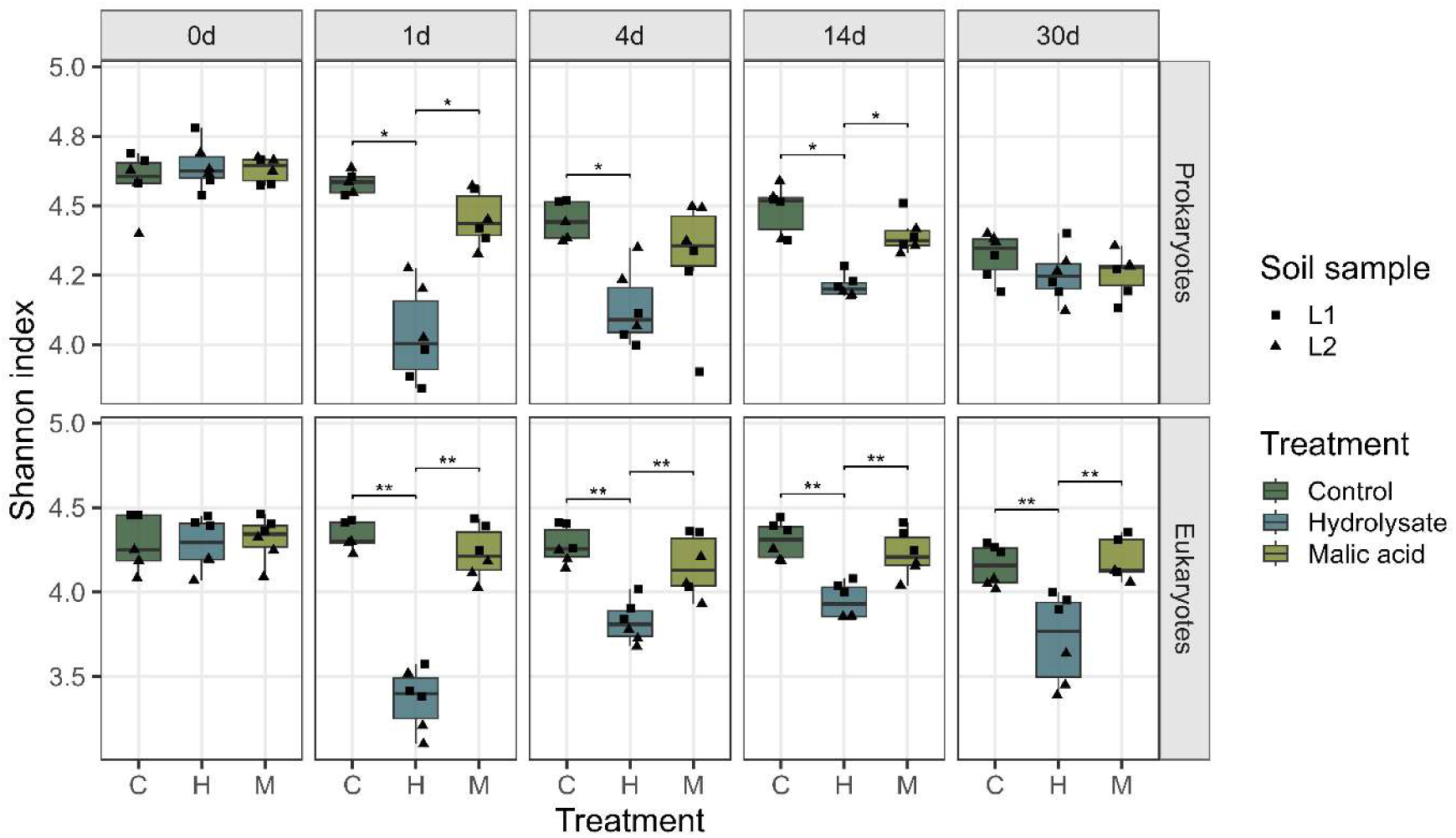
Alpha diversity of the soil microbial community 0–30 days after hydrolysate amendment. Boxes represent the inter-quartile range; whiskers indicate the range excluding outliers. Statistical significance (p < 0.05) is based on two-sided van Elteren test followed by Benjamini–Hochberg correction.

Pairwise differences between independent groups were evaluated using two-sided van Elteren tests with the soil sample as the blocking variable, followed by Benjamini–Hochberg correction. Whenever the figures depict prokaryotic and eukaryotic microbiota separately, the correction was likewise applied separately within each domain. In analyses where the L1 and L2 soil samples were assessed separately (e.g., Fig. S9), a standard two-sided Wilcoxon rank-sum test was used instead, as blocking by soil sample was not applicable within a single locality; correction was applied separately for each soil sample. Enrichment of soil saprotrophic genera among genera with significantly increased abundance was evaluated at each timepoint using two-sided Fisher’s exact tests on 2×2 contingency tables, followed by Benjamini–Hochberg correction applied separately for each comparison (hydrolysate vs. water control; hydrolysate vs. malic acid).

Differential abundance of individual taxa and functional groups between the hydrolysate treatment and each control was assessed at each sampling timepoint using the same two-sided van Elteren test framework described above, with Benjamini–Hochberg correction applied separately for each comparison (hydrolysate vs. control; hydrolysate vs. malic acid) across all taxa/functional groups and sampling days within that comparison.

## 3. Results and discussion

In this study, we evaluated the microbiome response of two soil samples to FH amendment, using water and malic acid as independent controls. Total soil DNA was extracted, and 16S, 18S, and ITS barcoding genes were sequenced at five time points: day 0 (immediately following treatment), and days 1, 4, 14, and 30. Monitoring of soil pH did not reveal any consistent trends (Tab. S5).

### 3.1. Microbiome diversity dynamics

Alpha diversity for both prokaryotes and eukaryotes declined progressively across all experimental variants (Fig. 1, Fig. S1), consistent with shifts typically observed during laboratory soil incubation (Zegeye et al., 2019). Hydrolysate amendments induced the most pronounced initial reduction in diversity on day 1; however, this effect gradually attenuated in both domains over time. By day 30, diversity values converged toward control levels, a trend more markedly observed in prokaryotic than eukaryotic communities. These findings are further supported by Bray-Curtis dissimilarity analysis (Fig. 2, Fig. S2): within the hydrolysate-treated group, day 1 showed the highest average pairwise dissimilarity to the other post-amendment timepoints (days 4, 14, and 30; Table S9), consistent with the most pronounced community shift occurring at this early stage.

**Figure 2.**
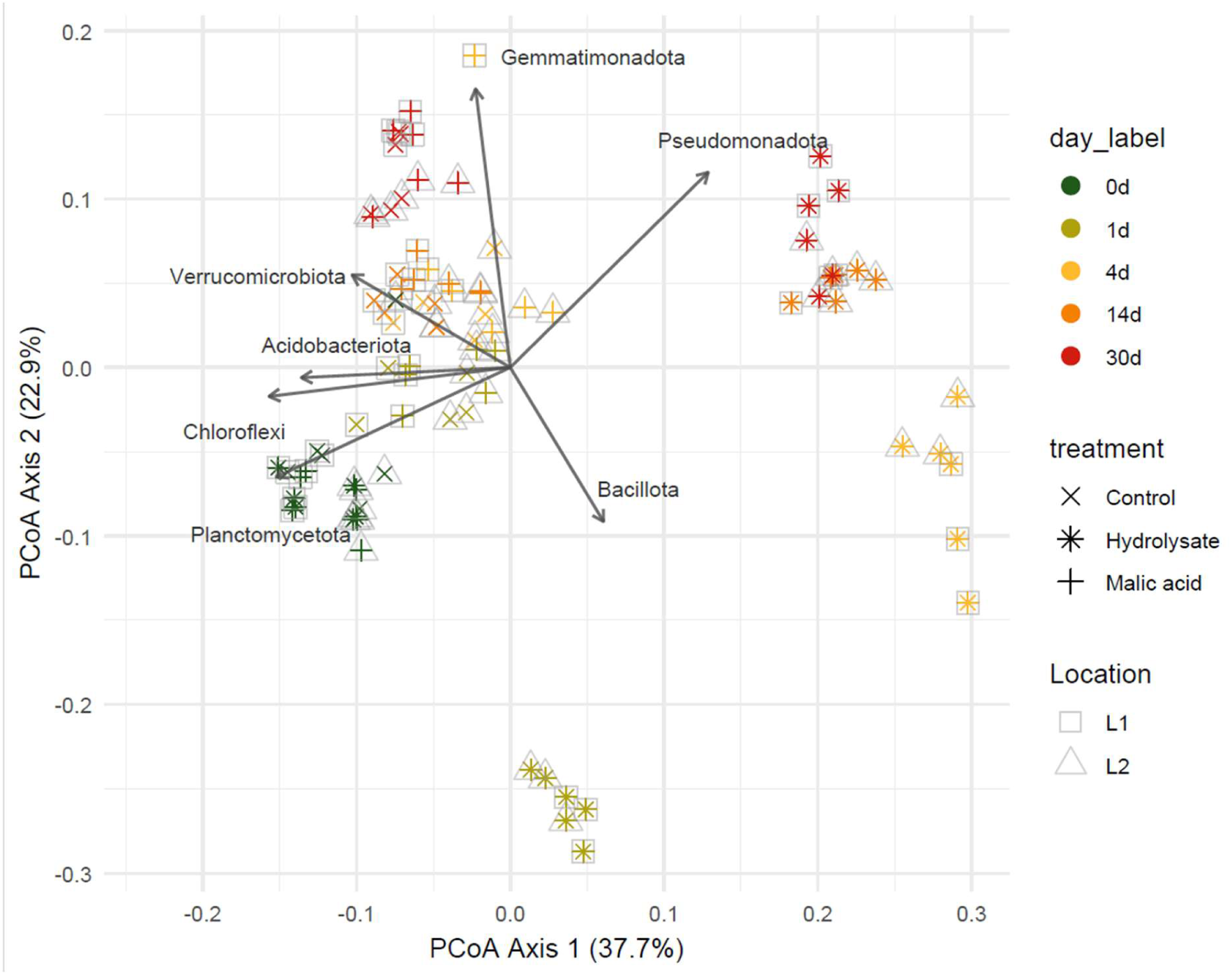
Prokaroytic beta-diversity of the soil samples, illustrating temporal dynamics following hydrolysate amendment.

A decrease in alpha diversity has been well documented following the addition of protein-or amino acid-rich materials (Caballero et al., 2020; Gu et al., 2020). In agreement with extensive literature, we hypothesize that this is driven by the rapid proliferation of a limited number of dominant, likely chemoheterotrophic and copiotrophic taxa, a phenomenon frequently observed after the addition of labile organic carbon (C_org_) (Costa et al., 2024; Fierer et al., 2007; Luo et al., 2023; Zegeye et al., 2019). This interpretation is supported by the shift in relative abundance of copiotrophic and oligotrophic bacteria, as assessed by the predicted *rrn* copy number of the detected genera (Fig. 3**Chyba****! Nenalezen zdroj odkazů.**, Fig. S3) (Bei et al., 2025). Specifically, the initial surge in copiotrophs peaking on day 1, followed by a subsequent increase in oligotrophs abundance by day 14, aligns closely with microbial succession theory (Figure S3) (Fierer et al., 2007).

**Figure 3.**
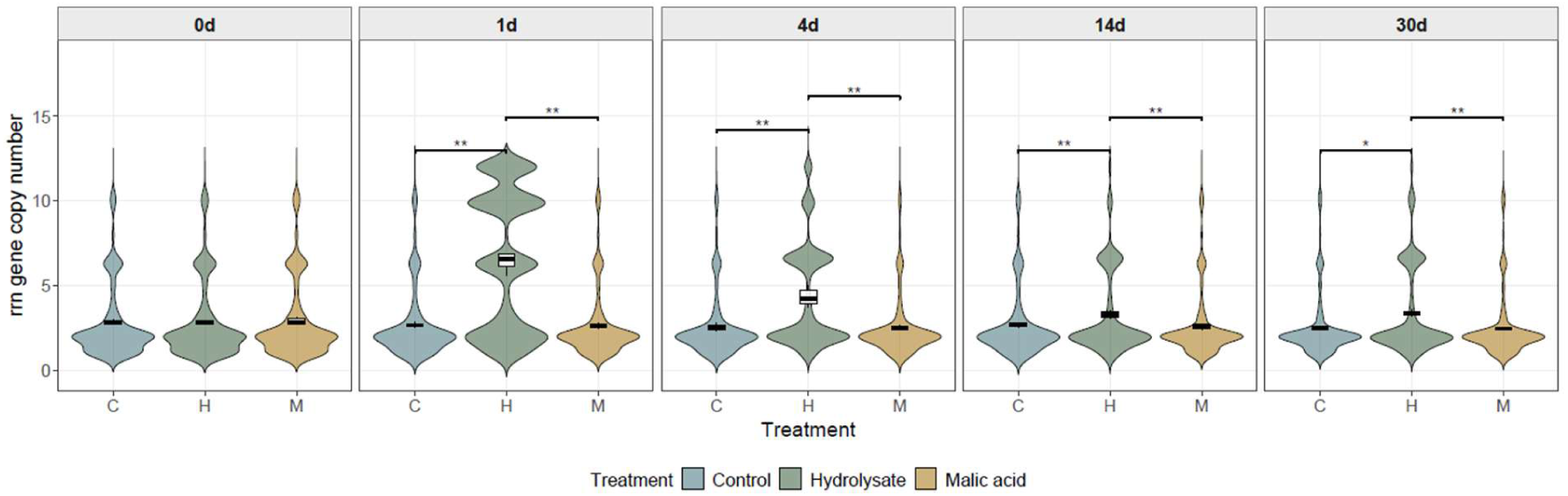
Temporal dynamics of predicted rrn gene copy number abundance. Genera with predicted low rrn copy number are generally classified as oligotrophic, and genera with higher rrn copy number as copiotrophic (Bei et al., 2025). Statistical significance (indicated by an asterisk, p < 0.05) is based on two-sided van Elteren test followed by Benjamini–Hochberg correction.

A methodological consideration of this study concerns the integration of taxonomic profiles derived from five independent amplicon datasets (see section 2.4, Soil microbiome analyses). To obtain a unified community composition estimate, we averaged relative abundances across the relevant amplicons for each taxonomic feature. We are aware that combining multiple amplicons is a less common approach and carries an inherent risk of obscuring otherwise clear signals from an individual marker gene, given that each marker differs in taxonomic resolution, amplification bias, and database coverage. However, since features absent from a given amplicon dataset were excluded rather than treated as zero, and prokaryotic and eukaryotic profiles were merged separately prior to integration, we believe that this conservative approach may in fact be beneficial: it can broaden the range of detected taxa and help smooth out biases specific to individual amplicons (Yue et al., 2025). In addition, we tested each amplicon dataset separately and observed broadly comparable results (Supplementary Data 2, Figures S15-S34). We acknowledge that this merging strategy remains non-standard, and future studies may benefit from analyzing each amplicon dataset independently or employing marker-gene-independent approaches such as shotgun metagenomics.

### 3.2. Major taxonomic shifts

Consistent with the observed decline in diversity, we found that among the nine most abundant prokaryotic phyla, five significantly decreased in abundance at multiple time points, while only two phyla increased (Fig. 4). The increased abundance of Pseudomonadota (formerly Proteobacteria) and Bacillota (formerly Firmicutes) in response to N enrichment aligns with previous findings (Ling et al., 2017). Similarly, the decline in Acidobacteria upon C_org_ input, as well as reductions in Acidobacteria, Chloroflexi, and Planctomycetes upon nitrogen (N) input, is well-documented (Fierer et al., 2007; Ling et al., 2017). More specifically, the decrease in Acidobacteria abundance observed here mirrors responses reported following glycine amendment to soil (Goldfarb et al., 2011).

**Figure 4.**
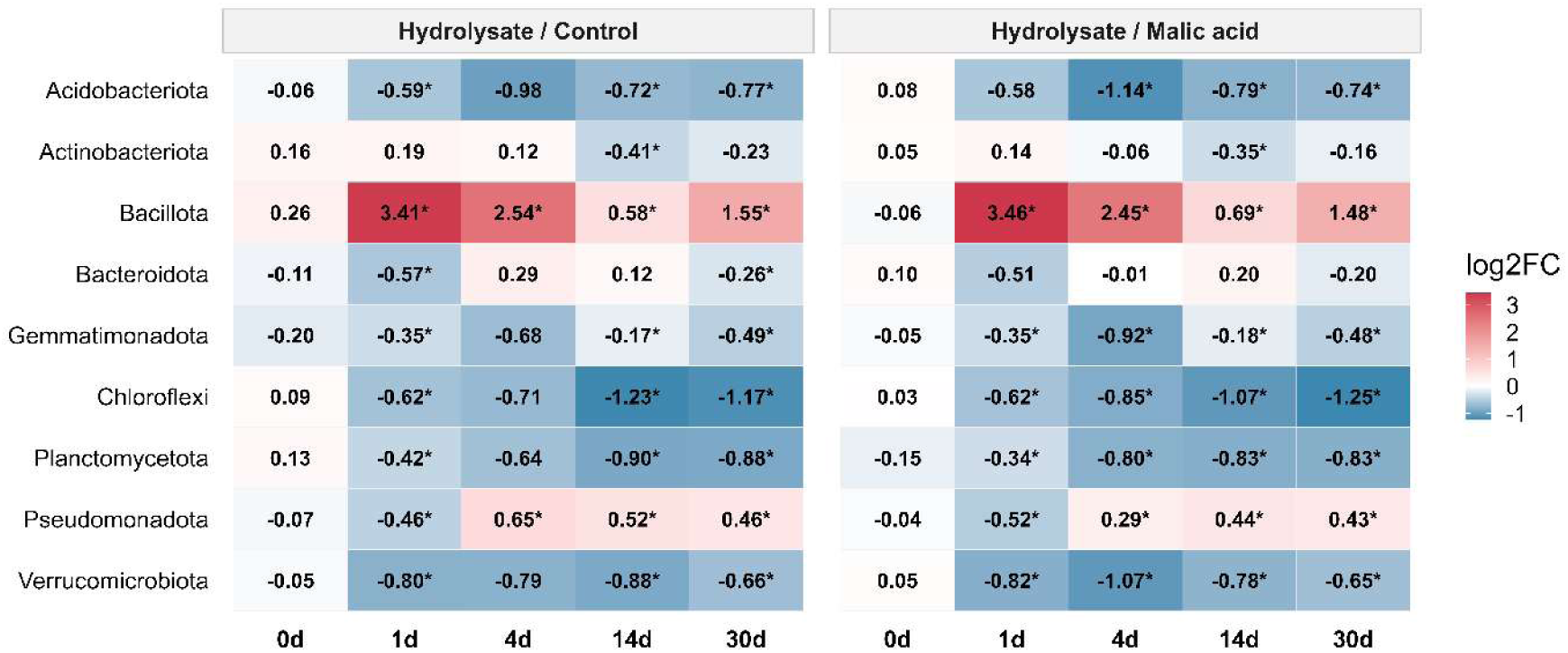
Temporal dynamics of the most abundant prokaryotic phyla.

**Figure 5.**
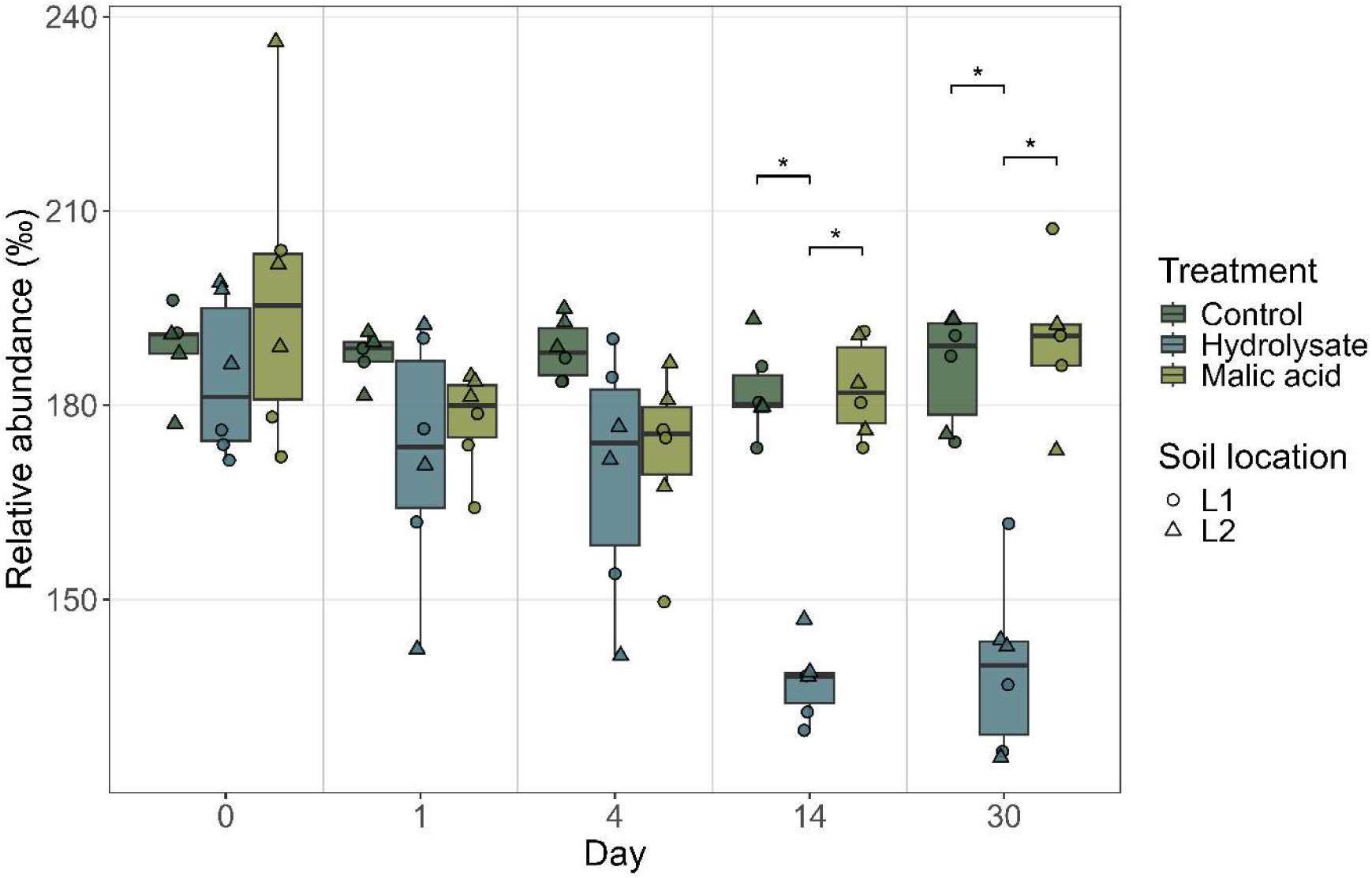
Decrease in the relative abundance of phytopathogenic genera based on UNITE predictions.

Regarding eukaryotic shifts, three clades increased in abundance at multiple time points while four clades decreased (Fig. S4). In particular, Mucoromycota are known to thrive in the presence of labile organic substrates (Pawłowska et al., 2019), similar to certain Basidiomycota that respond positively to organic nitrogen (Abuzinadah and Read, 1988; Tibbett et al., 2000). Interestingly, Gracilipodida increased in abundance later, starting from day 4, potentially reflecting the delayed response of bacterivorous protists to the preceding proliferation of bacterial populations.

At the genus level, the early responders, i.e. those showing increased relative abundance as early as day 1, were predominantly chemoheterotrophic bacteria (e.g. *Bacillus* and *Massilia*; Figure S5). Among eukaryotes, several saprotrophic fungi (such as *Mortierella* and *Derxomyces)* exhibited similar early-response patterns. Genera *Bacillus, Massilia*, and *Mortierella* have been previously identified as enriched in keratine-rich environments (Andreo-Jimenez et al., 2021; Gu et al., 2016; Russ et al., 2024). Furthermore, functional annotation using the UNITE database revealed that the group of genera significantly enriched by hydrolysate amendment contained a significantly higher proportion of soil saprotrophs, compared to the groups that either decreased or remained unaffected (Fig. S6).

Subsequently, the genera that initiated their proliferation at later stages (from day 4 or 14) were dominated either by chemoheterorophs (e.g. *Pigmentiphaga, Bosea*, and *Flavisolibacter*) or *Amoebozoa* (e.g. *Flamella*, *Acanthamoeba*, and *Heliamoeba*, Fig. S5B). In addition to amoebae, bacterial predators belonging to the family *Bacteriovoraceae* increased their abundance (Tab. S10), consistent with the findings of Russ et al. (Russ et al., 2024). Conversely, the genera that decreased in abundance following hydrolysate amendment were primarily associated with specialized microbial predation (e.g. *Bdellovibrio*, *Haliangium*, and *Arboramoeba*) (Adl et al., 2019; Lovering and Sockett, 2021; Petters et al., 2021; Whitworth et al., 2021), an oligotrophic life strategy (e.g. *Chthonomonas*, *Pirellula, and Phreatobacter*) (Fuerst, 1995; Dunfield et al., 2012; Tóth et al., 2014; Lee et al., 2016, 2017) or a phototrophic lifestyle (e.g. *Rhodomonas* and *Dictyococcus*) (da Silva et al., 2009; Di Accadia et al., 1966). Notably, many plant-associated fungi were also identified in this declining category, including both potential endophytes and phytopathogens.

To distinguish between the response of the indigenous soil microbiota and the potential introduction of exogenous microorganisms, we characterized the microbiome of the hydrolysate itself using Oxford Nanopore sequencing. Although the hydrolysate preparation includes sterilization-level processing, low-level secondary colonization was detected post-production. Approximately 20 genera were consistently detected in at least two independent hydrolysate samples, most belonging to the *Pseudomonadota* phylum (e.g. *Escherichia*, *Sphingobium,* etc.), however, the abundance of most of these genera in the soil samples either remained negligible or did not show any marked increase in the hydrolysate-amended soil (Tab. S11 and S12). Three of the genera that significantly increased in abundance during the experiment, *Rhodococcus, Pseudoxanthomonas* and *Pseudomonas*, were also detected in the hydrolysate itself. Although all these bacteria are ubiquitous soil inhabitants, we cannot rule out an inoculation effect in this case.

### 3.3. Hydrolysate-driven suppression of phytopathogenic load

Predictions of primary ecological niches using the UNITE database indicated a substantial decrease in the total abundance of potentially phytopathogenic genera (**Chyba! Nenalezen zdroj odkazů.**5, **Chyba! Nenalezen zdroj odkazů.**8). While the reduction in cumulative phytopathogen abundance became significant starting from day 14, several dominant genera declined as early as day 1 (Fig. 6). Conversely, among 16 most abundant genera, five exhibited mild abundance increase, including *Fusarium* at day 4 and *Alternaria* at later experiment stages. These results were consistent across both soil samples (Fig. S9 and S10).

**Figure 6.**
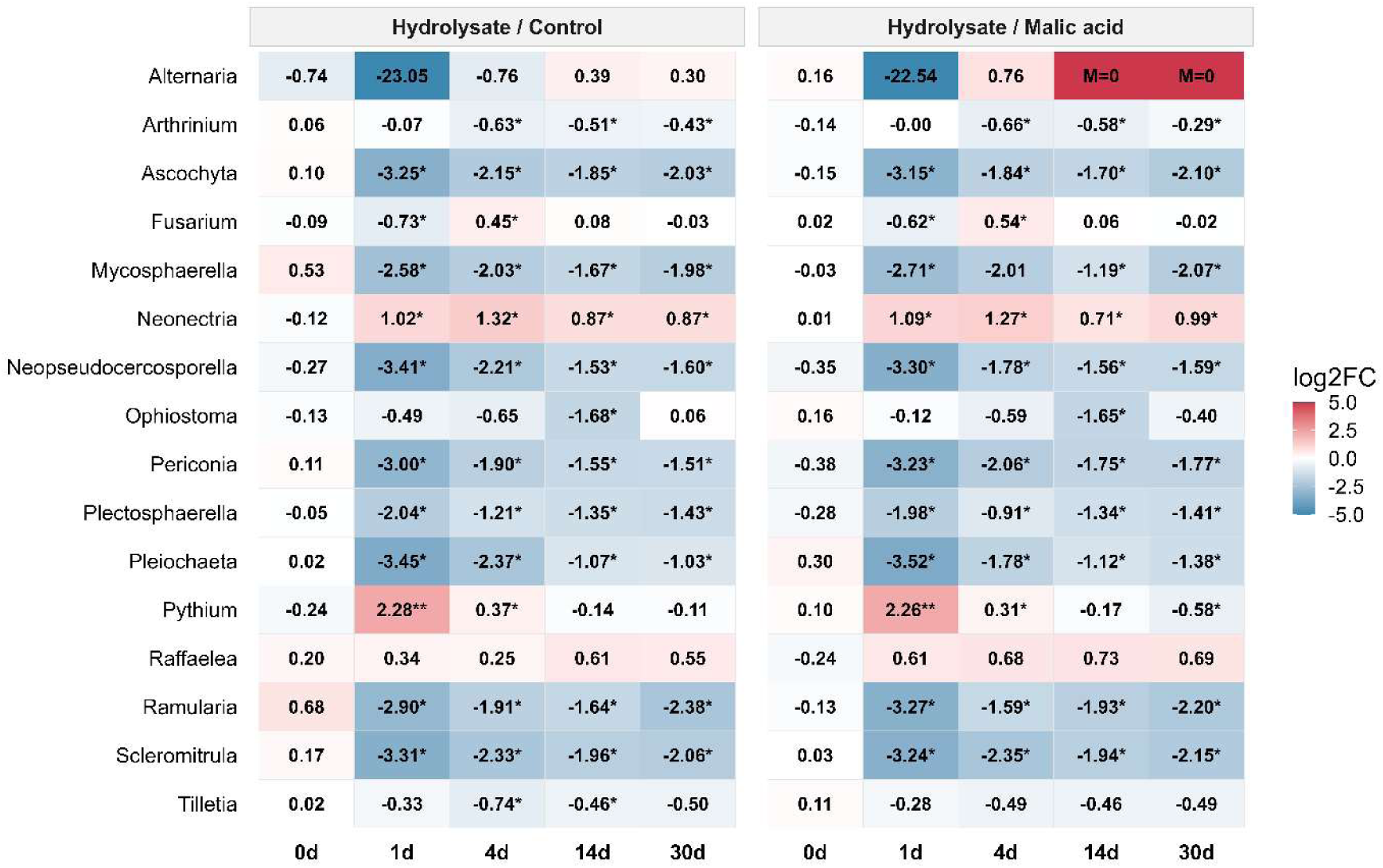
Temporal dynamics of the 16 most abundant phytopathogenic genera, shown as log-fold changes in hydrolysate-amended samples relative to controls (water and pH-adjusted malic acid solution), both soil samples (L1 and L2) are pooled.

In line with these results, PHs are expected to be beneficial for their pathogen suppression by increasing microbial biomass, diversity or overall soil suppressiveness (Colla et al., 2017). Such protective effects have been documented for various keratin-rich materials; for instance, human hair hydrolysate was shown to increase the survival of hot pepper plants infected with *Ralstonia solanacearum* (Oh et al., 2011), while non-hydrolyzed keratin reduced disease symptoms of *Rhizoctonia solani* (Andreo-Jimenez et al., 2021). Our findings also align with the role of specific amino acids in these processes, as previously demonstrated for *Botrytis* and *Fusarium* suppression (Kim et al., 2021; Papazlatani et al., 2025). On the other hand, even contrasting effects of PH amendments on plant infections have been reported, as well as poor relation between pathogenic load and disease severity (Gu et al., 2020; Schmidt et al., 2020). Still, we believe that this successional data may provide a basis for identifying windows of enhanced suppressiveness against particular pathogens, which could inform the optimal timing of FH application relative to plant growth stages. However, fully leveraging this potential would require further integration with direct observations of disease incidence.

### 3.4. Hydrolysate-driven microbial shifts in plant growth-promoting functional niches

Interestingly, several taxa that maintained elevated abundance throughout the experimental period, such as *Bacillus*, *Burkholderia,* and *Pseudomonas*, are widely recognized for their plant growth-promoting capacity (Fig S4A) (Saeed et al., 2021; Upadhyay et al., 2022; Vejan et al., 2016). Most PGPR are heterotrophs that thrive on labile carbon sources originating from root exudates; in this study, such conditions were likely mimicked by the pulse of amino acids (Upadhyay et al., 2022). To investigate these functional shifts in greater detail, we compiled a list of typical PGPR genera and their functional subgroups from the literature (Tab. S7). While total inferred PGPR abundance showed a significant increase in hydrolysate-amended soil only on day 1, a more granular analysis revealed that multiple specific functional potentials increased significantly at multiple time points (Fig. 7, S11 and S12).

**Figure 7.**
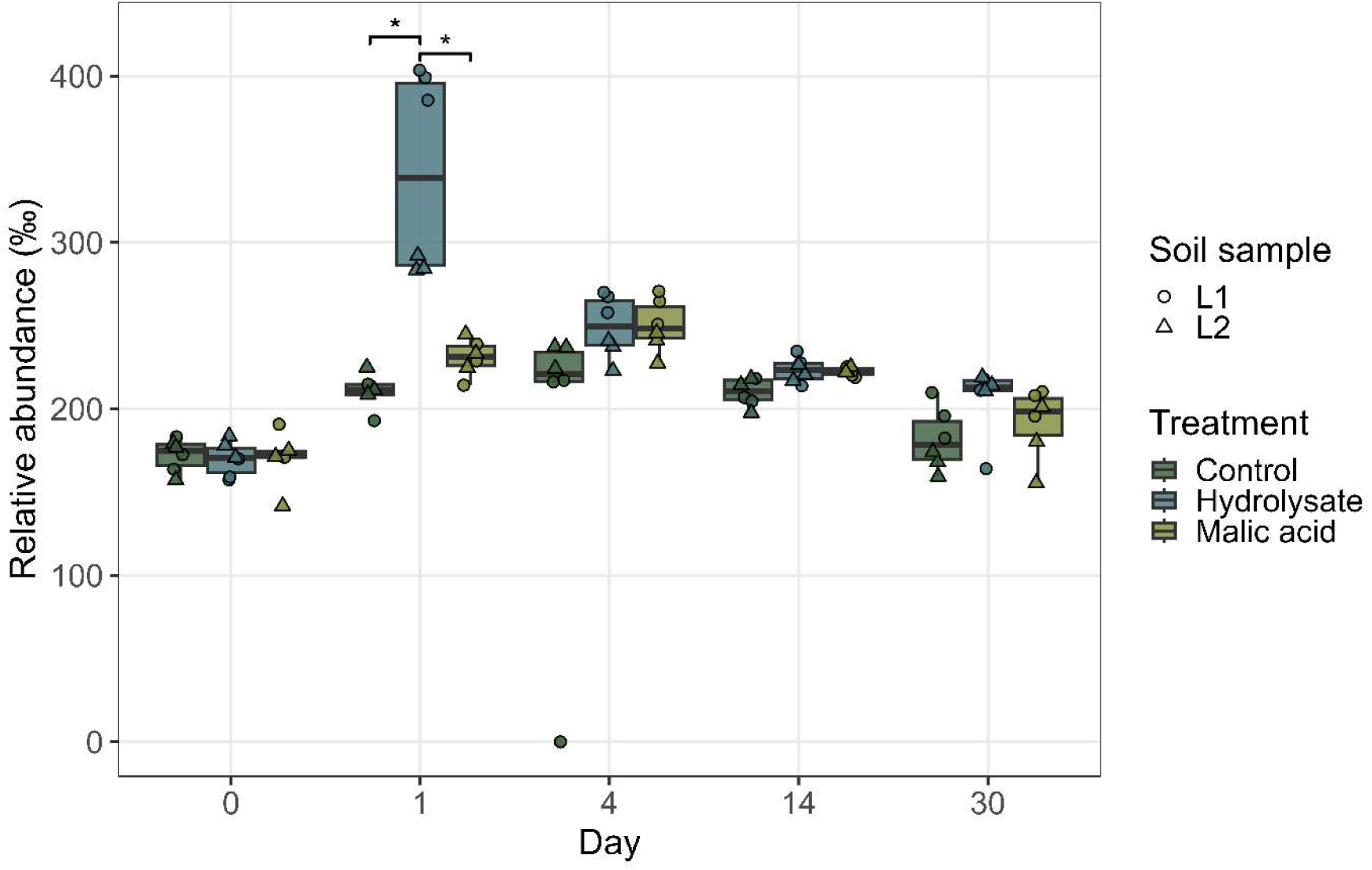
Temporal dynamics of the relative abundance of bacterial taxa classified as PGPR.

In line with the decline in phytopathogenic genera, we observed a 300% increase in the cumulative abundance of potential pathogen suppressors on day 1, with elevated levels persisting through day 30 (Fig. S11). This recruitment of Sphingobacteriaceae, Oxalobacteraceae, Comamonadaceae, and *Lysobacter* (Tab. S10) aligns with previous reports on keratin-rich amendments (Caballero et al., 2020; Russ et al., 2024). Furthermore, the enrichment of Burkholderiaceae, Pseudomonadaceae, and Mortierella directly supports the observed biocontrol potential (Andreo-Jimenez et al., 2021). Notably, the expansion of *Streptomyces*, Bacillaceae, and Burkholderiaceae observed in our study (Tab. S10) mirrors the effects of glutamic acid treatment, which has been shown to suppress *Botrytis*- and *Fusarium*-induced diseases by modulating these specific taxa (Kim et al., 2021).

The observed pathogen suppression likely involves both direct and indirect mechanisms (Philippot et al., 2013). Beyond the potential production of antibiotics (Russ et al., 2024) and antimicrobial VOCs (Papazlatani et al., 2025), our data suggest that indirect suppression via iron sequestration may play a significant role, as evidenced by the elevated abundance of putative siderophore-producing genera (Fig. S11) (Olanrewaju et al., 2017).

Similar to pathogen-suppressing taxa, we observed significant increases in genera associated with P, K, and Zn solubilization, as well as putative phytohormone and ACC-deaminase producers (Fig. S11). The enrichment of putative P and K solubilizers following feather-based amendments aligns with previous reports (Jagadeesan et al., 2023; Paul et al., 2013), as does the proliferation of specific phytohormone-producing genera such as *Pseudomonas*, *Paraburkholderia*, and *Streptomyces* (Costa et al., 2024; Kim et al., 2021). Notably, while previous studies have focused on individual taxa, our approach offers a more holistic assessment by evaluating phytohormone production potential as a comprehensive functional group.

Interestingly, several authors showed auxin-like effects of PHs on coleoptile and root explants, even under limitation of rhizobial bacteria (Buffagni et al., 2021; Colla et al., 2014; Lucini et al., 2020; Massaya et al., 2022; Ugolini et al., 2015). Therefore, it appears that PHs can exert phytohormone-like activity on plants both directly and indirectly through the stimulation of microbial phytohormone production. The relative contribution of each pathway remains a subject for future research.

Putative nitrogen fixers exhibited a transient decline followed by a significant increase relative to controls (Fig. S13). We believe that this initial shift likely reflects the rapid expansion of copiotrophic heterotrophs following the nutrient pulse rather than an absolute decrease in diazotrophic populations. This successional trend aligns with the initial dip and subsequent recovery observed by Paul et al. (Paul et al., 2013) and the late-stage enrichment of nitrogen fixers reported by Jagadeesan et al. (Jagadeesan et al., 2023). Analysis of genera classified as potential nitrifiers and denitrifiers showed a peak on day 1, with abundance remaining significantly higher than controls through day 30 (Fig. S13). This pattern suggests a gradual depletion of available nitrogen in the hydrolysate-amended soil over time. Additionally, our analysis showed a gradual decrease in active carbon-indicating genera, suggesting a reduction in available C_org_ within the system from day 1, likely due to rapid microbial sequestration (Fig. S14). These changes are concordant with the known half-life of amino acids in soil, which is approximately 3-6 hours in topsoil (Kemmitt et al., 2008; Kielland et al., 2007).

The observed overlap in successional patterns across various PGPR functional groups is consistent with the well-documented multifunctionality of these taxa (Rosier et al., 2018). For instance, genera with the potential to solubilize mineral elements (P, K, Zn, and Fe) frequently overlap, and many phytohormone-producing genera are capable of synthesizing multiple growth-regulating substances (Tab. S7). While the herein presented findings are derived from broad taxonomic resolution and represent functional potential, essentially a probabilistic assessment, their consistency and strong alignment with existing literature suggest they accurately reflect the successional dynamics of soil microbial potential following FH amendment.

## 4. Conclusions

This study establishes a novel temporal framework for understanding soil microbiome successions following feather hydrolysate (FH) amendment. Our results demonstrate a sustained reduction in the majority of dominant phytopathogenic genera, occurring alongside a concurrent recruitment of potential pathogen suppressors and diverse PGPR groups, including inferred nutrient solubilizers and phytohormone producers.

These successional dynamics suggest the existence of specific temporal windows of phytopathogen suppression, which could be strategically aligned with critical plant phenological stages to optimize biostimulant efficacy. Furthermore, the observed shifts in carbon and nitrogen cycling indicators may reflect a transition from a nutrient pulse to a rather stabilized microbial state.

While our findings provide a consistent, literature-supported genomic perspective on FH-mediated restructuring, they represent functional potential rather than realized ecosystem services. Future research should bridge this gap by linking these genomic potentials to direct measurements of disease progression, nutrient fluxes, and plant performance. Such functional validation across diverse environments will be essential for the precise integration of circular biostimulants into sustainable global crop production.

## Supporting information

Table_S11_metagenome_of_hydrolysate

Table_S8_read_depths

Table_S9_BC_distances

Table_S10_significant_taxon_abundance_changes

Table_S12_all_taxa_changes

Tab_S6_PCR_primers_and_conditions

Tab_S7_list_of_literature_derived_taxa_functions

## 5. Acknowledgements

This work was supported by the from the Technology Agency of the Czech Republic (TA CR) and the National Center of Competence BIOCIRCL [project number TN02000044], the Institute of Biophysics of the Czech Academy of Sciences [internal grant number 888861], the project TowArds Next GENeration Crops of the European Regional Development Fund (ERDF), Johannes Amos Comenius programme [registration number CZ.02.01.01/00/22_008/0004581]. Computational resources were provided by the e-INFRA CZ project [ID:90254], supported by the Ministry of Education, Youth and Sports of the Czech Republic.

## 6. Data availability

Raw sequencing data have been deposited in Zenodo repository and are are openly available at https://doi.org/10.5281/zenodo.22710395 under a CC BY 4.0 licence.

## 7. Author contributions

**Jan Veselský:** Conceptualization, Methodology, Investigation, Formal analysis, Data curation, Writing – original draft

**Martina Dlasková:** Conceptualization, Methodology, Investigation, Writing – original draft

**Radim Čegan:** Formal analysis, Data curation

**Šárka Kobzová:** Methodology, Investigation

**Zdeněk Kubát:** Methodology, Investigation

**Lenka Vaňková:** Methodology, Investigation

**Roman Hobza:** Conceptualization, Funding acquisition, Project administration

**Olga Šolcová:** Conceptualization, Funding acquisition, Project administration

**Lucie Kajan Grodecká:** Conceptualization, Methodology, Investigation, Formal analysis, Data curation, Writing – original draft, Funding acquisition

**All authors:** Writing – review and editing

## 8. Declaration of generative AI and AI-assisted technologies in the manuscript preparation process

During the preparation of this work the authors used ChatGPT, Gemini and Claude in order to improve language, structure and clarity of the text, and to help with figures (R-scripts) preparation. After using these tools, the authors reviewed and edited the content as needed and take full responsibility for the content of the published article.

