## Supplementary_data1 for "Organic-acid-derived feather hydrolysate drives soil microbiome succession toward copiotrophic and putatively plant-beneficial taxa"

**Supplementary data 1**

***Organic-acid-derived feather hydrolysate drives soil microbiome succession toward copiotrophic and putatively plant-beneficial taxa* Veselský J. et al.**

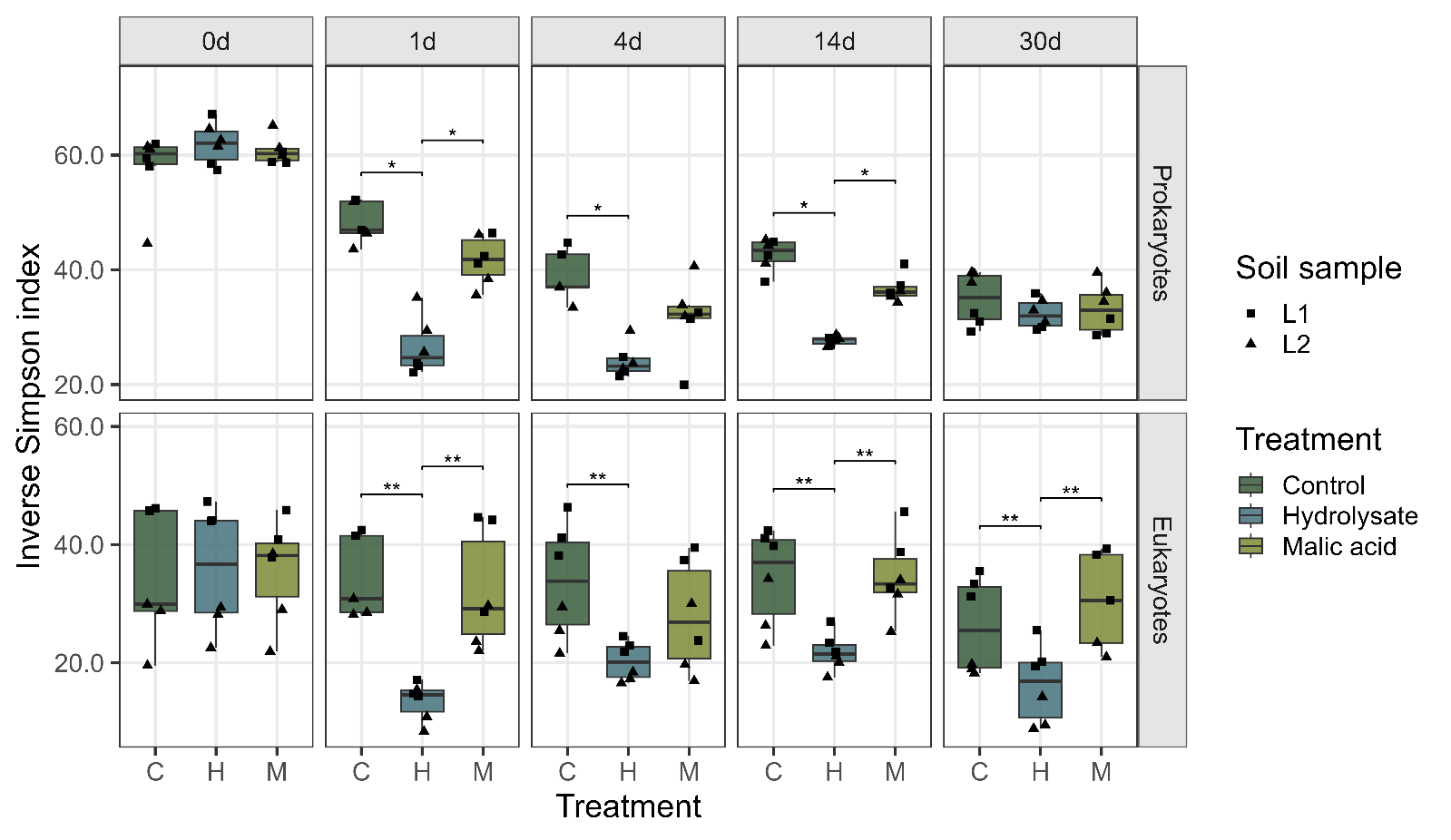

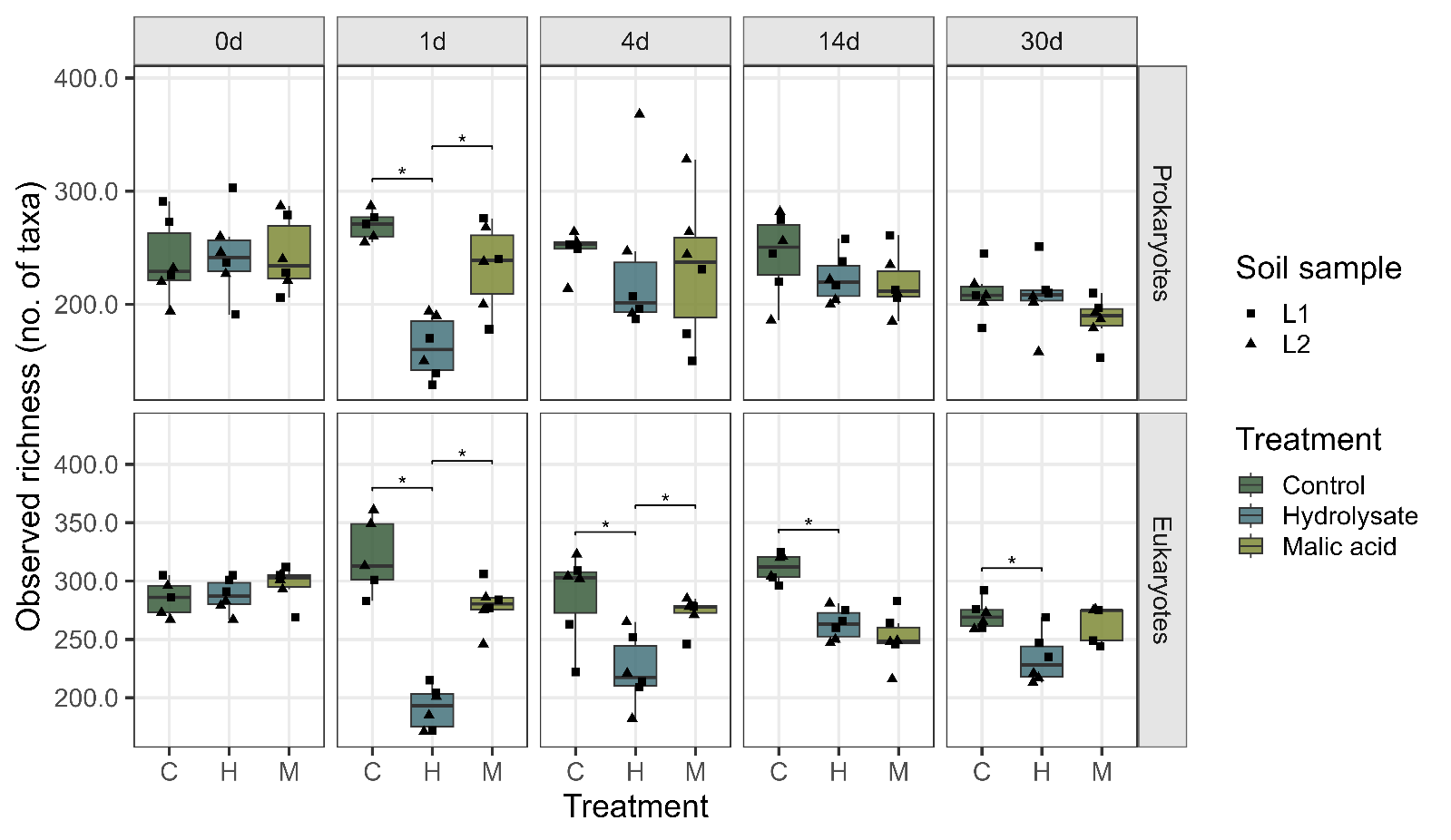

**Figure S1.** Alpha diversity of the soil microbial community 0–30 days after hydrolysate amendment. Boxes represent the interquartile range; whiskers indicate the range excluding outliers. Statistical significance (* p(adj.) < 0.05; ** p(adj.) < 0.01) is based on two-sided van Elteren test followed by Benjamini–Hochberg correction.

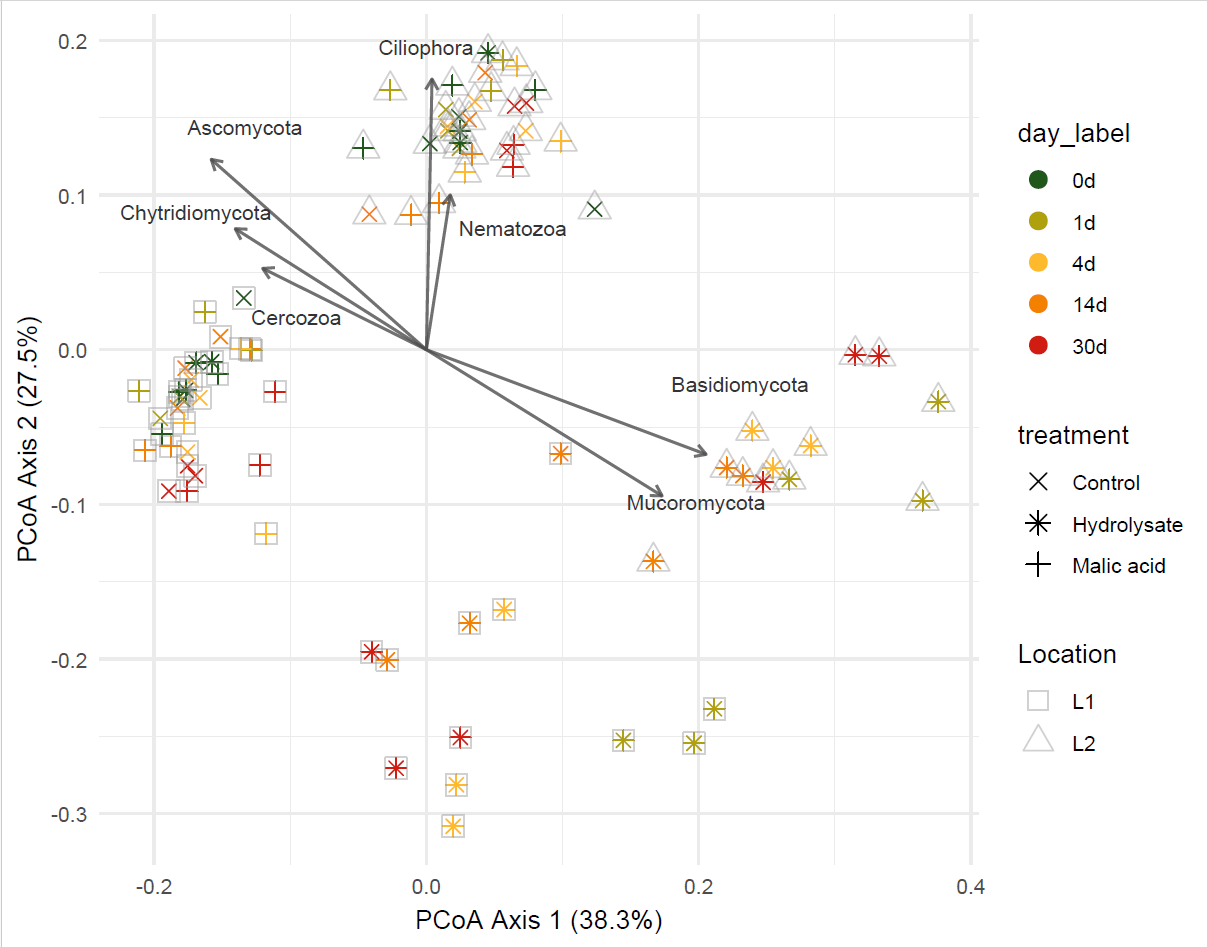

**Figure S2**. Eukaryotic beta-diversity of soil samples, illustrating temporal dynamics following hydrolysate amendment.

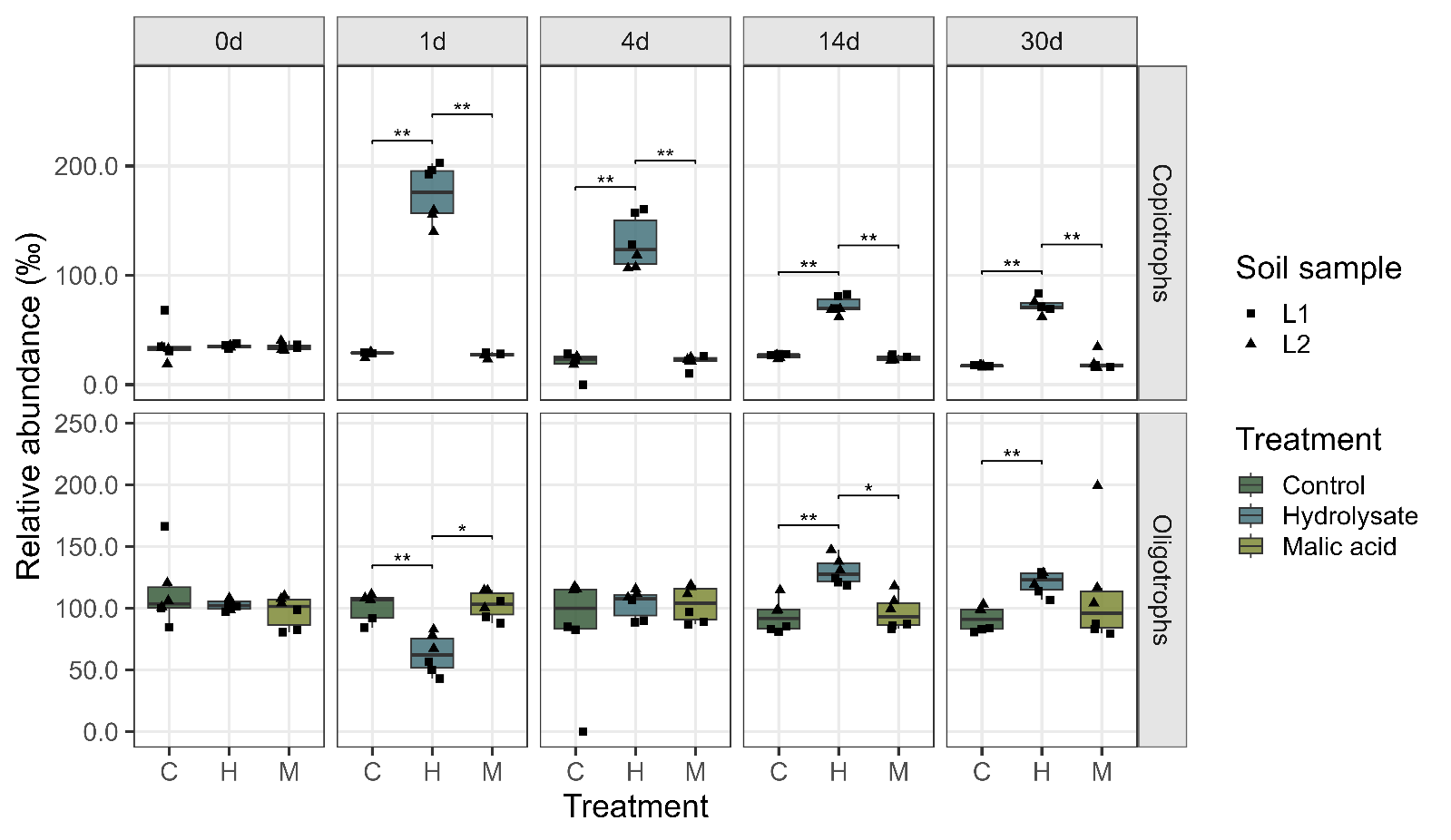

**Figure S3**. Temporal dynamics of copiotrophic and oligotrophic bacteria abundance based on predicted rrn copy numbers. Genera with a predicted rrn copy number ≤ 2 were classified as oligotrophic, and genera with rrn copy number of ≥ 5 as copiotrophic.

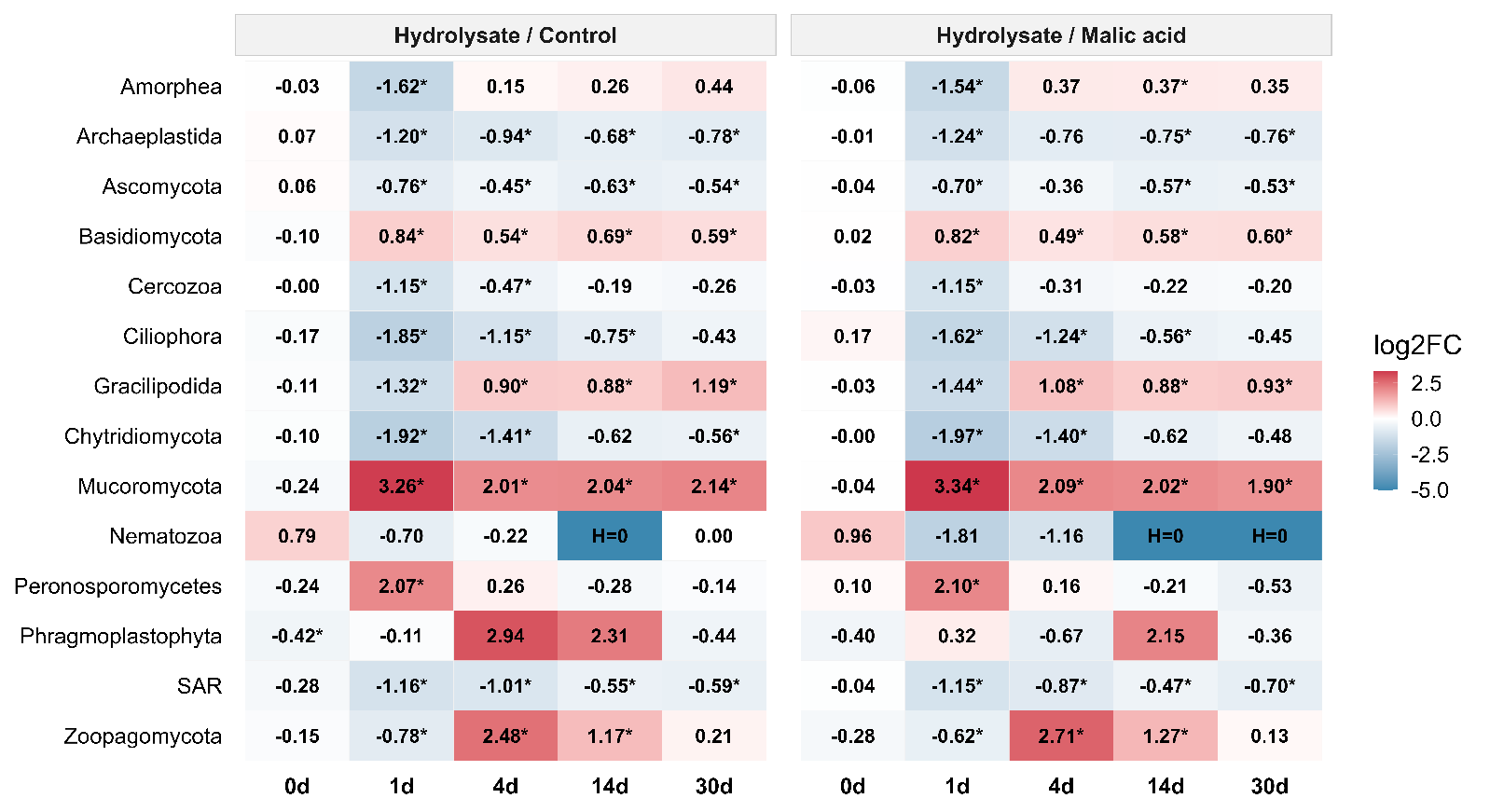

**Figure S4.** Temporal dynamics of the most abundant eukaryotic clades.

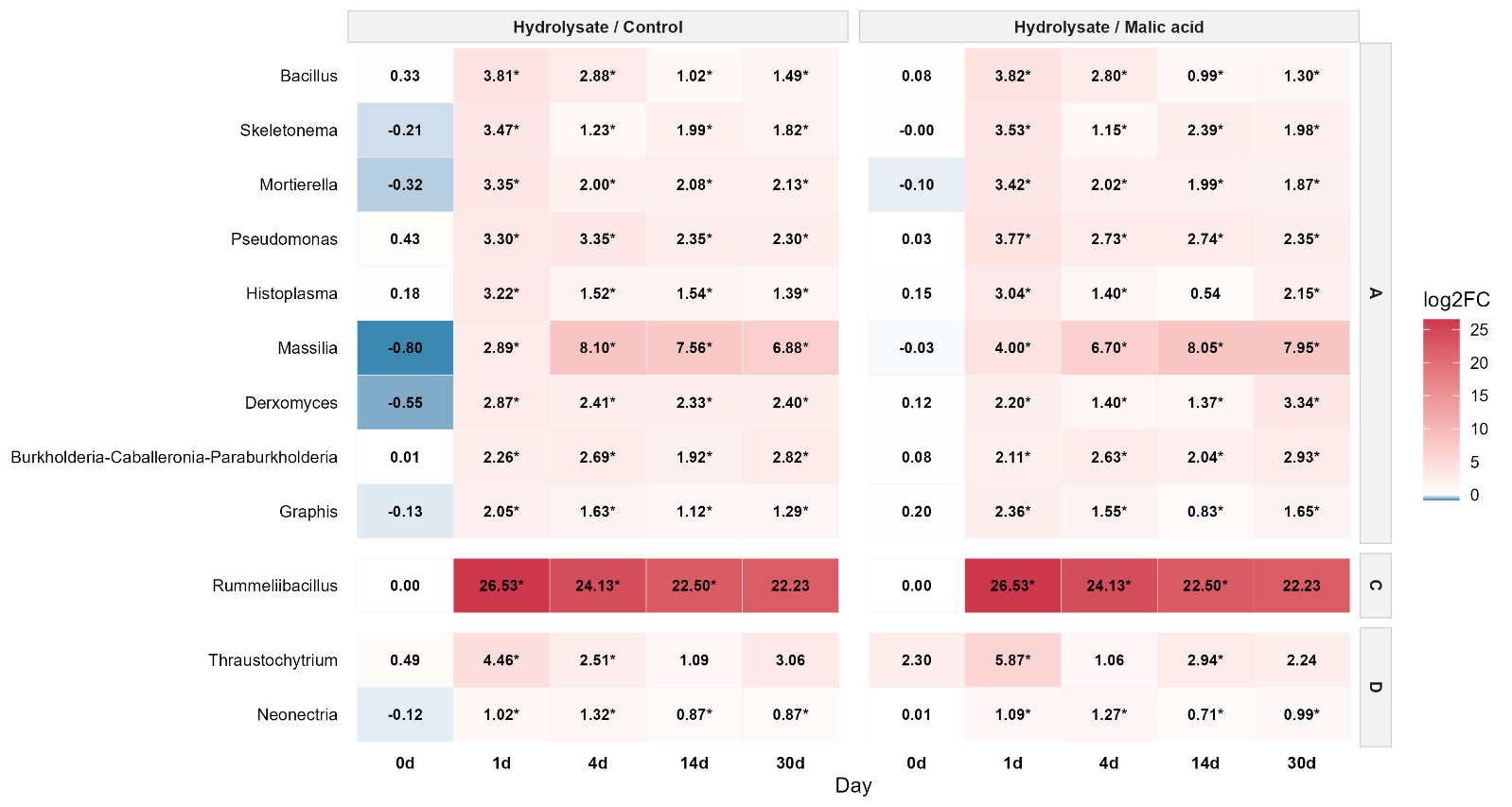

Early responders

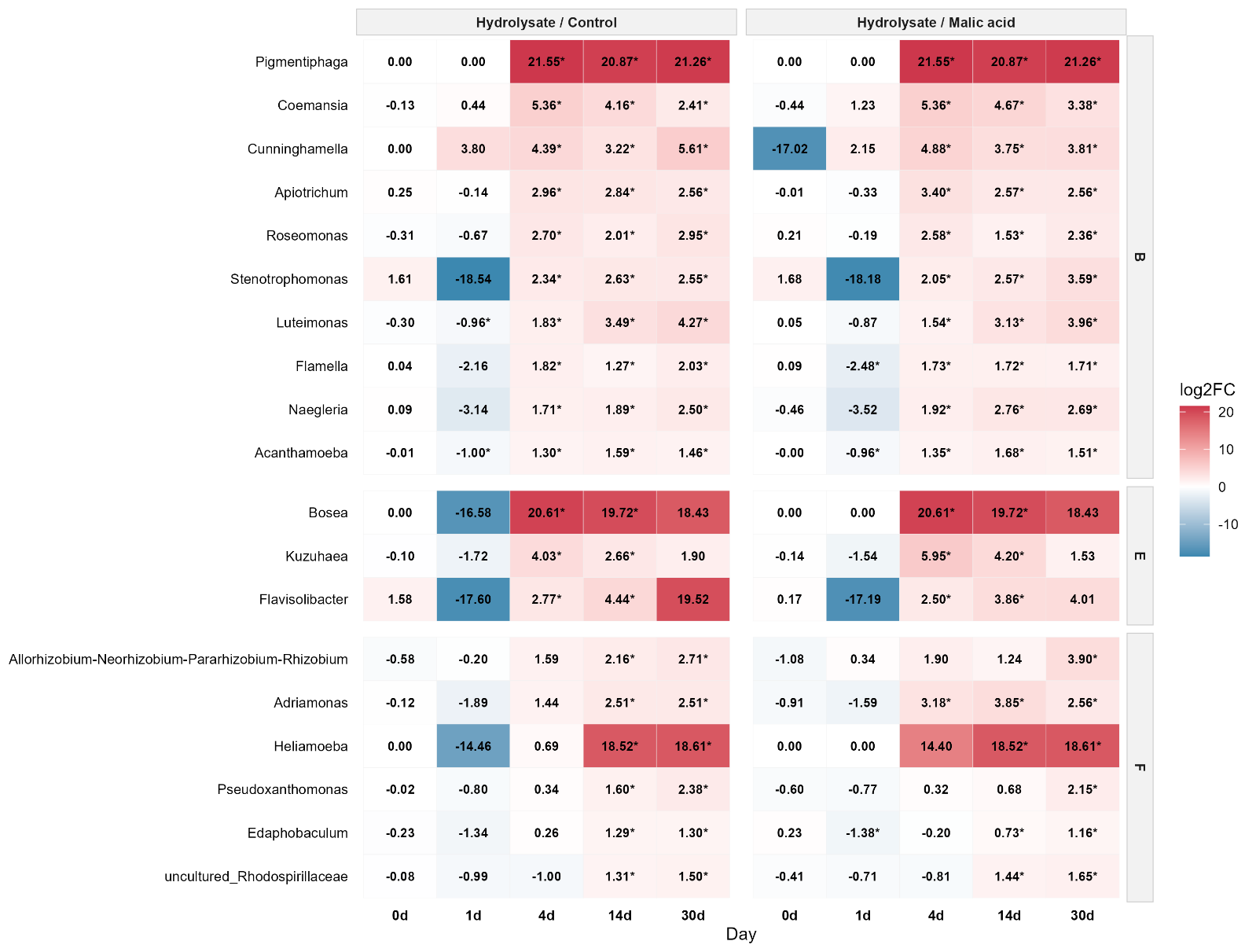

Late responders

**Figure S5. Positive responders**: genera showing increased abundance at early and late experimental stages. The groups A-F represent temporal response classification as follows: **Early Responders (A, C, D)** — genera whose response was already significant on day 1; **A** — significant at all four timepoints (1, 4, 14, and 30 d); sustained response; **C** — significant at 1, 4, and 14 d, but not at 30 d; response fades by the end of the experiment; **D** — significant only at 1 and 4 d; brief, transient early response. **Late Responders (B, E, F)** — genera not yet significant on day 1, with the response emerging later; **B** — significant at 4, 14, and 30 d (not at 1 d); sustained late response; **E** — significant only at 4 and 14 d; transient response, fading by day 30; **F** — significant only at 14 and 30 d; delayed response emerging only from day 14 onward.

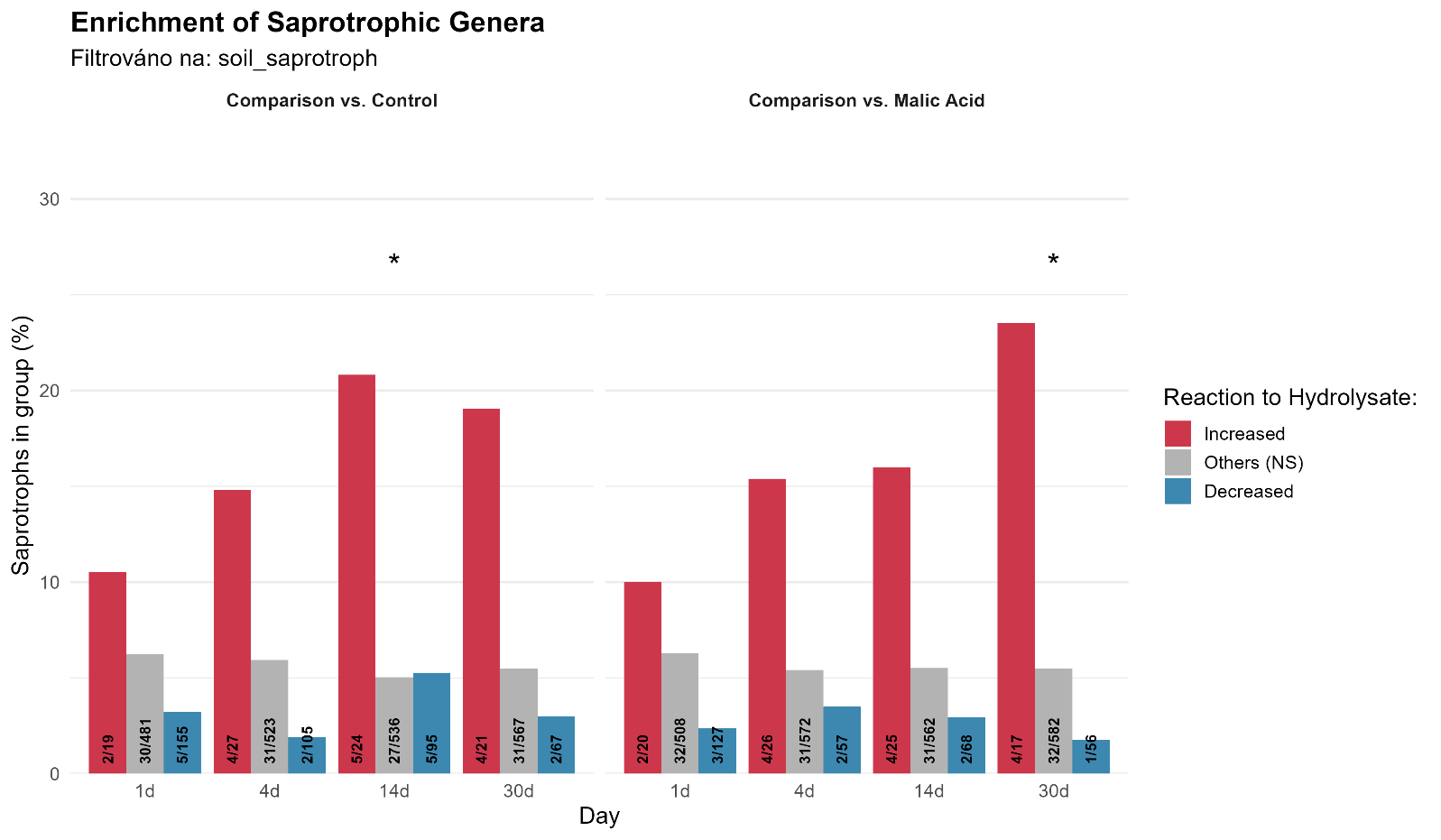

**Figure S6.** Enrichment of saprotrophic genera (UNITE-annotated as soil saprotrophs) among significantly increased genera at individual experiment days, statistical comparison with all other eukaryotic genera was provided using Fisher’s exact test, p(adj.) <0,05 is marked as *.

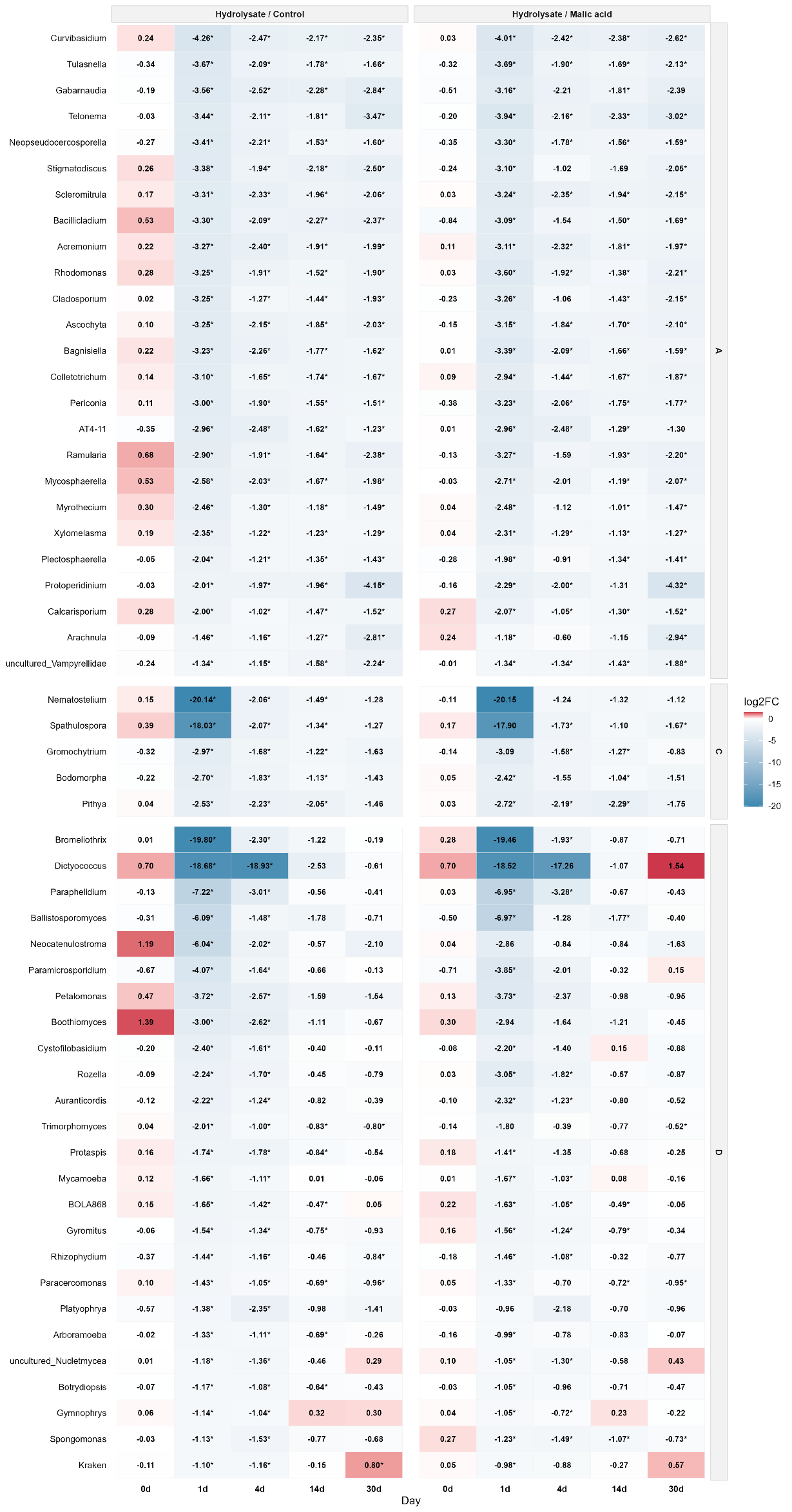

A

Early responders

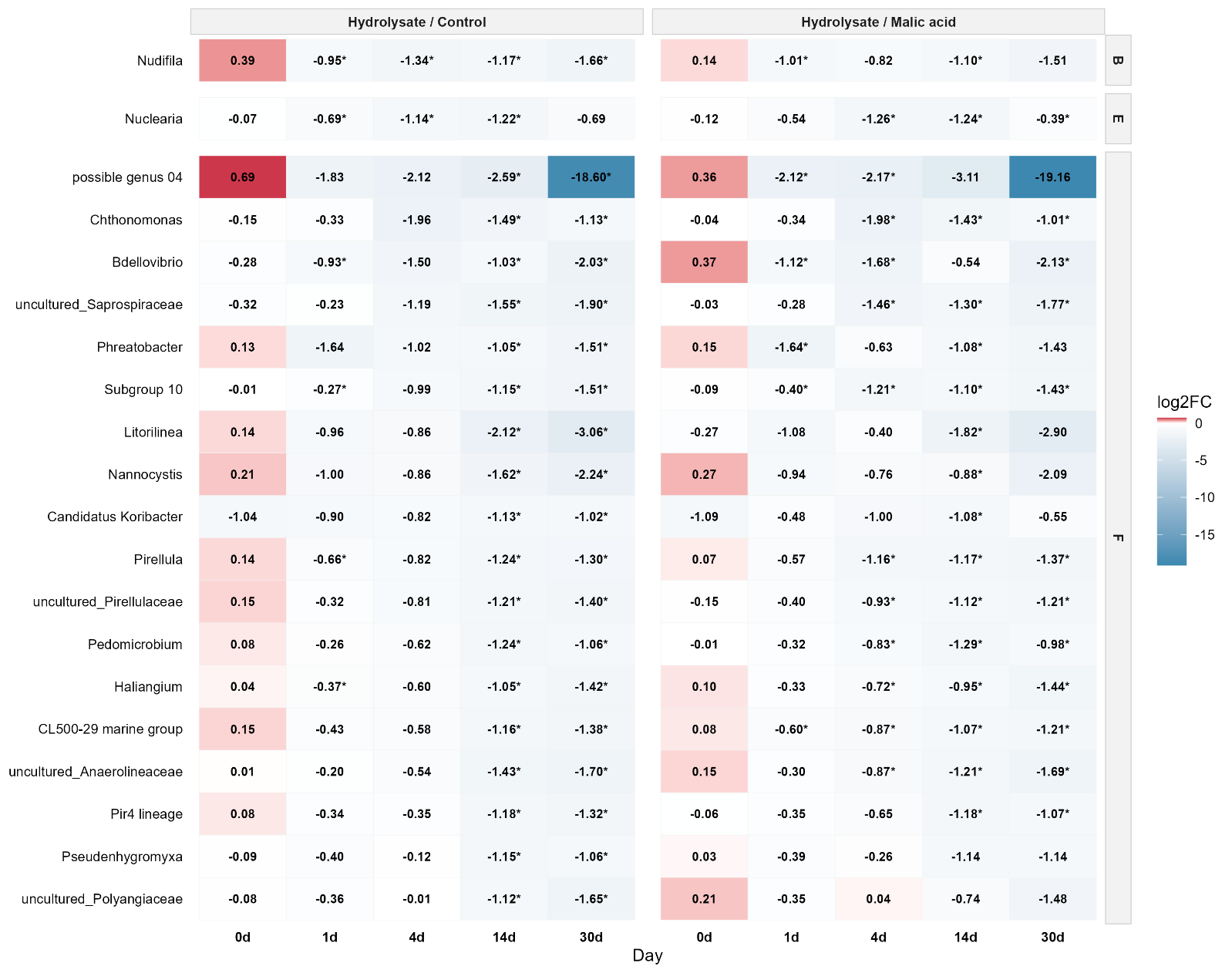

Late responders

**Figure S7.** Negative responders: genera showing decreased abundance at early and late experimental stages. The groups A-F represent temporal response classification as follows: **Early Responders (A, C, D)** — genera whose response was already significant on day 1; **A** — significant at all four timepoints (1, 4, 14, and 30 d); sustained response; **C** — significant at 1, 4, and 14 d, but not at 30 d; response fades by the end of the experiment; **D** — significant only at 1 and 4 d; brief, transient early response. **Late Responders (B, E, F)** — genera not yet significant on day 1, with the response emerging later; **B** — significant at 4, 14, and 30 d (not at 1 d); sustained late response; **E** — significant only at 4 and 14 d; transient response, fading by day 30; **F** — significant only at 14 and 30 d; delayed response emerging only from day 14 onward.

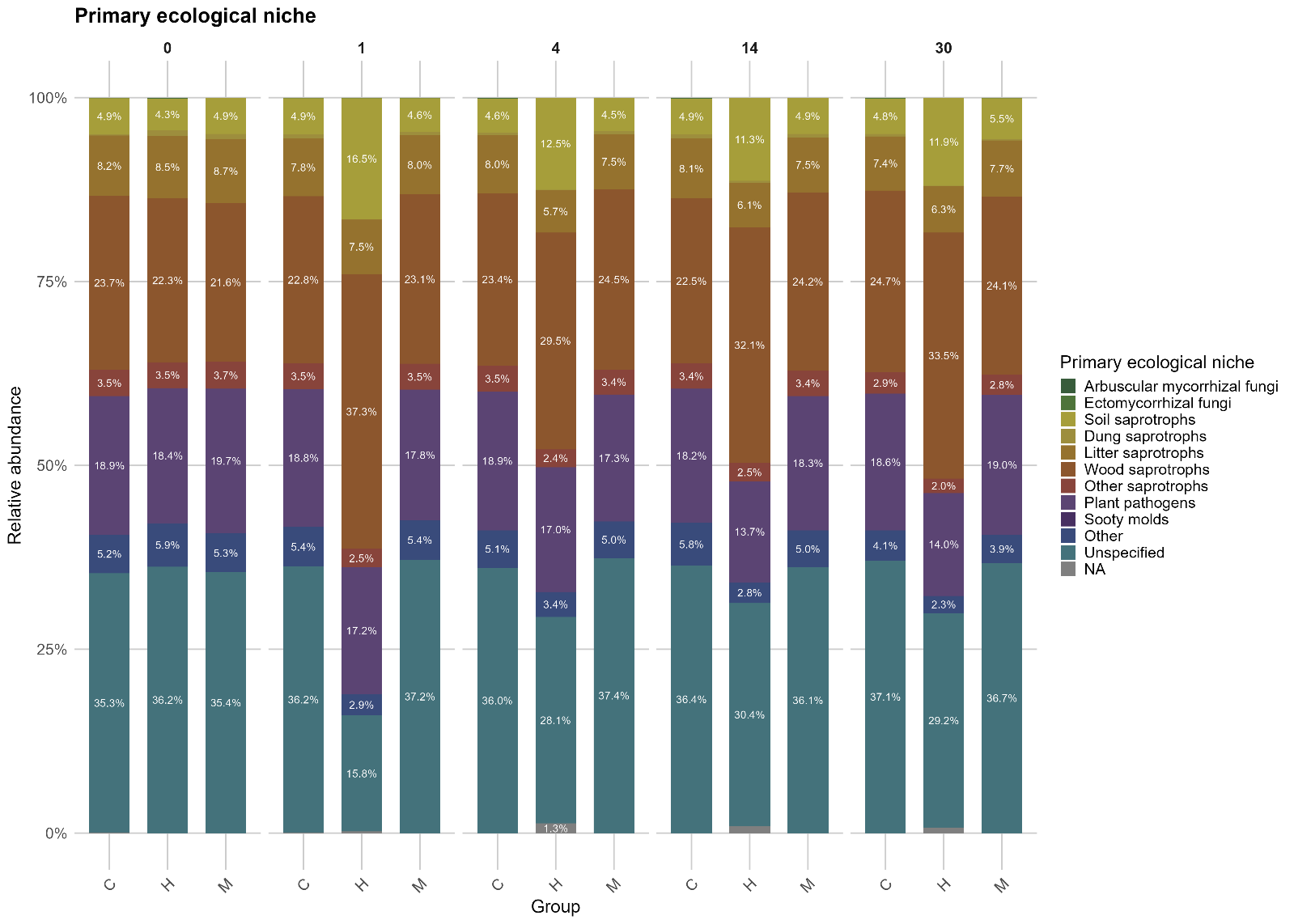

**Figure S8.** Primary ecological niches of eukaryotic microbiota according to UNITE database predictions.

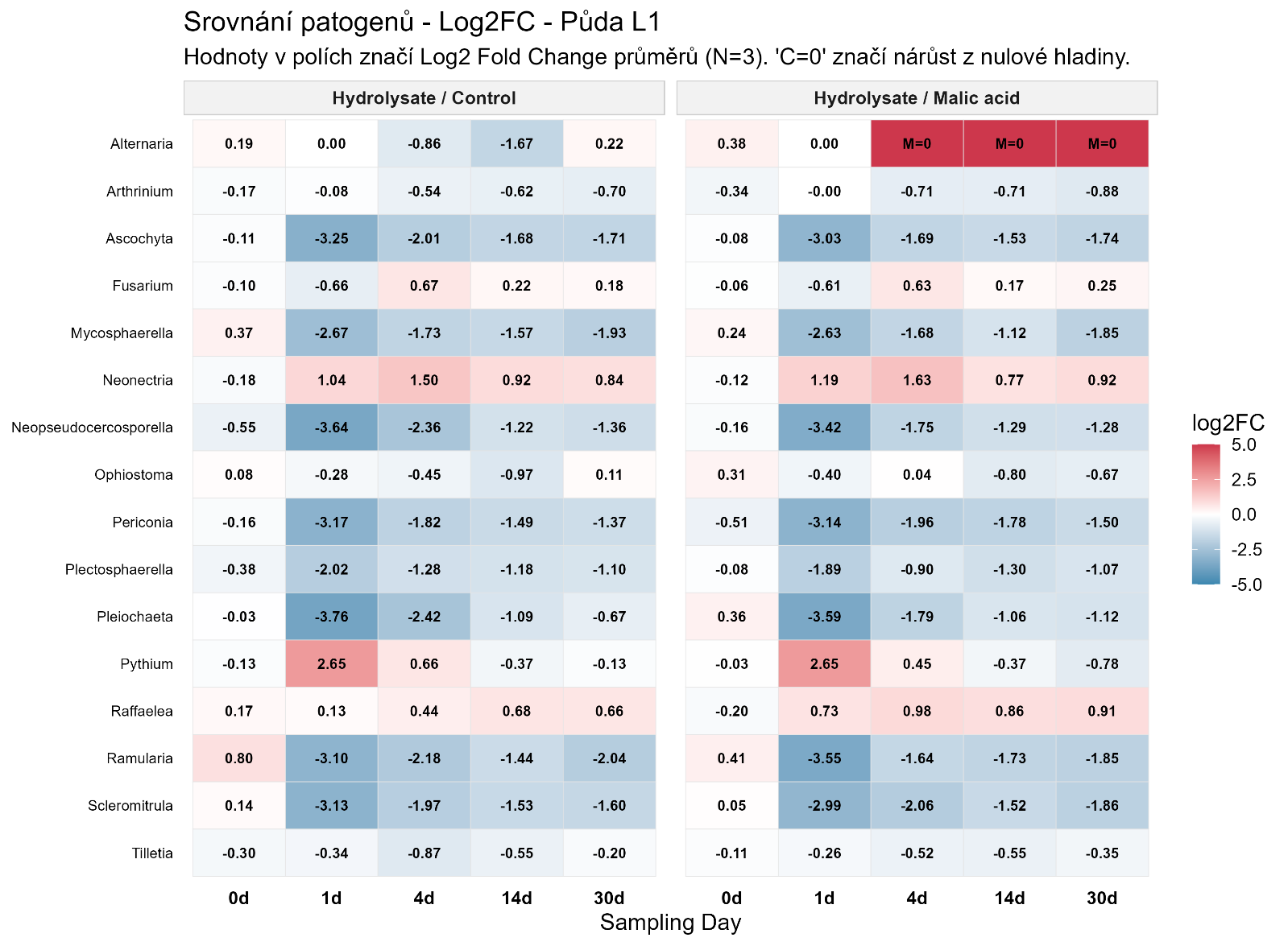

A

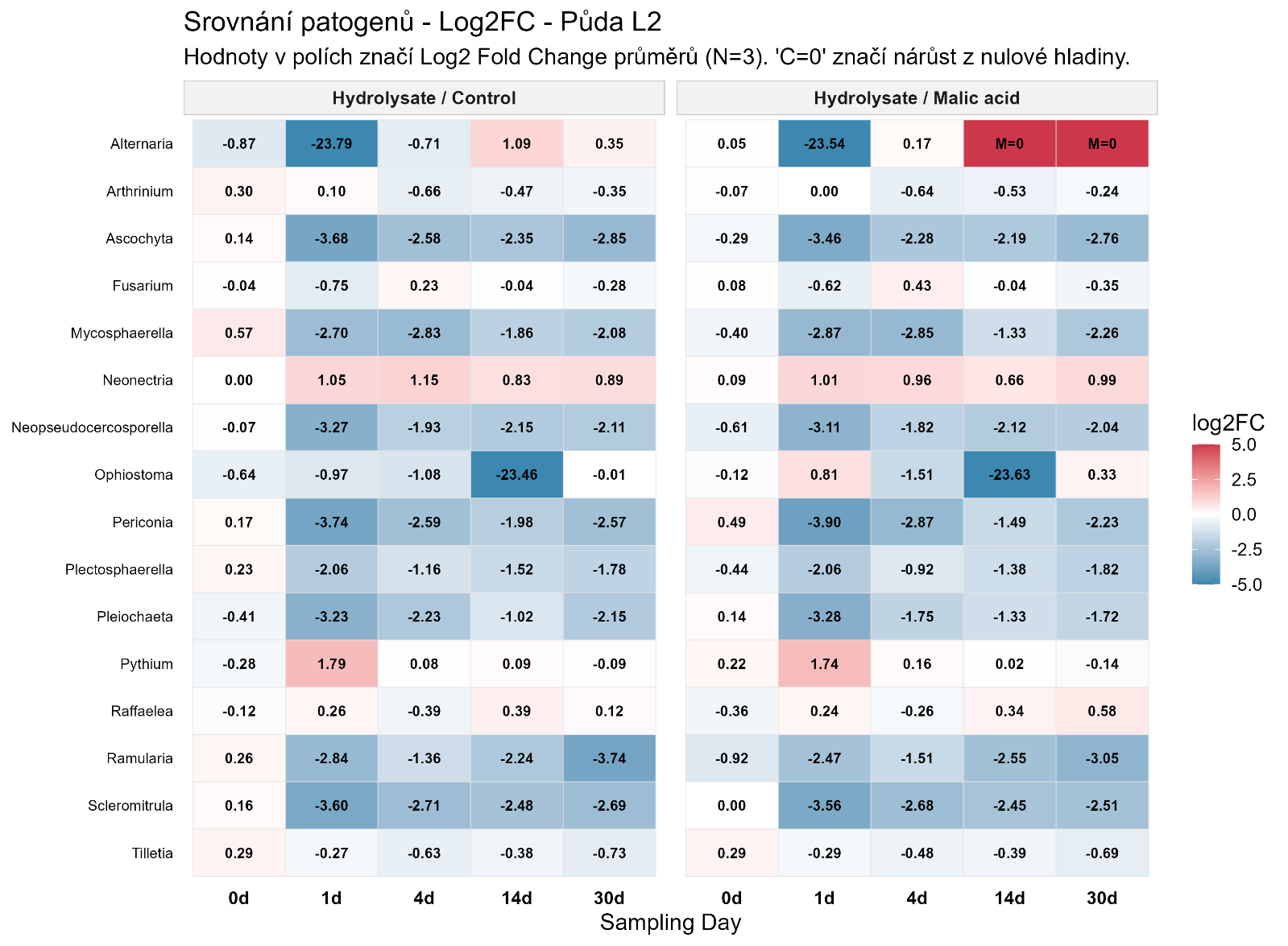

B

**Figure S9.** Temporal dynamics of the 16 most abundant phythopathogenic genera, shown as detected fractions (‰) in control (water, C) hydrolysate-amended (H), and malic acid-treatment (M) samples- Individual soil samples are shown as L1 (A) and L2 (B).

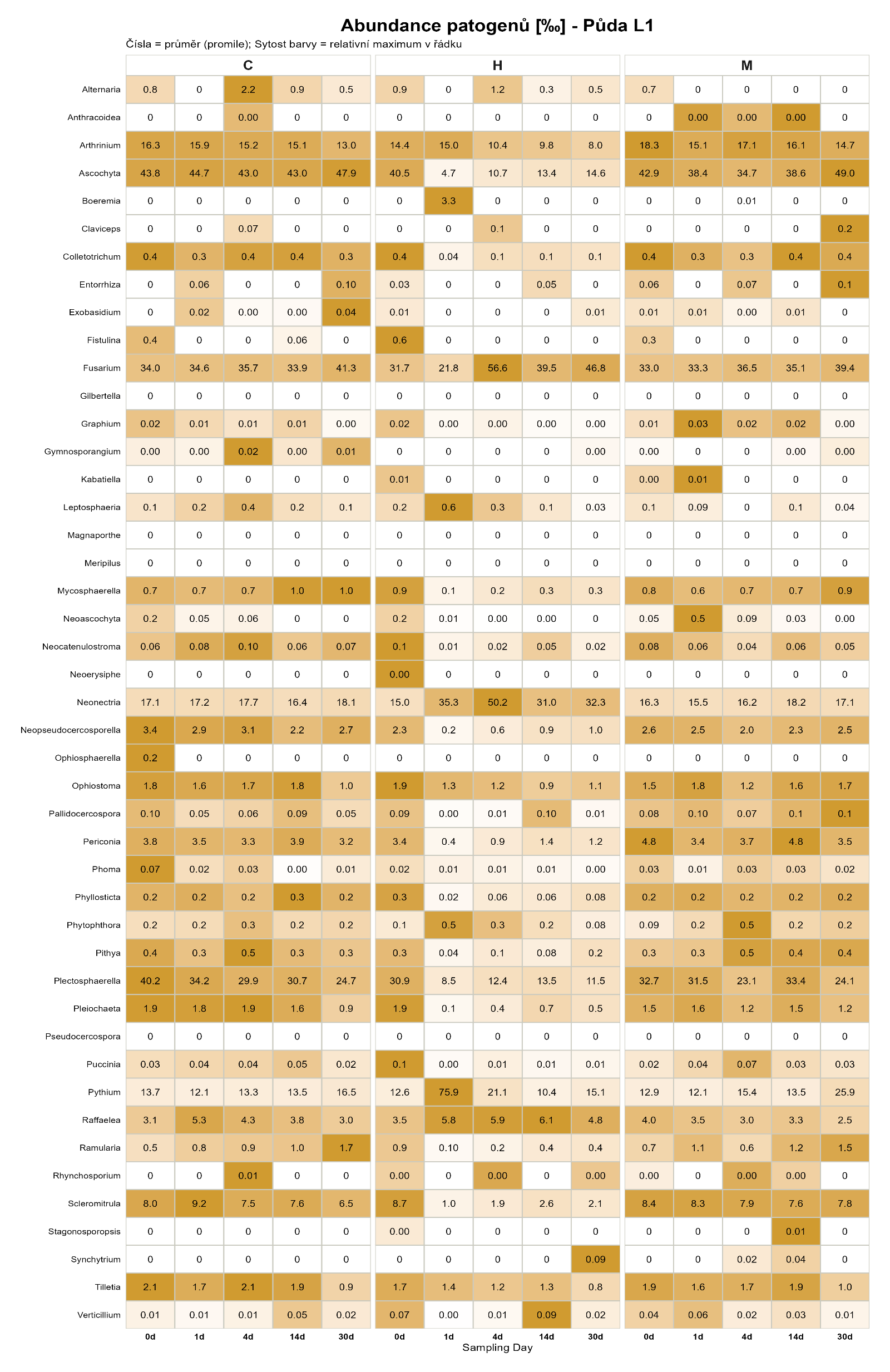

A

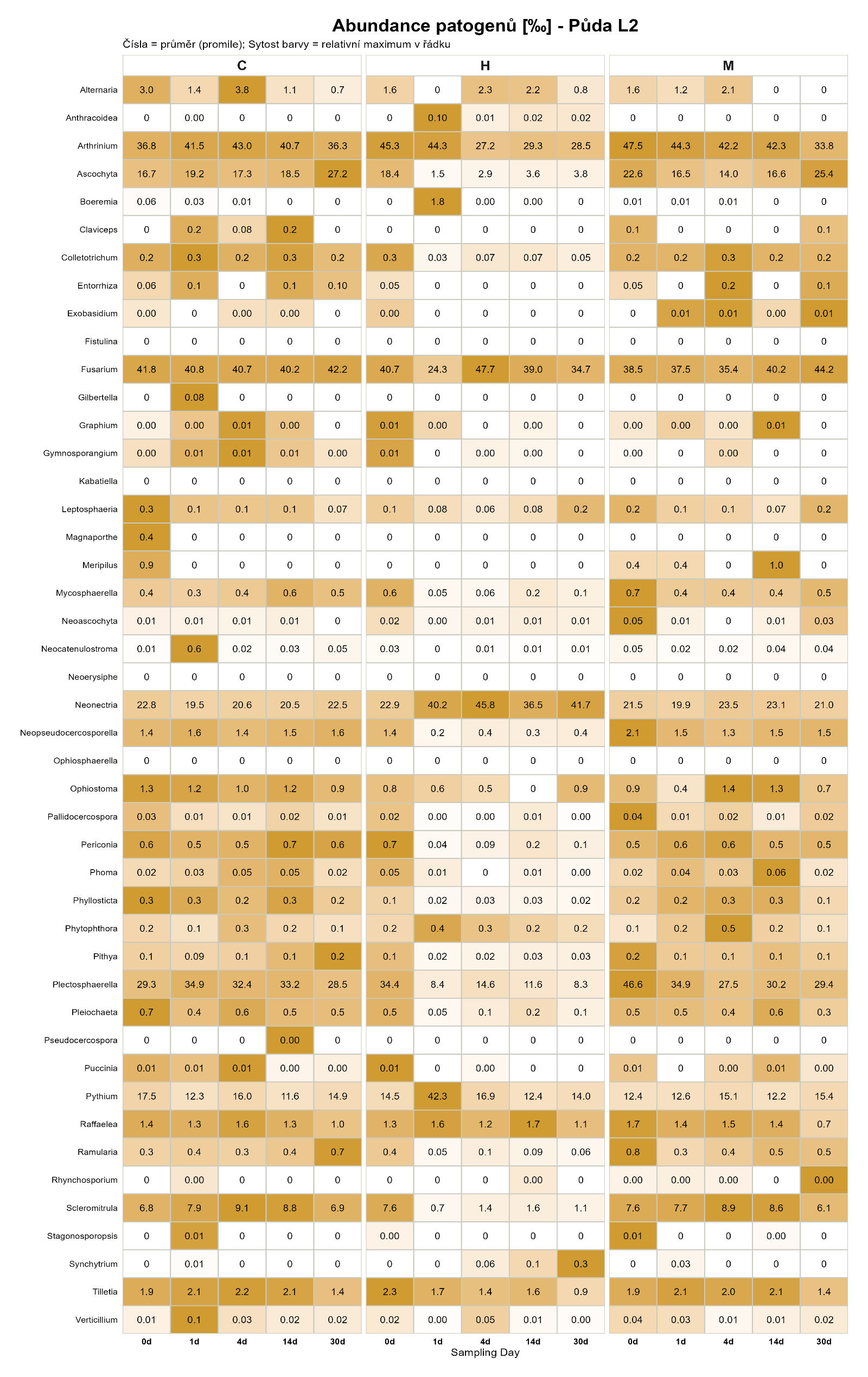

B

**Figure S10.** Temporal dynamics of all UNITE-annotated phythopathogenic genera, shown as detected fraction (‰) in control (water, C), hydrolysate-amended (H), and malic acid-treated (M) samples. Individual soil samples are shown for L1 (A) and L2 (B).

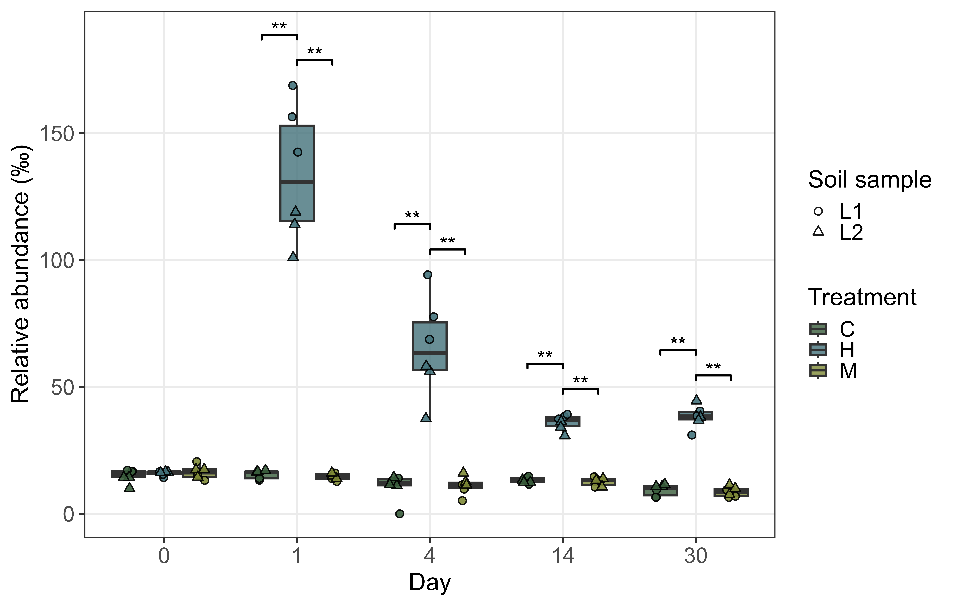

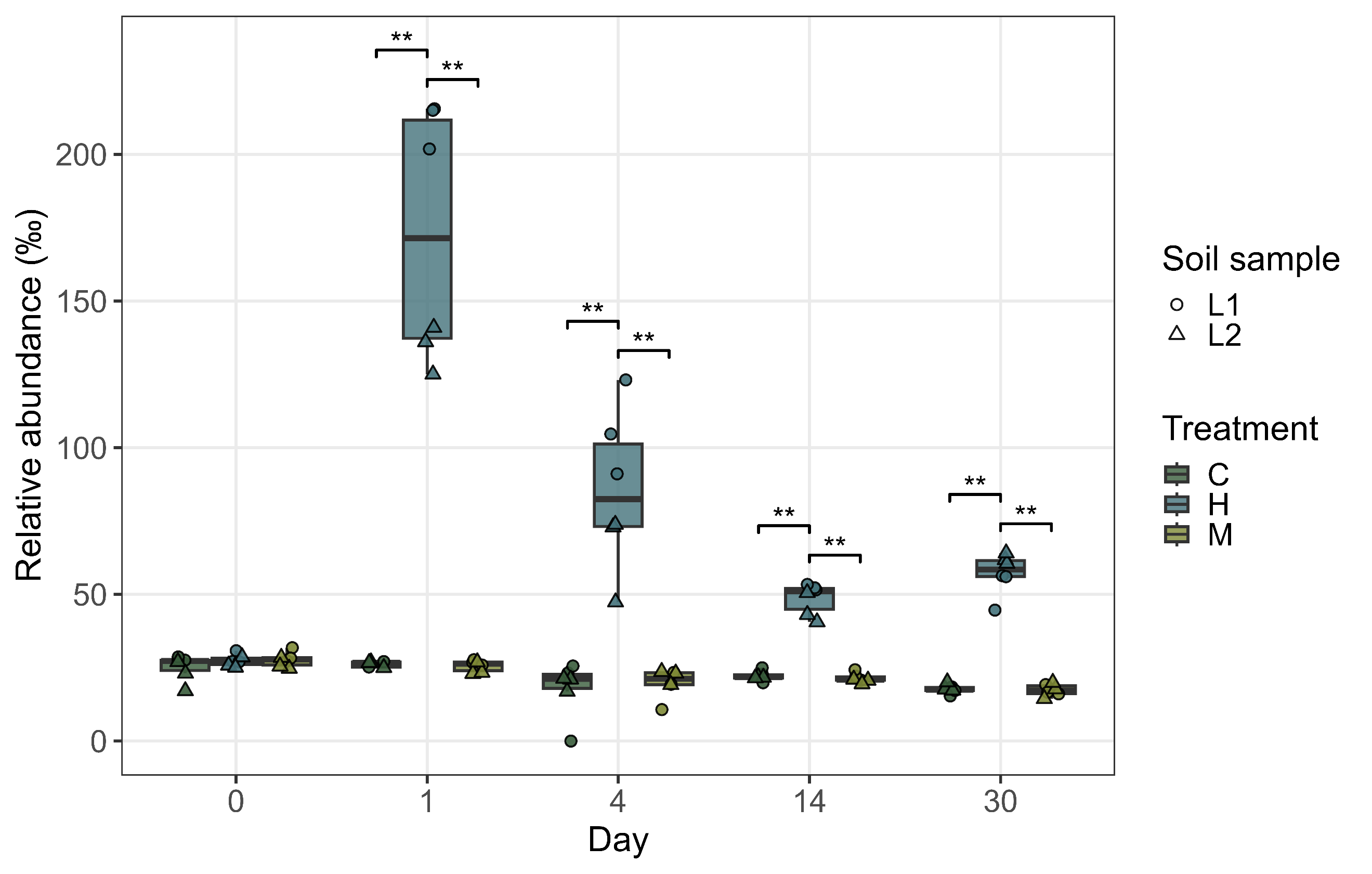

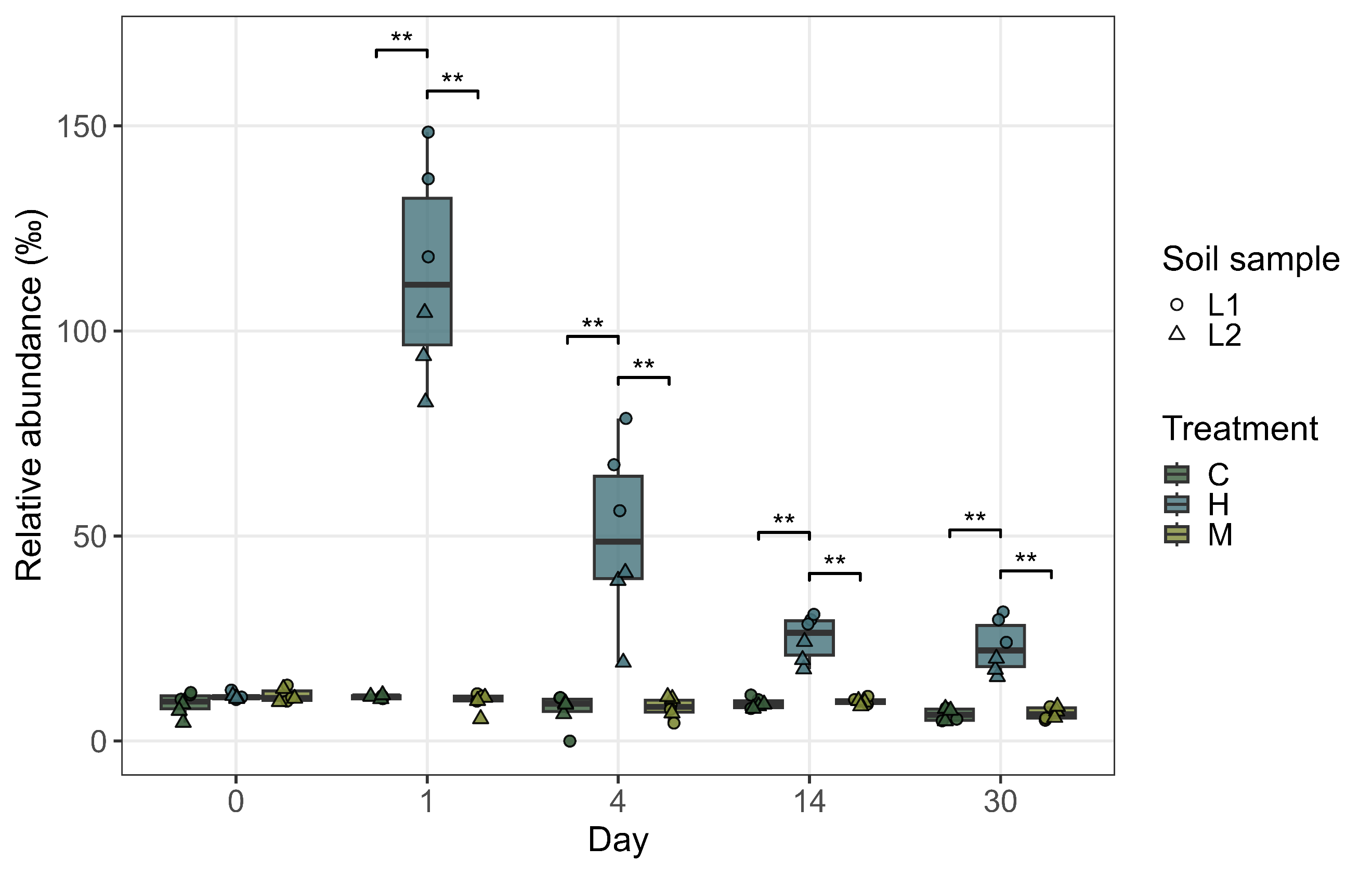

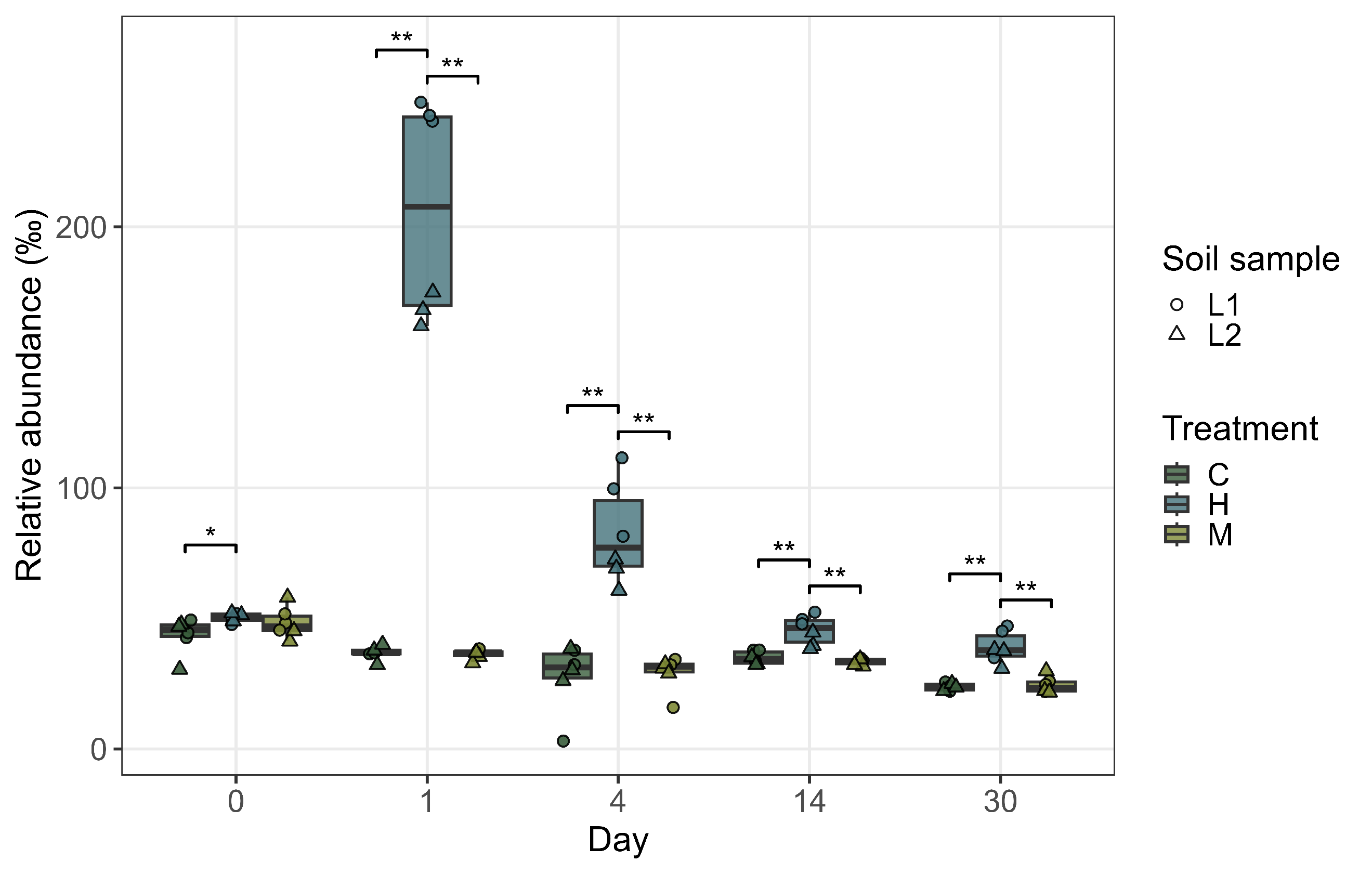

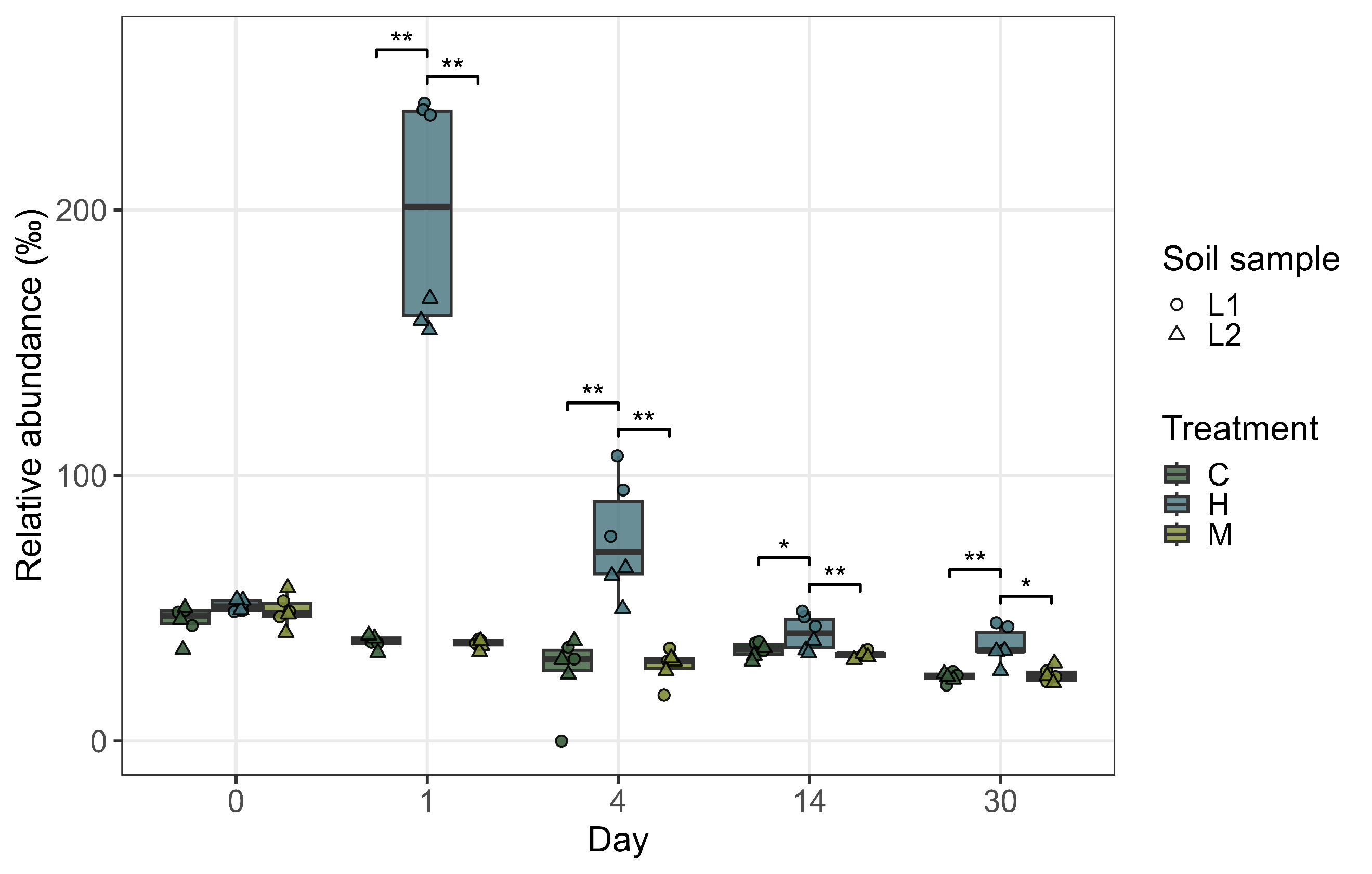

A

B

C

D

E

**Figure S11.** Temporal dynamics of the abundance of bacterial taxa belonging to specific PGPR subsets related to nutrient solubilisation: K (A), P (B), Zn (C); siderophore production (D) and pathogen suppression (E).

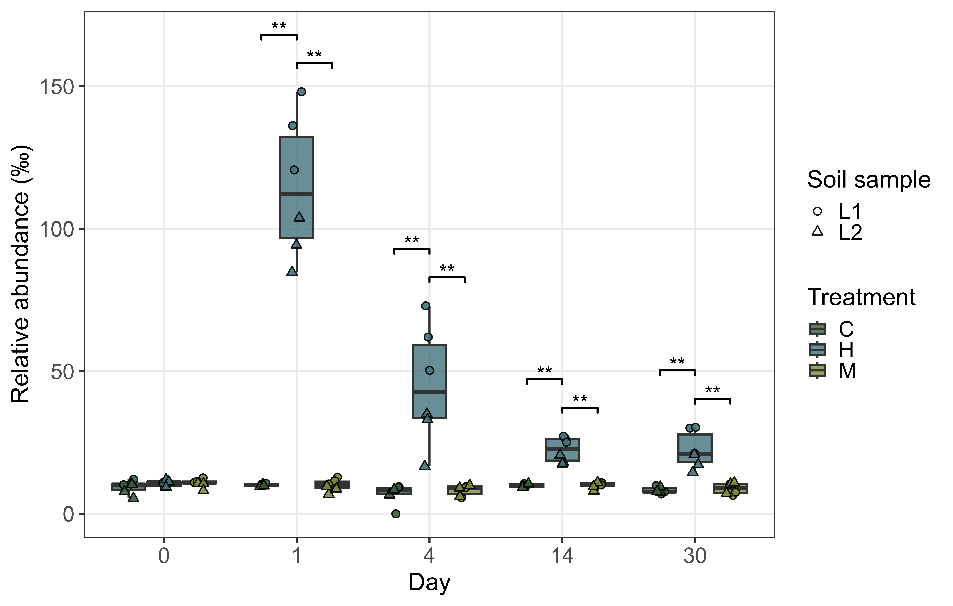

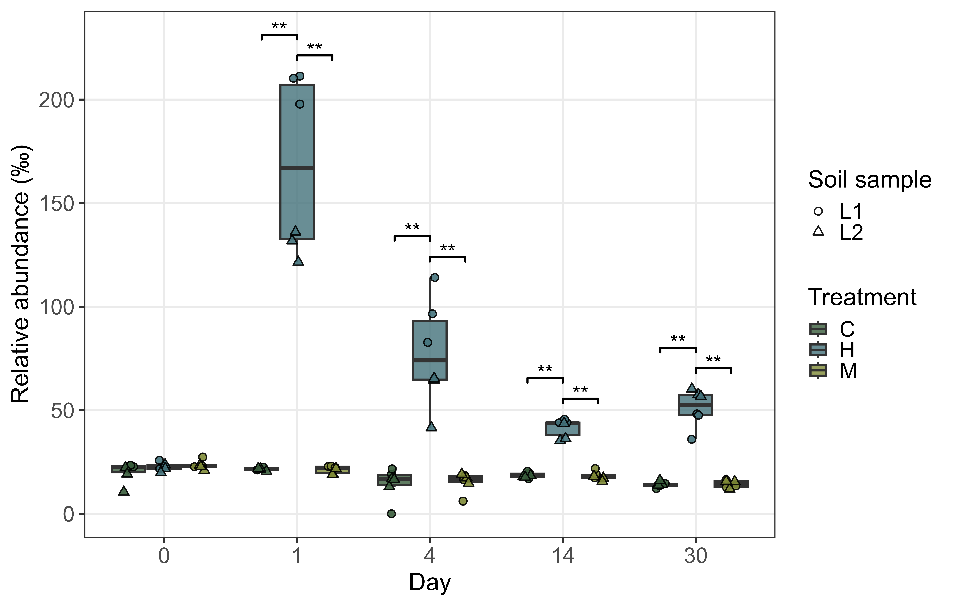

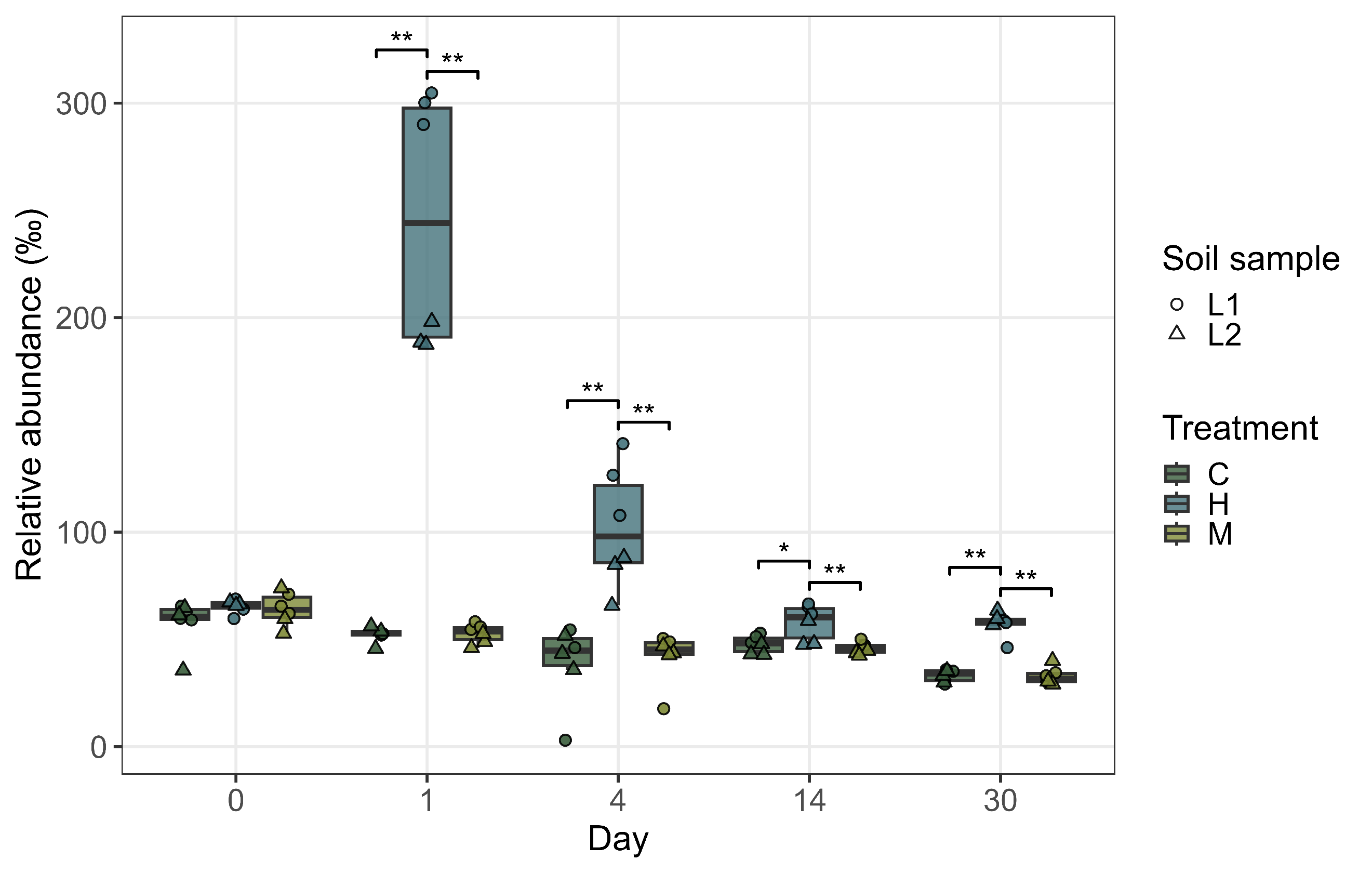

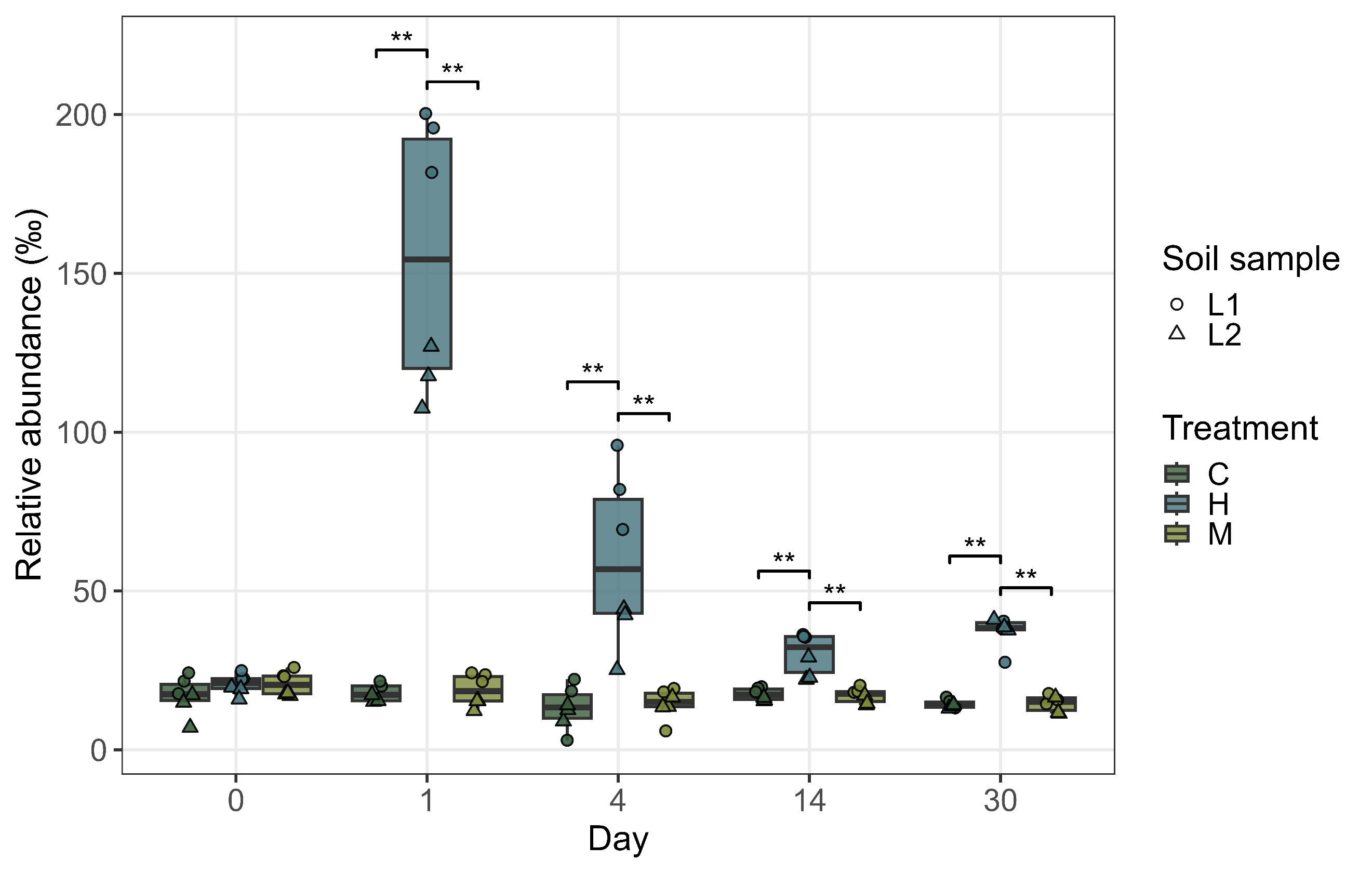

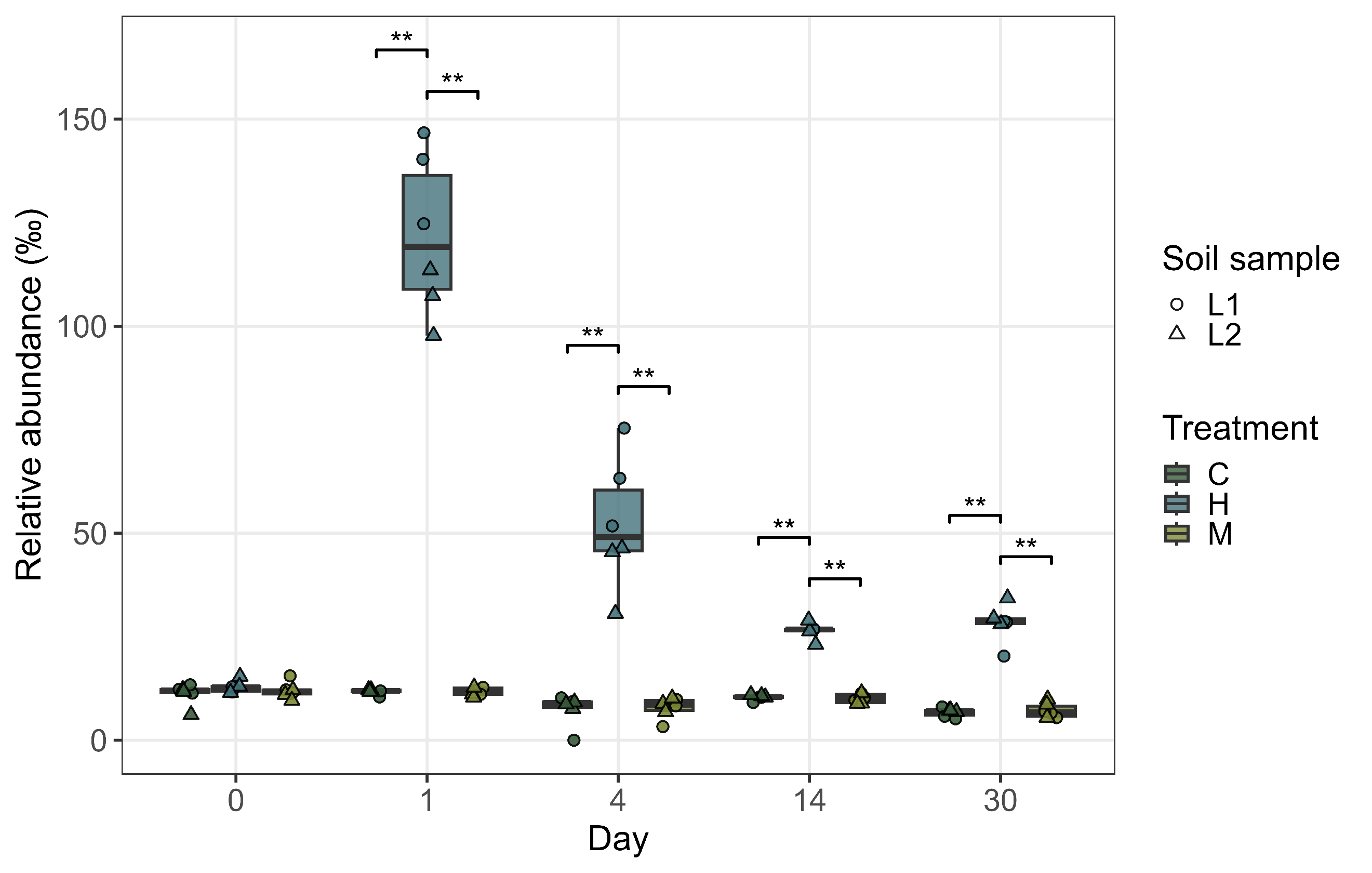

A

B

C

D

**Figure S12.** Temporal dynamics of the abundance of bacterial taxa belonging to specific PGPR subsets related to phytohormones production: ACC deaminase (A), auxin (B), cytokinine (C), GABA (D), ABA (E).

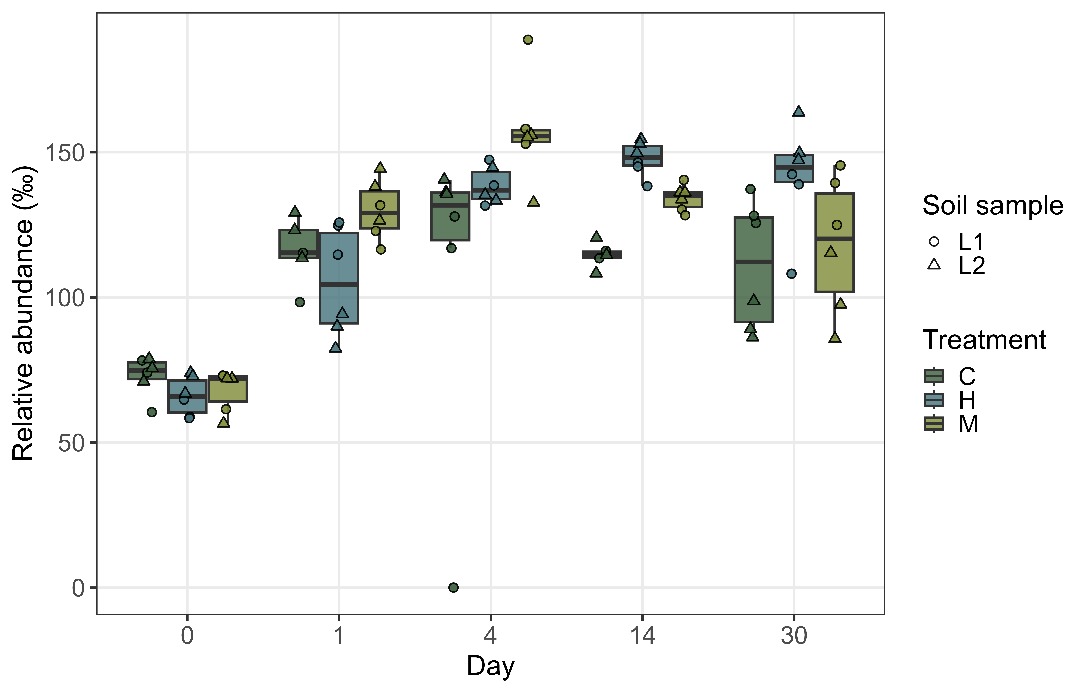

B

A

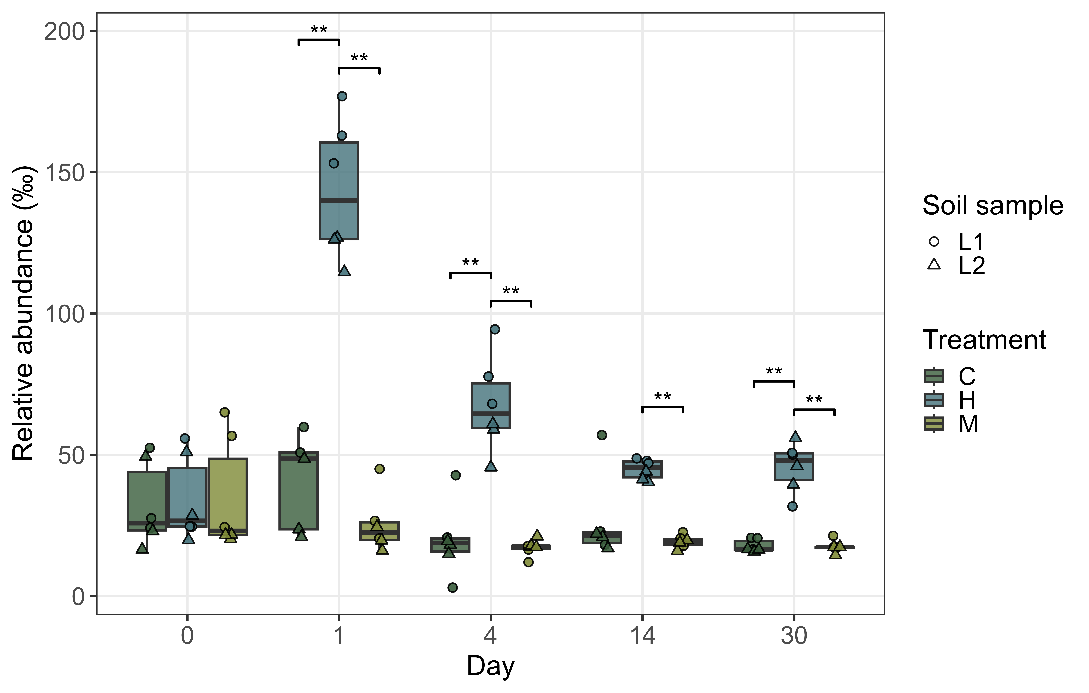

C

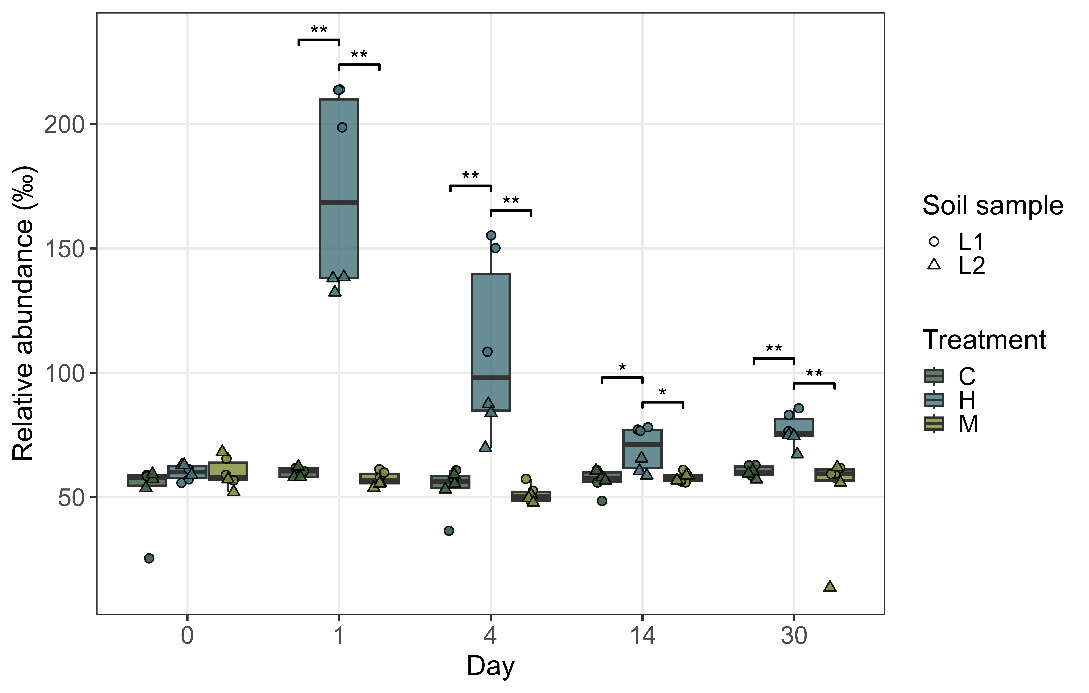

C

**Figure S13.** Temporal dynamics of the abundance of bacterial taxa classified as potential nitrogen fixers (A), nitrifiers (B), or denitrifiers (C).

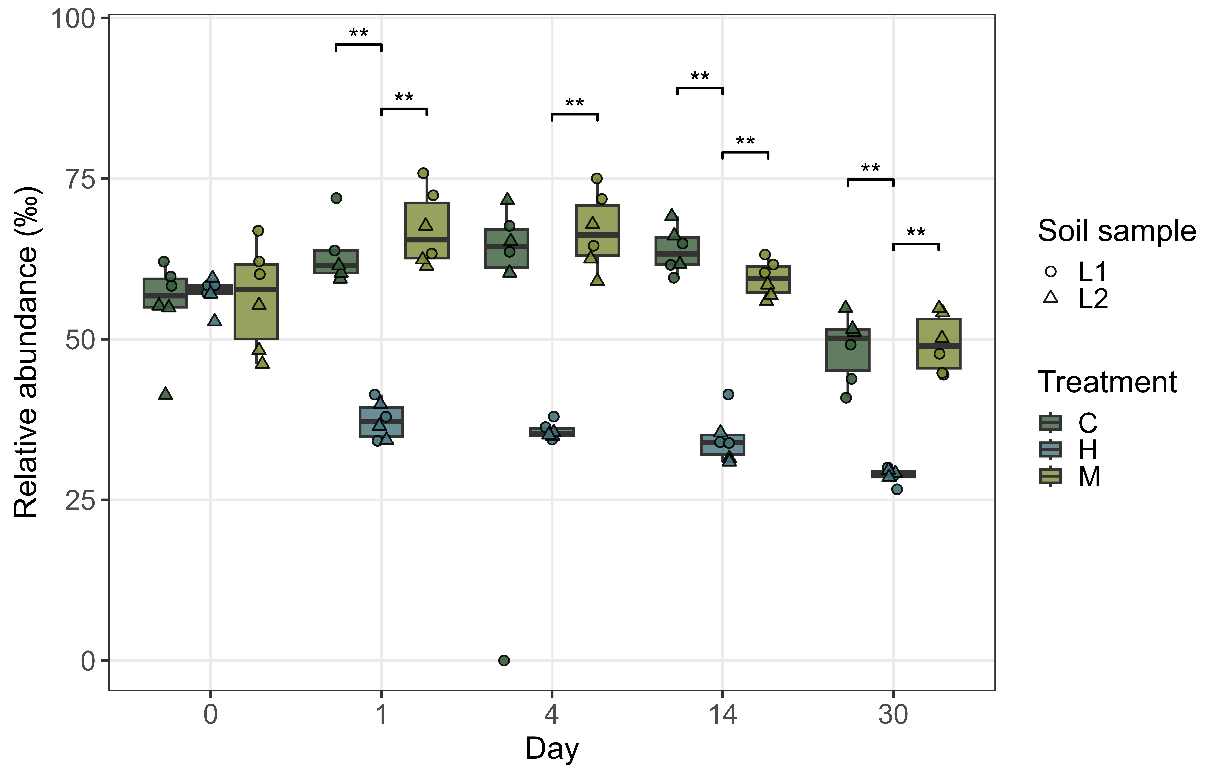

**Figure S14.** Temporal dynamics of the abundance of bacterial taxa classified as active carbon indicators.

| **Soil** | **C** | **N** | **Mg** | **Fe** | **Mn** | **Al** | **Na** | **K** |
| --- | --- | --- | --- | --- | --- | --- | --- | --- |
| mg / kg | 16.18 | 4.43 | 7.52 | 31.77 | 0.84 | 38.53 | 0.26 | 9.82 |

**Table S1.** Elemental composition of the experimental soil.

| **Dry matter in**  **hydrolysate** | **C** | **N** | **O** | **Na** | **P** | **S** | **K** | **Ca** |
| --- | --- | --- | --- | --- | --- | --- | --- | --- |
| **wt. (%)** | 43.9 | 14.9 | 39.5 | 0.1 | 0.1 | 1 | 0.5 | 0.2 |

**Table S2.** Elemental composition of the dry matter in feather hydrolysate.

| **Wet hydrolysate** | **Zn** | **As** | **Ba** | **Fe** | **Pb** | **Mg** | **Mn** | **Al** | **Na** | **Ca** | **Cr** |
| --- | --- | --- | --- | --- | --- | --- | --- | --- | --- | --- | --- |
| **mg/l** | 4.24 | <LOD | 0.2 | 5.885 | <LOD | 35.36 | 2.09 | 0.93 | 49.8 | 93.89 | 0.065 |

**Table S3.**  Elemental composition in liquid hydrolysate as determined by ICP-OES. < LOD stands for below the detection limit.

|  | **Ala** | **Arg** | **Asn** | **Asp** | **Phe** | **Gln** | **Glu** | **Gly** | **His** | **H-Lys** | **H-Pro** | **Ile+Leu** | **Lys** | **Met** | **Pro** | **Ser** | **Thr** | **Trp** | **Tyr** | **Val** |
| --- | --- | --- | --- | --- | --- | --- | --- | --- | --- | --- | --- | --- | --- | --- | --- | --- | --- | --- | --- | --- |
| **mg**/l | 95.6 | 21.6 | 4.1 | 1194.9 | 47.9 | 1.8 | 17.8 | 69.8 | 11.8 | 1.1 | 3.0 | 54.5 | 43.1 | 14.9 | 75.7 | 91.8 | 35.7 | 4.0 | 36.5 | 53.0 |
| ***wt.%*** | *5.1* | *1.2* | *0.2* | *63.6* | *2.6* | *0.1* | *0.9* | *3.7* | *0.6* | *0.1* | *0.2* | *2.9* | *2.3* | *0.8* | *4.0* | *4.9* | *1.9* | *0.2* | *1.9* | *2.8* |

**Table S4.**  Amino acid composition in liquid hydrolysate. Cystein and cystin were not detected.

| **Sample** | **pH, day 0** | **pH, day 20** | **pH, day 30** |
| --- | --- | --- | --- |
| L1, before treatment, | 5,59 |  |  |
| L1, before treatment, replicate | 6,08 |  |  |
| L1 C (water) |  | 6,24 | 6,20 |
| L1 M (malic acid) |  | 6,15 | 6,15 |
| L1 H (hydrolysate) |  | 5,92 | 6,84 |
| L2, before treatment | 5,97 |  |  |
| L2, before treatment, replicate | 5,93 |  |  |
| L2 C (water) |  | 5,94 | 5,94 |
| L2 M (malic acid) |  | 5,89 | 5,91 |
| L2 H (hydrolysate) |  | 5,74 | 5,63 |

***Table S5.*** *pH profiles of the soil samples.*
