## Supplementary_data2 for "Organic-acid-derived feather hydrolysate drives soil microbiome succession toward copiotrophic and putatively plant-beneficial taxa"

**Supplementary data 2**

***Organic-acid-derived feather hydrolysate drives soil microbiome succession toward copiotrophic and putatively plant-beneficial taxa* Veselský J. et al.**

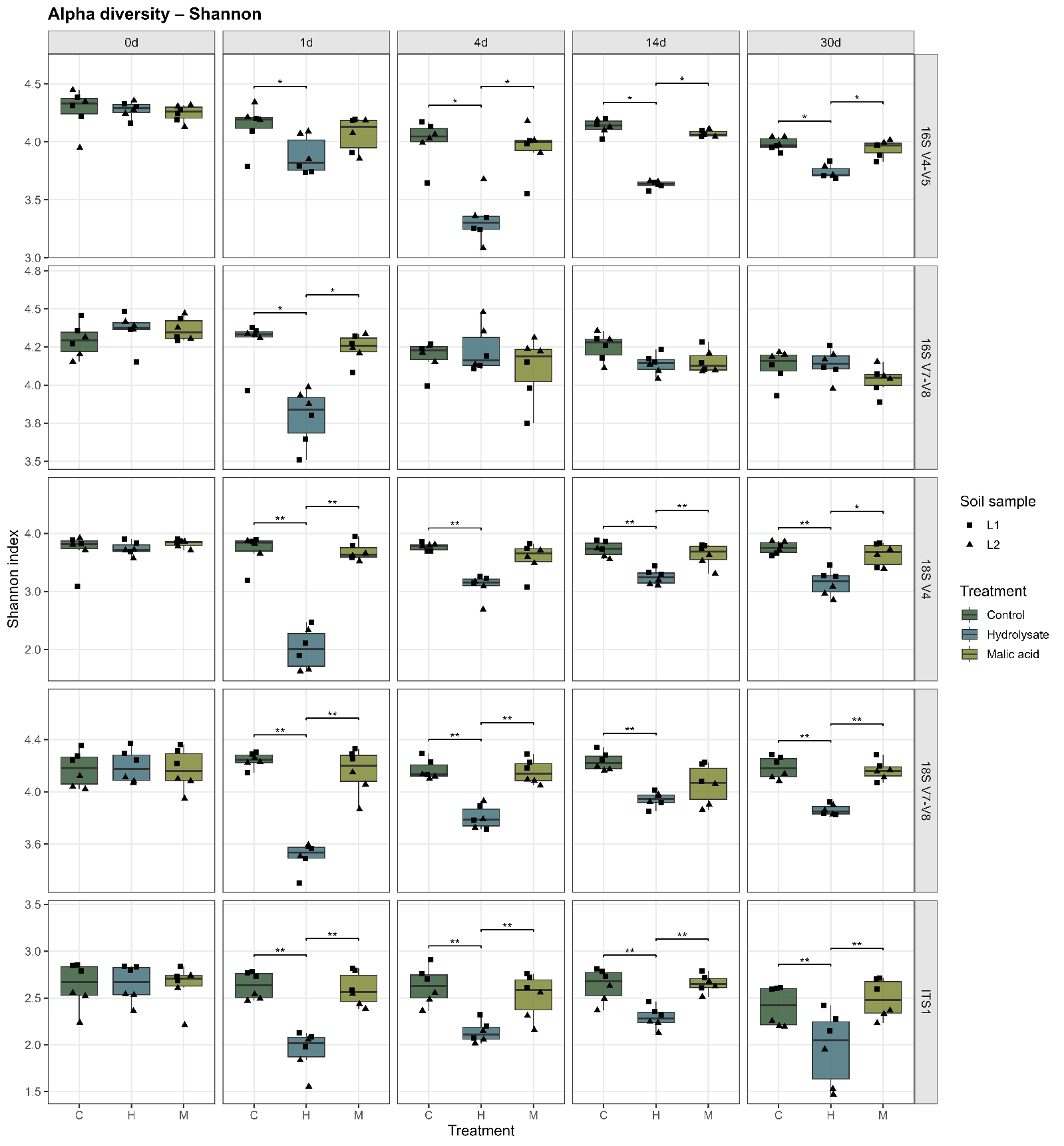

**Figure S15.** Alpha diversity (Shannon index) of the soil microbial community 0–30 days after hydrolysate amendment. Boxes represent the interquartile range; whiskers indicate the range excluding outliers. Statistical significance (p < 0.05) is based on Mann‑Whitney U test followed by Benjamini–Hochberg correction. The metrics shown here are inferred from individual sequencing subsets as depicted on the right side of the panels. For more details, see Table S6.

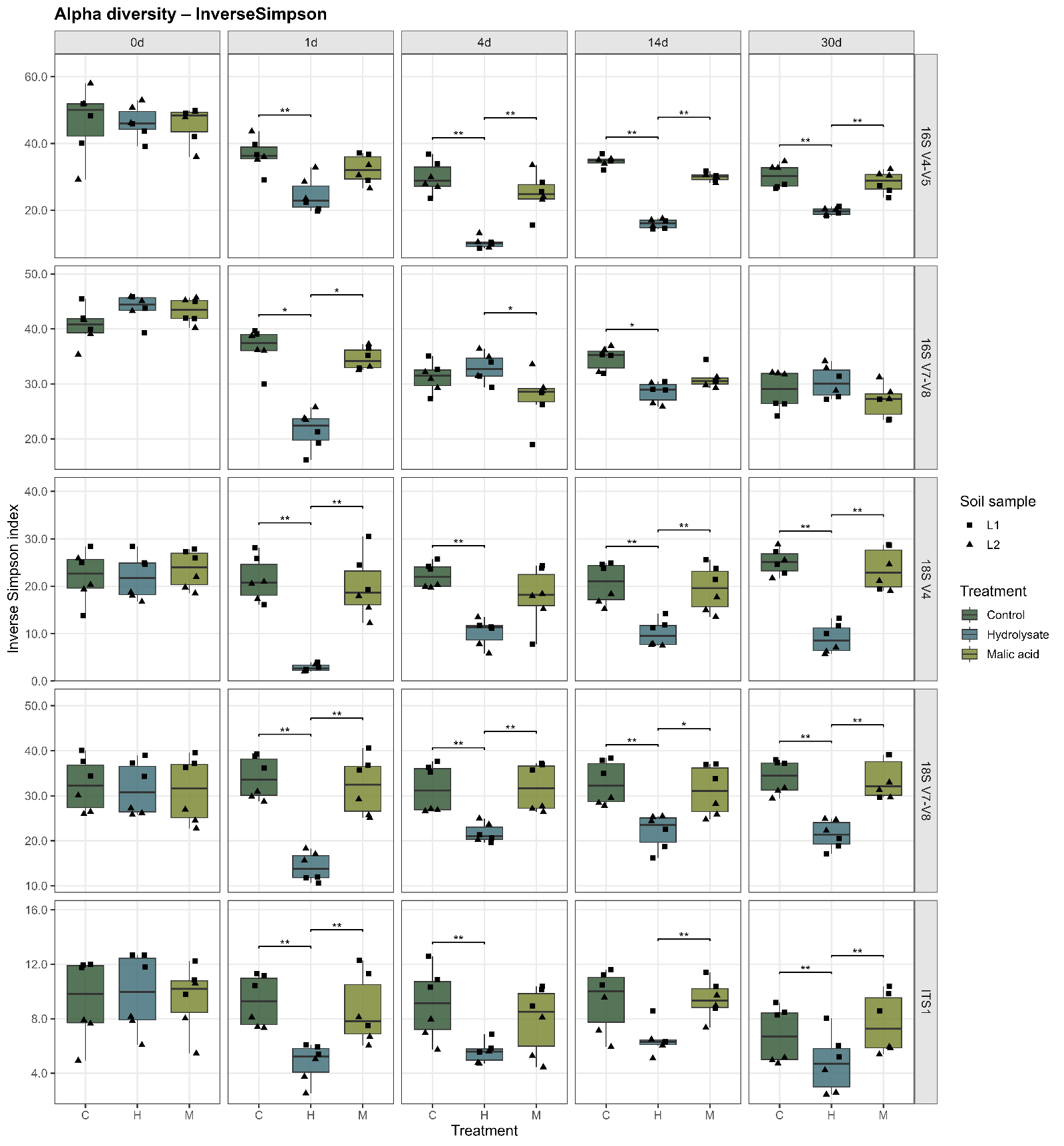

**Figure S16.** Alpha diversity (Inversed Simpson index) of the soil microbial community 0–30 days after hydrolysate amendment. Boxes represent the interquartile range; whiskers indicate the range excluding outliers. Statistical significance (p < 0.05) is based on Mann‑Whitney U test followed by Benjamini–Hochberg correction. The metrics shown here are inferred from individual sequencing subsets as depicted on the right side of the panels. For more details, see Table S6.

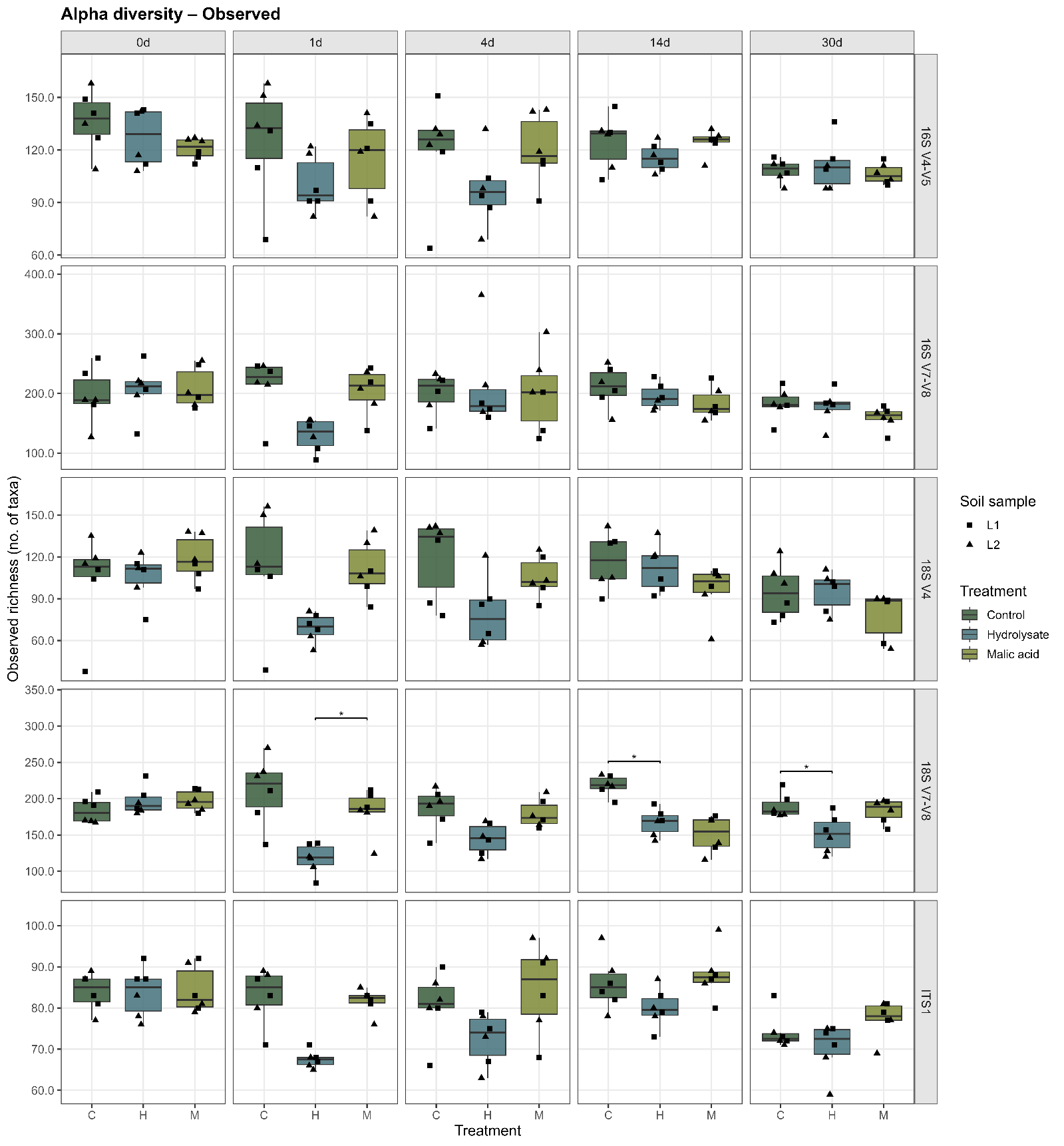

**Figure S17.** Alpha diversity (number of observed taxa) of the soil microbial community 0–30 days after hydrolysate amendment. Boxes represent the interquartile range; whiskers indicate the range excluding outliers. Statistical significance (p < 0.05) is based on Mann‑Whitney U test followed by Benjamini–Hochberg correction. The metrics shown here are inferred from individual sequencing subsets as depicted on the right side of the panels. For more details, see Table S6.

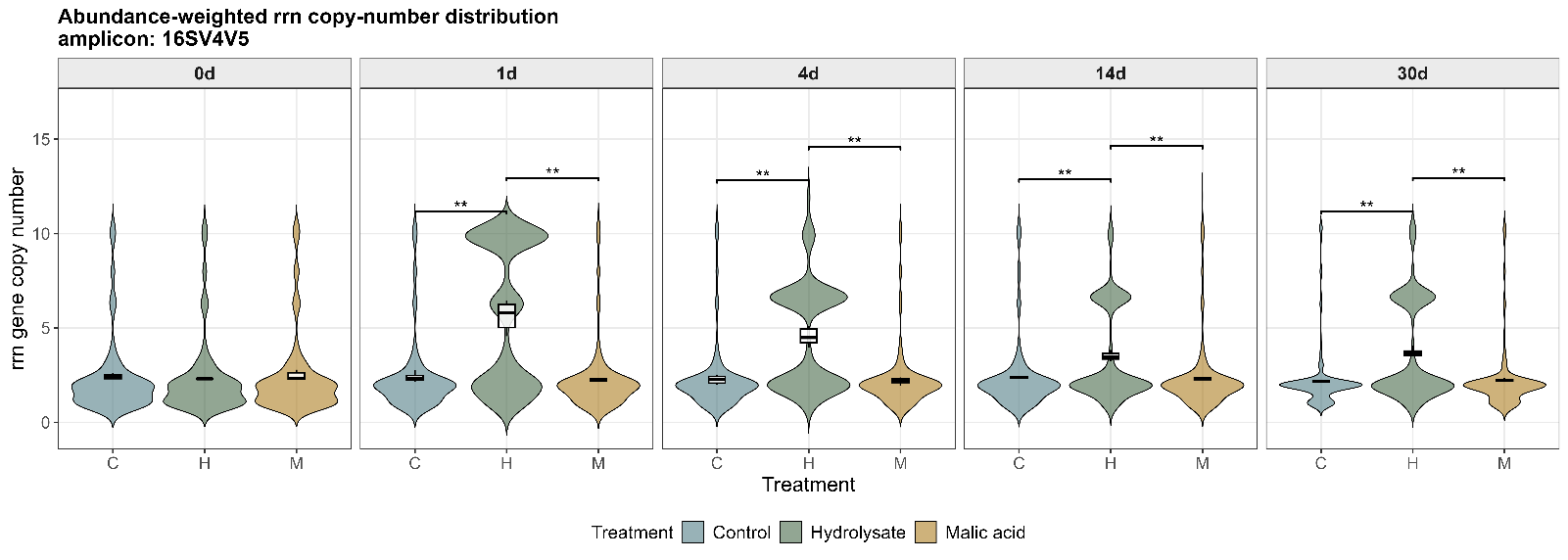

A

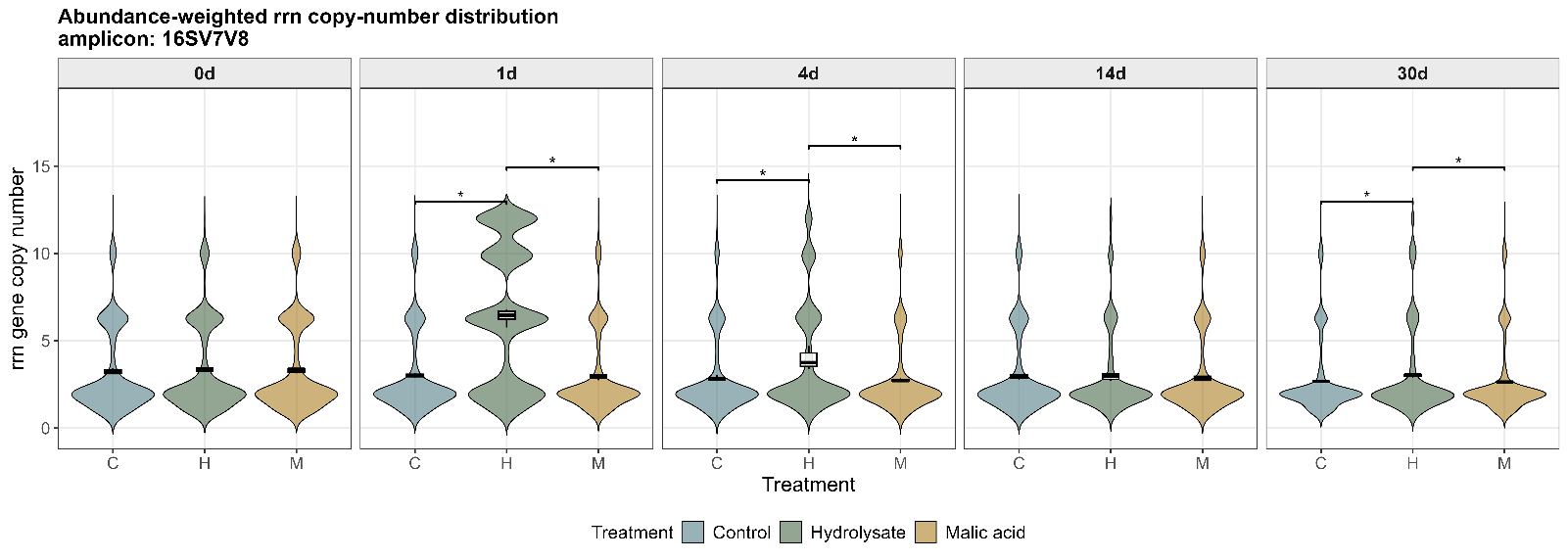

B

*Figure S18. Temporal dynamics of predicted rrn gene copy number abundance. Genera with predicted low rrn copy number are generally classified as oligotrophic, and genera with higher rrn copy number as copiotrophic (Bei et al., 2025). Statistical significance (indicated by an asterisk, p < 0.05) is based on Mann-Whitney U test followed by Benjamini–Hochberg correction. A) Inferred from the sequencing subset “set1” (variable regions V4 and V5 of the 16S); B) Inferred from the sequencing subset “set2” (variable regions V7 and V8 of the 16S), see Table S6 for details.*

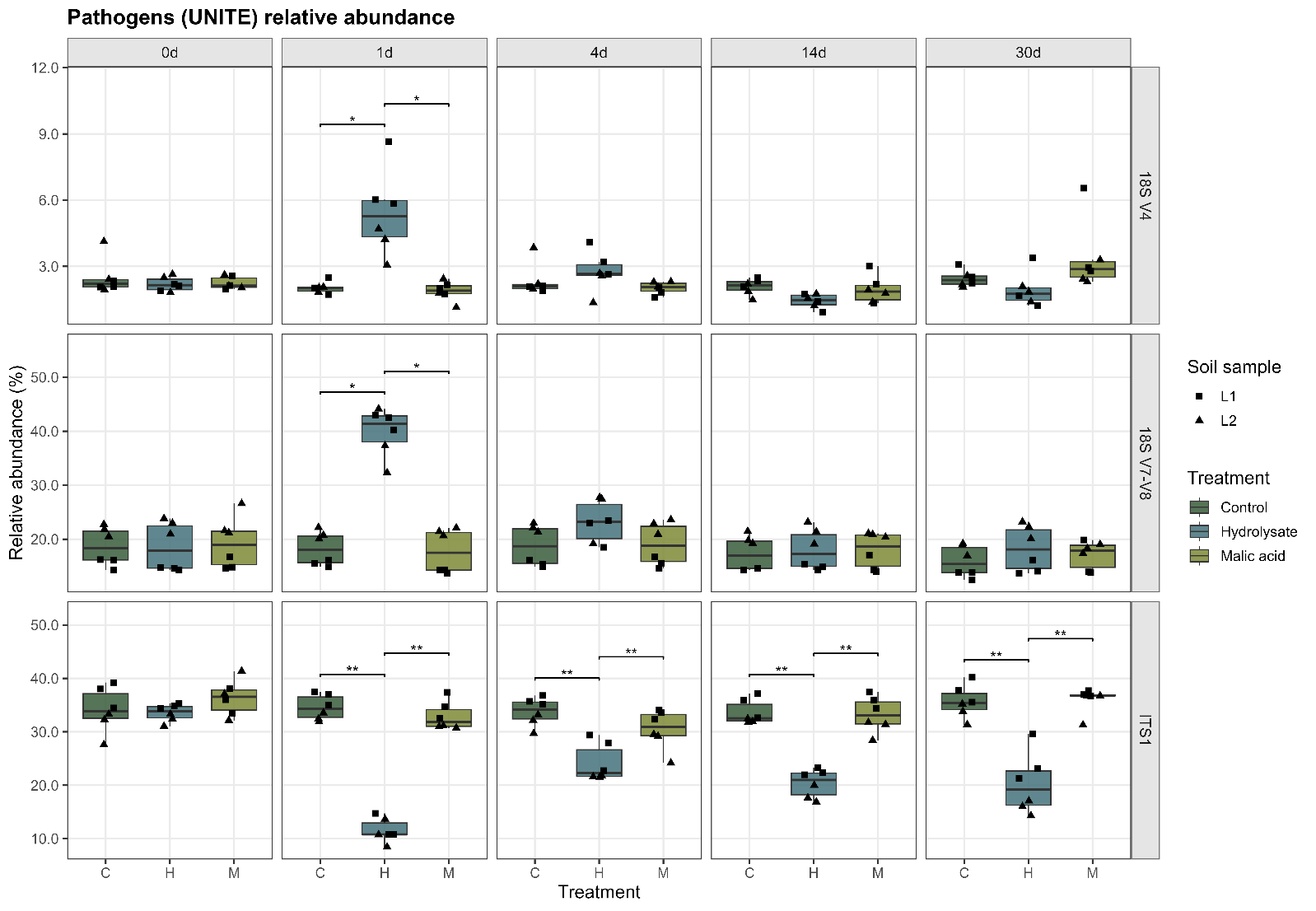

**Figure S19.** Decrease in the relative abundance of phytopathogenic genera based on UNITE predictions. Inferred from the sequencing subsets: “set3” (variable region V4 of the 18S); “set4” (variable regions V7 and V8 of the 18S), “ITS set”; see Table S6 for details.

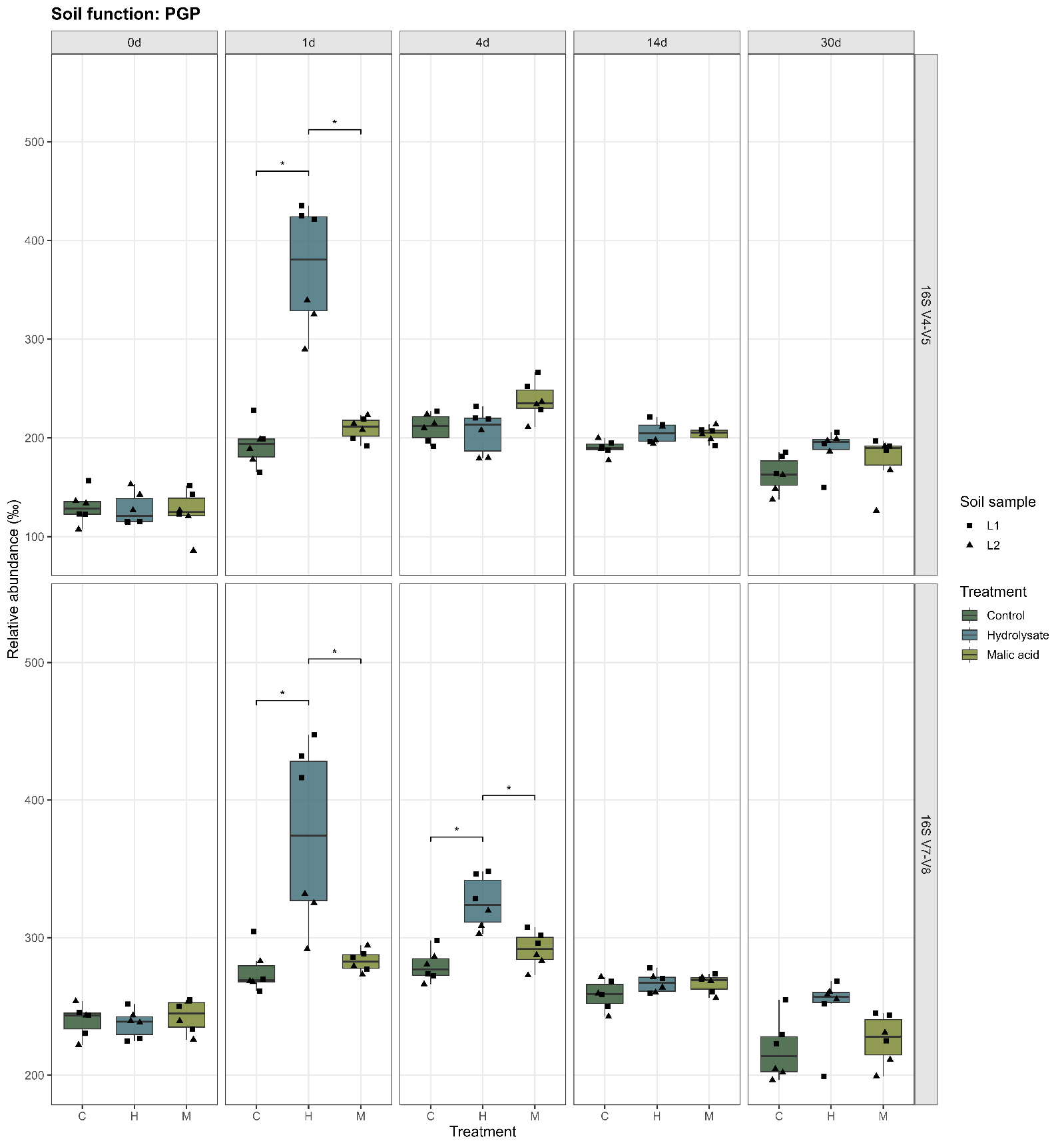

**Figure S20.** Temporal dynamics of the abundance of bacterial taxa classified as PGPR. Inferred from the sequencing subset “set1” (variable regions V4 and V5 of the 16S) or subset “set2” (variable regions V7 and V8 of the 16S), see Table S6 for details.

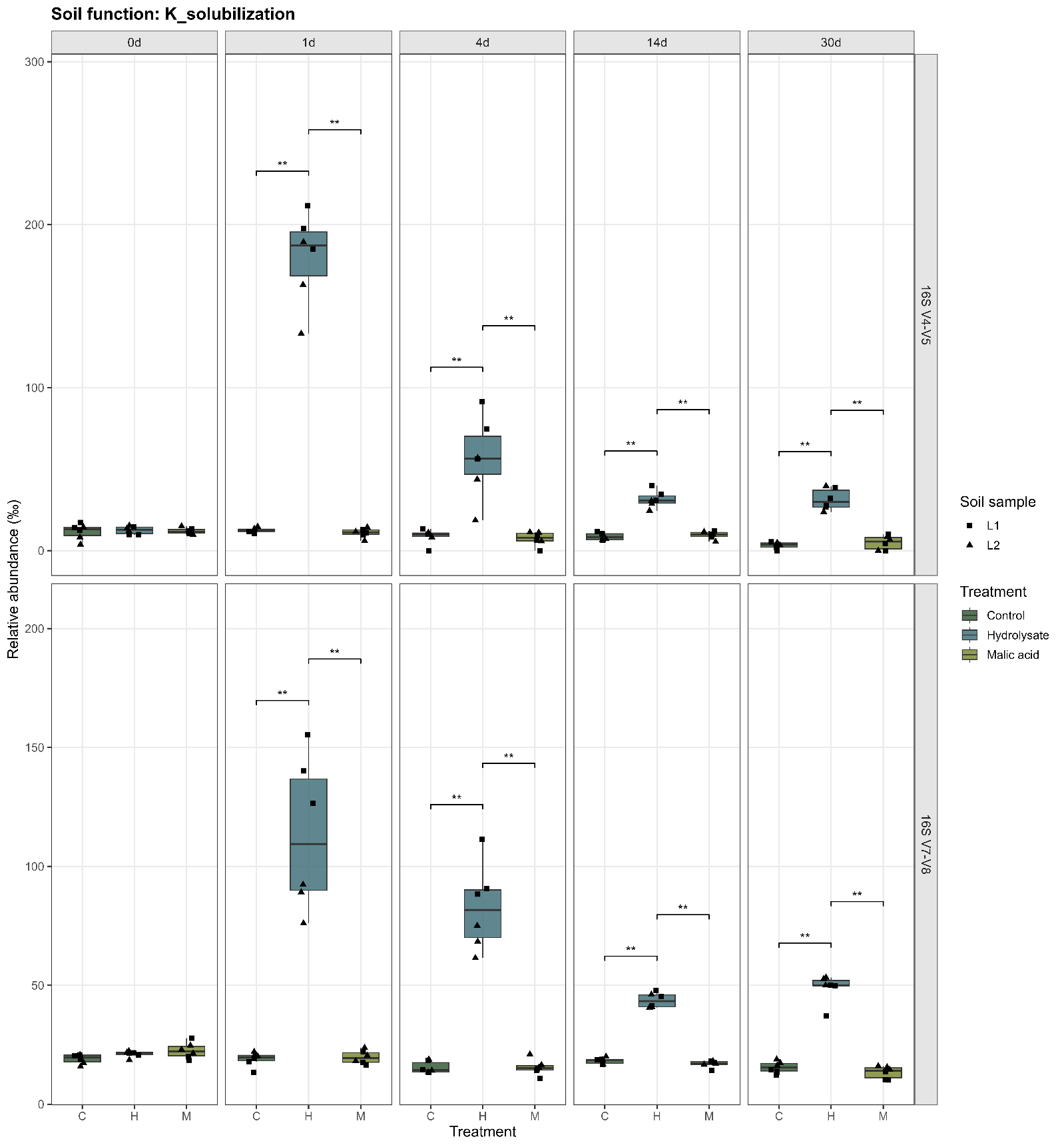

**Figure S21.** Temporal dynamics of the abundance of bacterial taxa classified as putative K solubilizers. A) Inferred from the sequencing subset “set1” (variable regions V4 and V5 of the 16S) or subset “set2” (variable regions V7 and V8 of the 16S), see Table S6 for details.

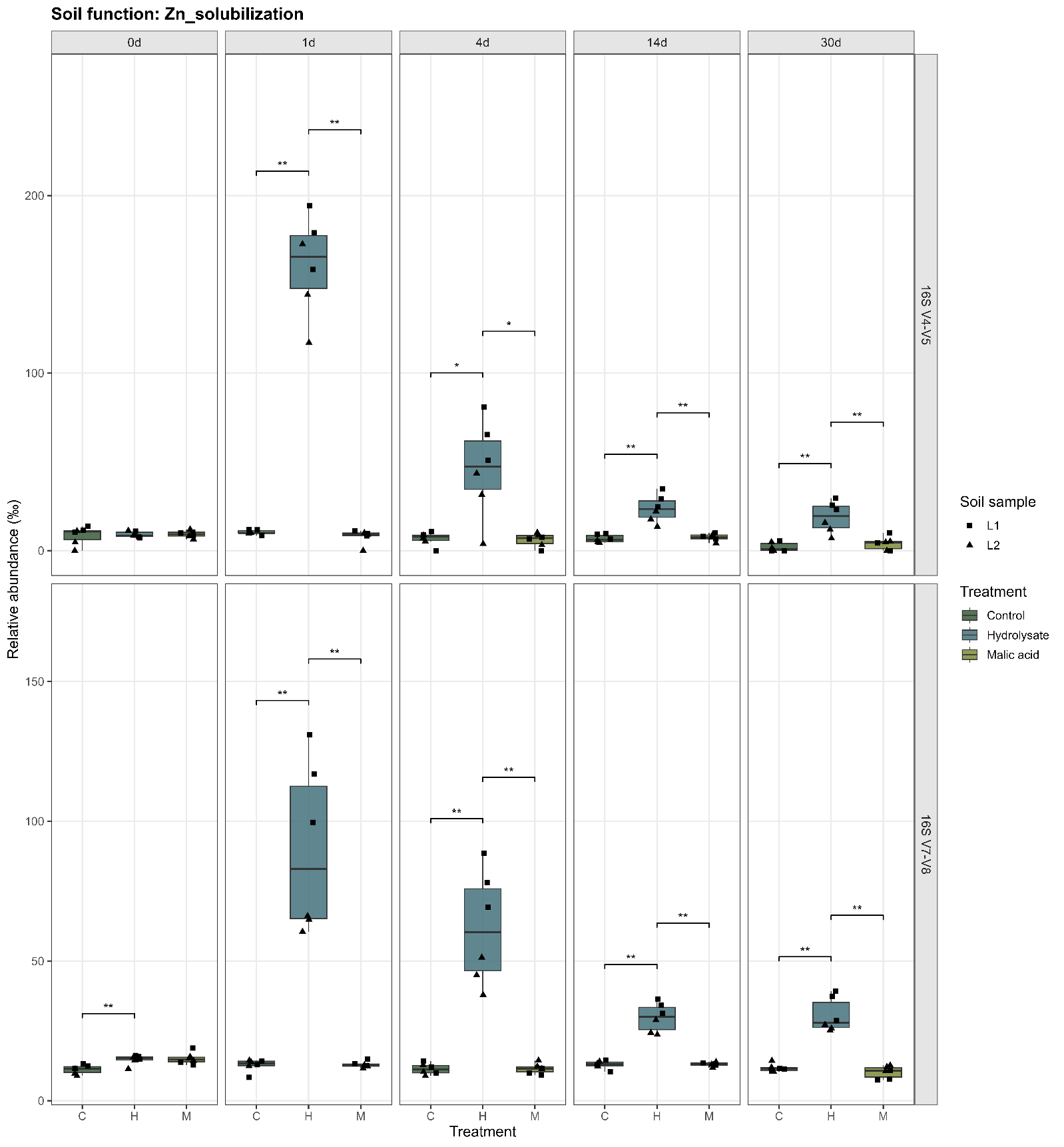

**Figure S22.** Temporal dynamics of the abundance of bacterial taxa classified as putative Zn solubilizers. A) Inferred from the sequencing subset “set1” (variable regions V4 and V5 of the 16S) or subset “set2” (variable regions V7 and V8 of the 16S), see Table S6 for details.

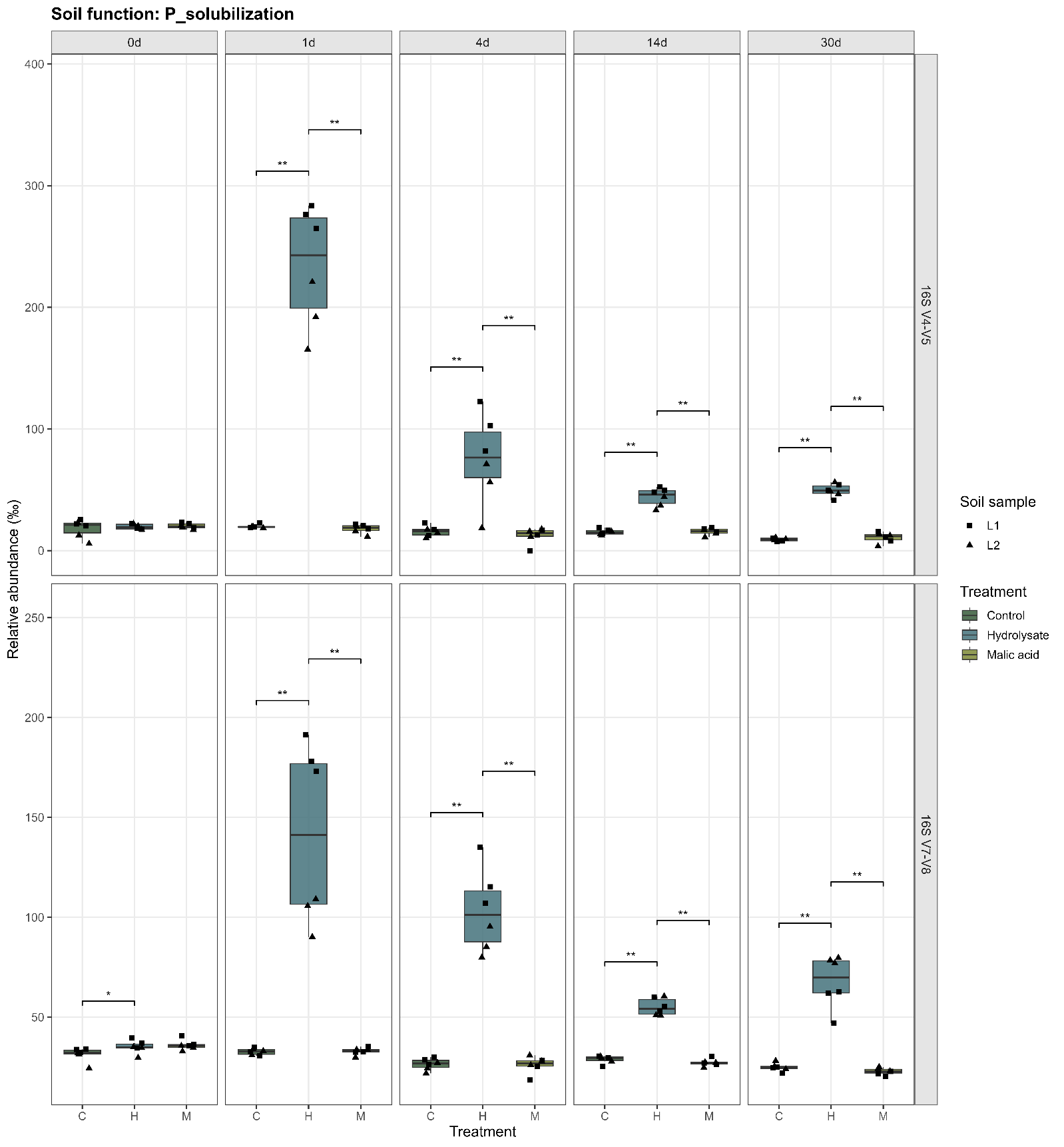

**Figure S23.** Temporal dynamics of the abundance of bacterial taxa classified as putative P solubilizers. A) Inferred from the sequencing subset “set1” (variable regions V4 and V5 of the 16S) or subset “set2” (variable regions V7 and V8 of the 16S), see Table S6 for details.

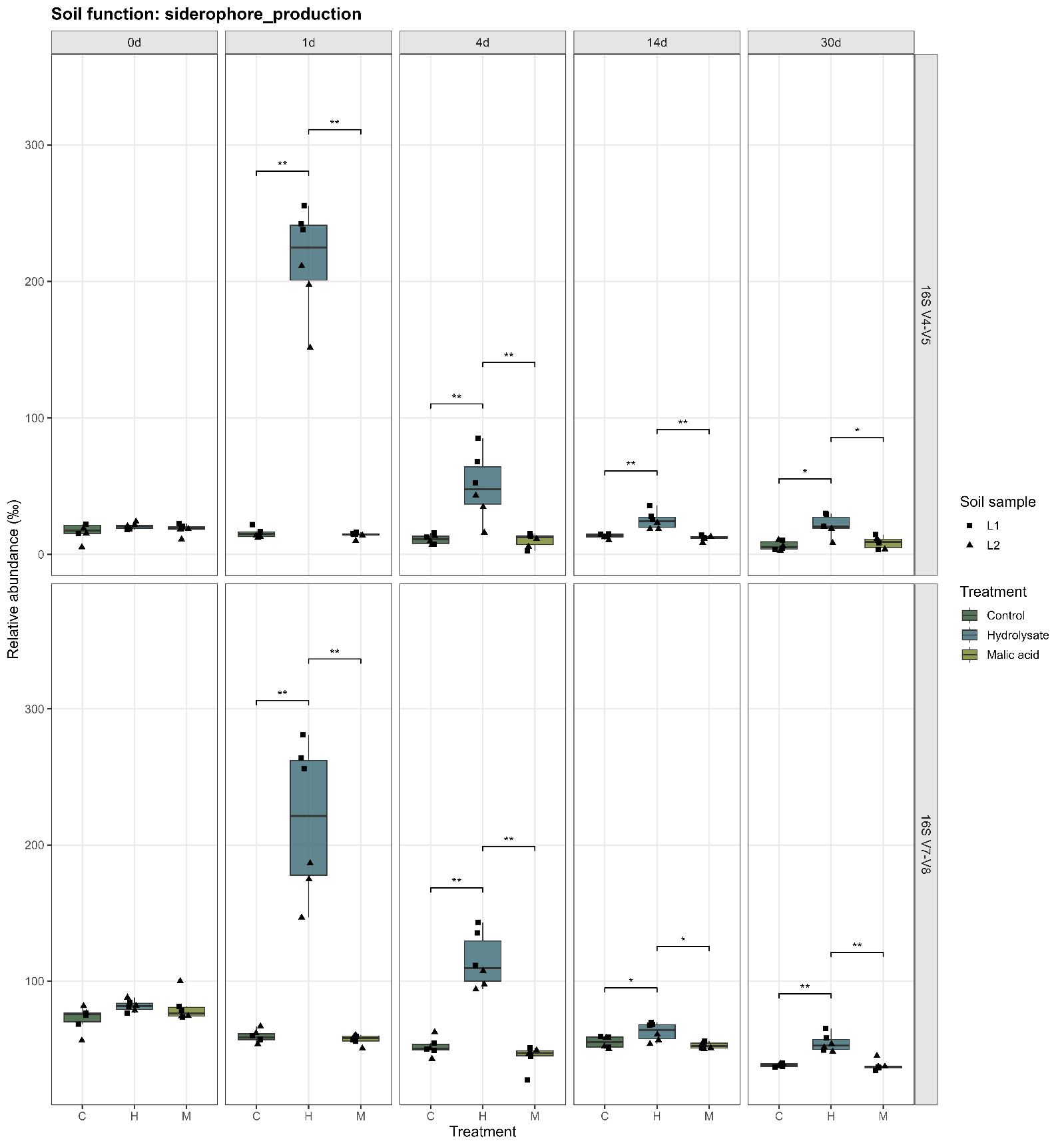

**Figure S24.** Temporal dynamics of the abundance of bacterial taxa classified as putative siderophore producers. A) Inferred from the sequencing subset “set1” (variable regions V4 and V5 of the 16S) or subset “set2” (variable regions V7 and V8 of the 16S), see Table S6 for details.

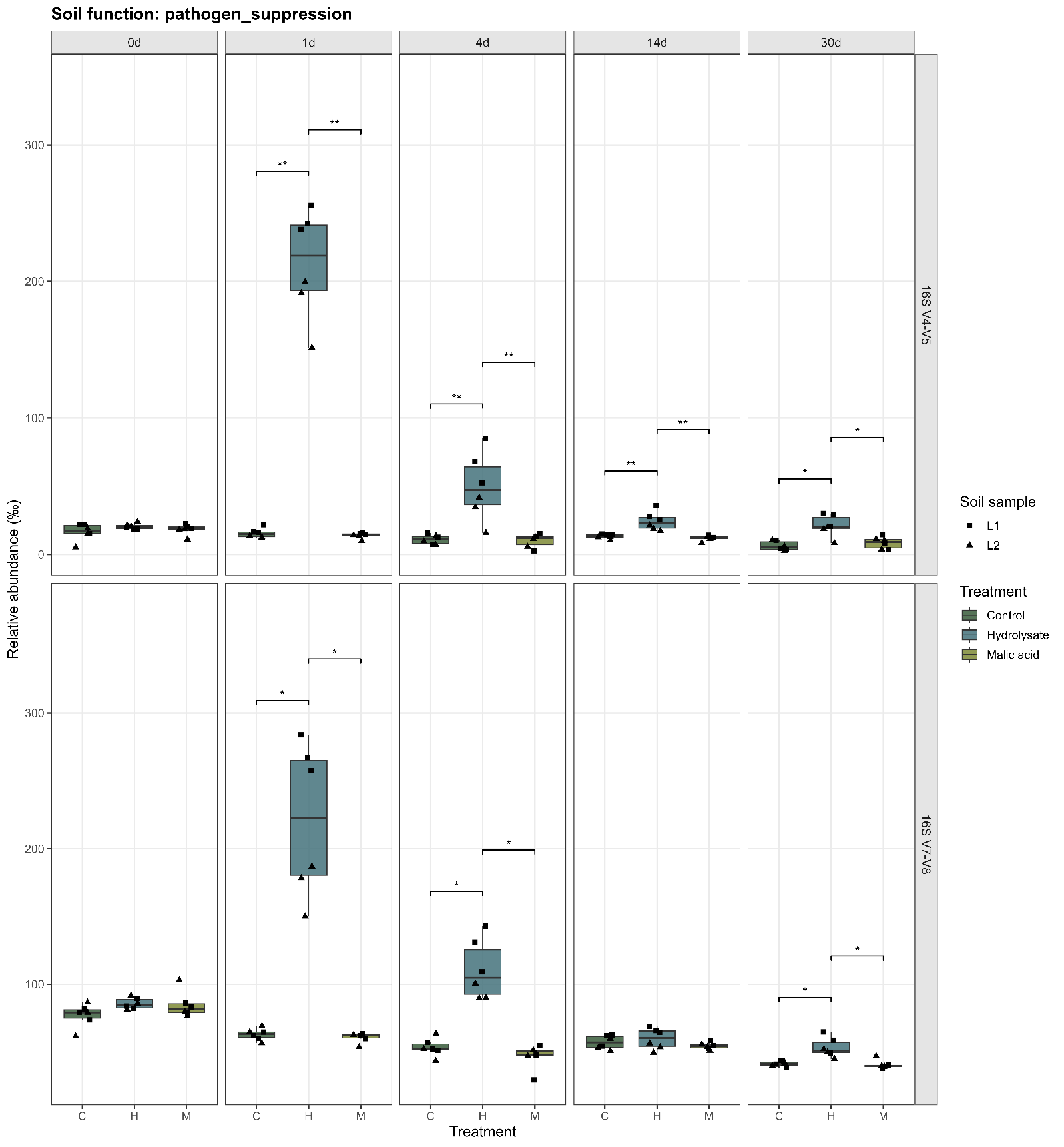

**Figure S25.** Temporal dynamics of the abundance of bacterial taxa classified as putative pathogen suppressors. A) Inferred from the sequencing subset “set1” (variable regions V4 and V5 of the 16S) or subset “set2” (variable regions V7 and V8 of the 16S), see Table S6 for details.

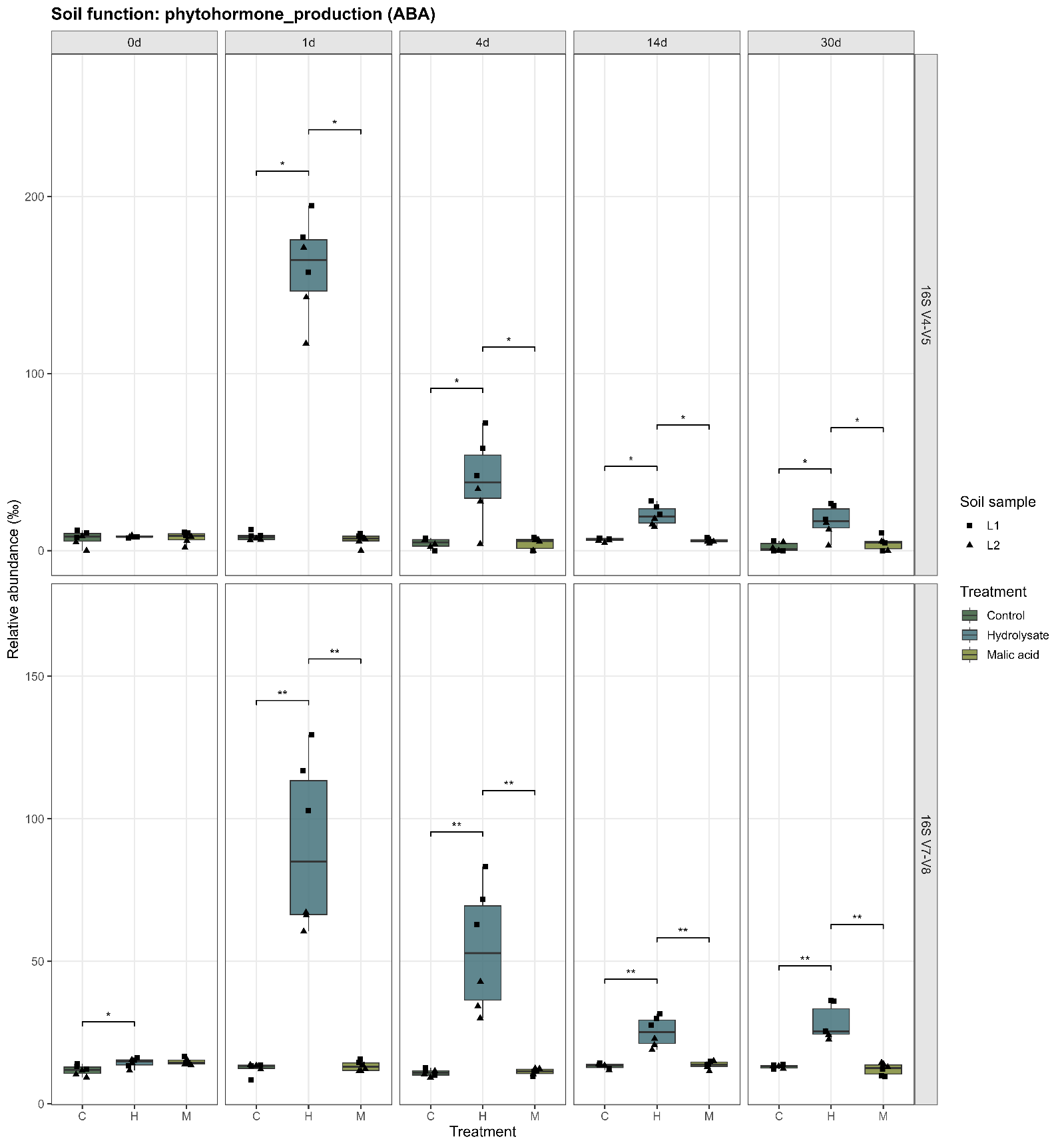

**Figure S26.** Temporal dynamics of the abundance of bacterial taxa classified as putative ABA producers. A) Inferred from the sequencing subset “set1” (variable regions V4 and V5 of the 16S) or subset “set2” (variable regions V7 and V8 of the 16S), see Table S6 for details.

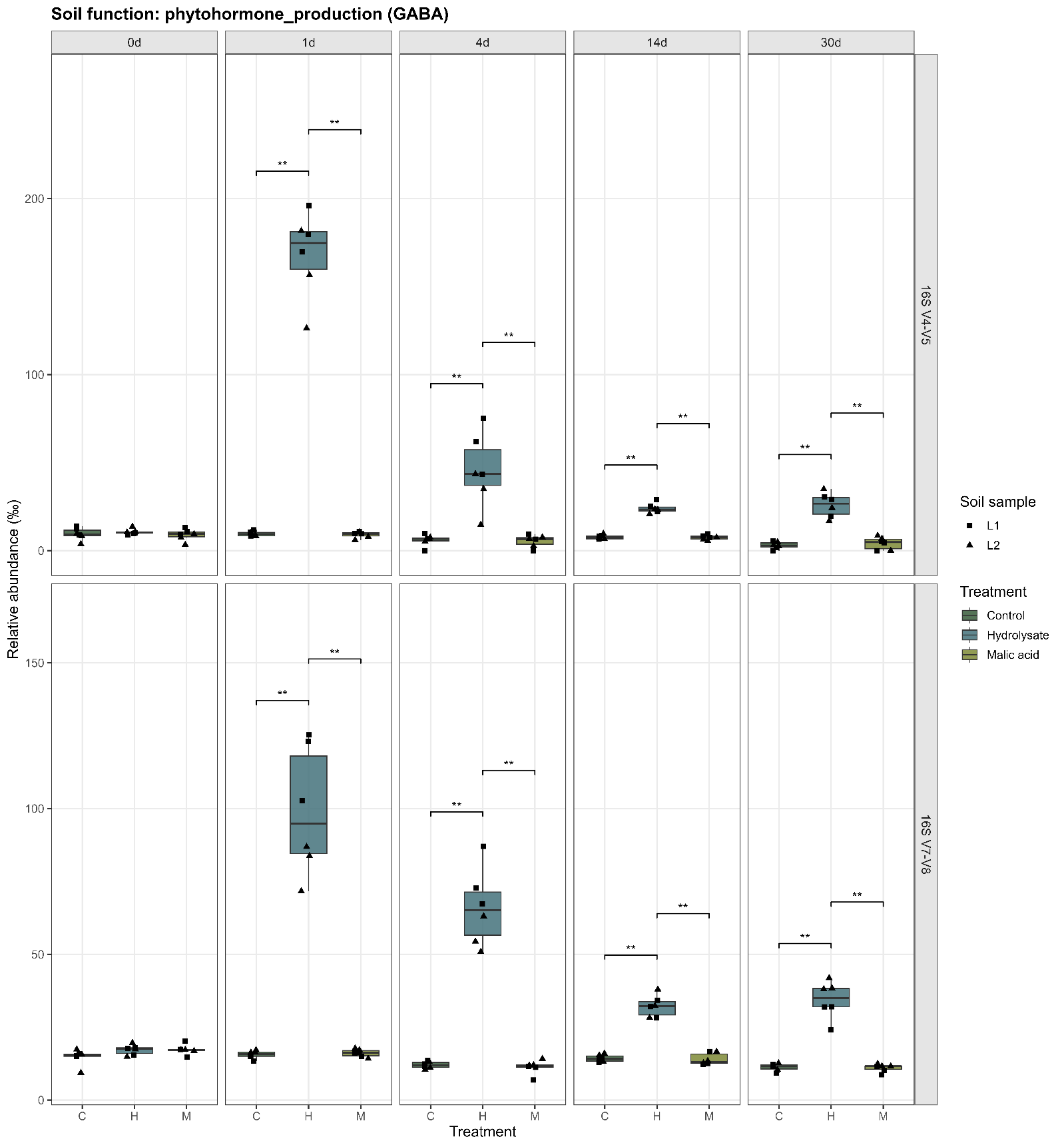

**Figure S27.** Temporal dynamics of the abundance of bacterial taxa classified as putative GABA producers. A) Inferred from the sequencing subset “set1” (variable regions V4 and V5 of the 16S) or subset “set2” (variable regions V7 and V8 of the 16S), see Table S6 for details.

A

A

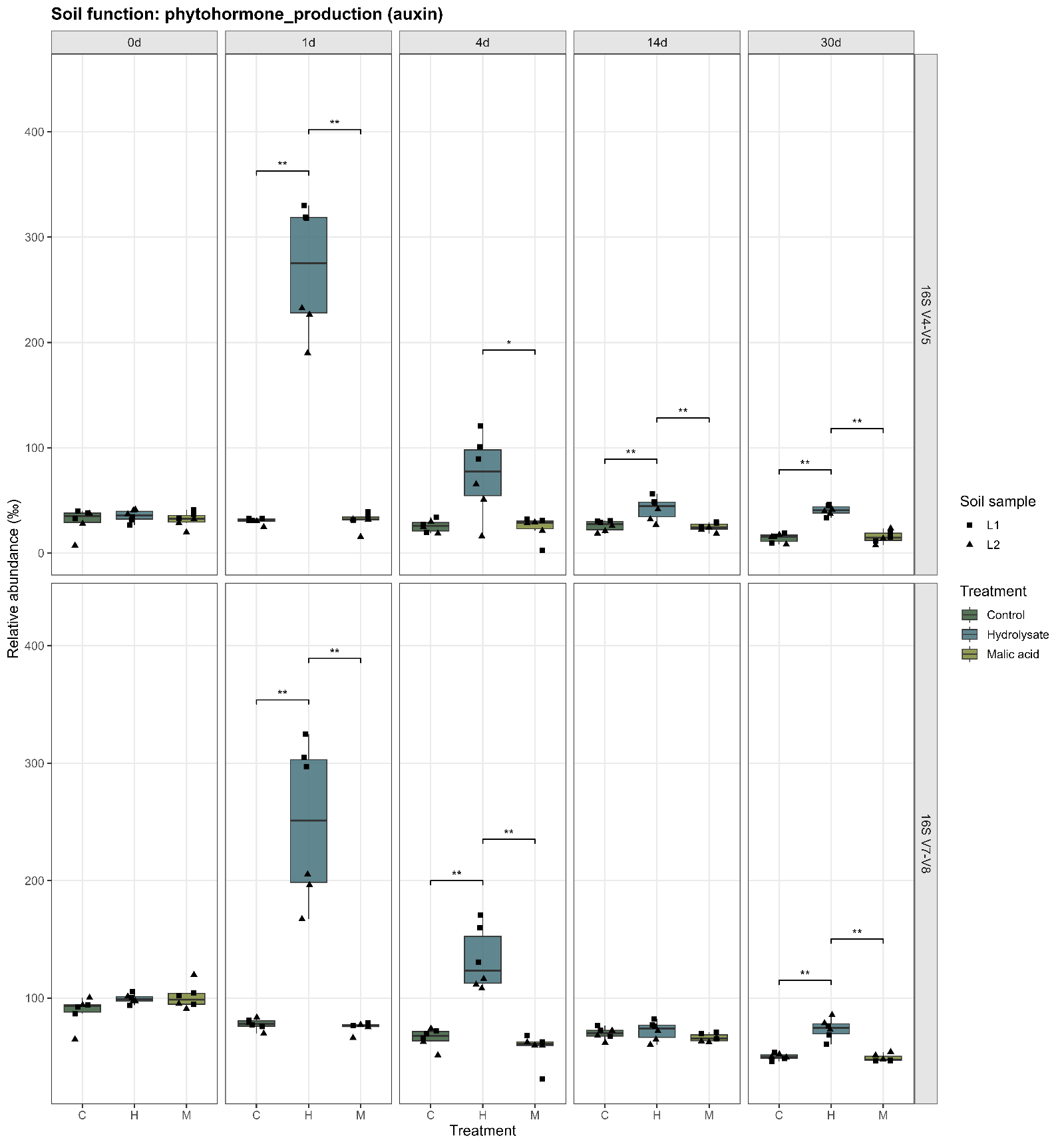

**Figure S28.** Temporal dynamics of the abundance of bacterial taxa classified as putative auxin producers. A) Inferred from the sequencing subset “set1” (variable regions V4 and V5 of the 16S) or subset “set2” (variable regions V7 and V8 of the 16S), see Table S6 for details.

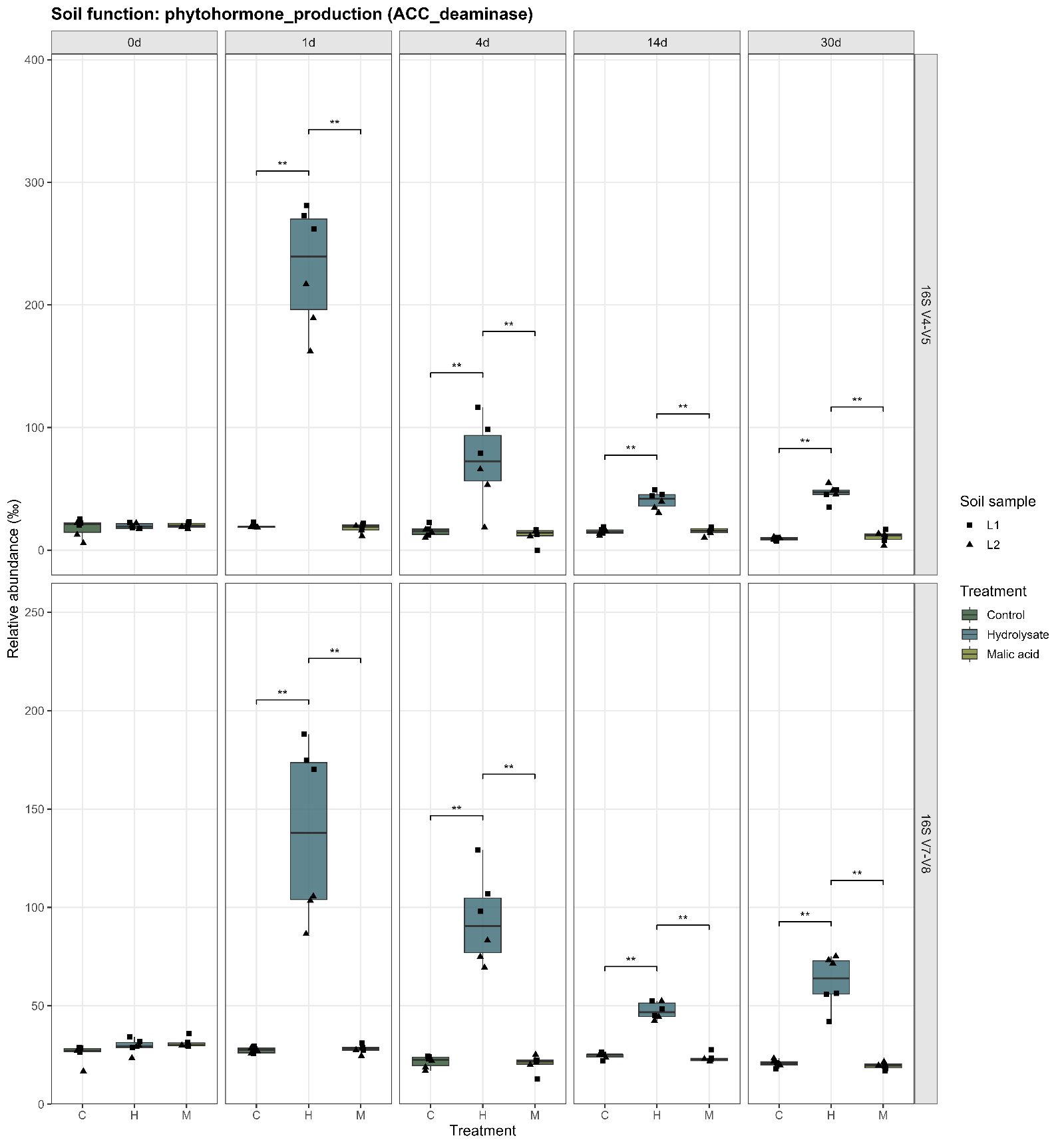

**Figure S29.** Temporal dynamics of the abundance of bacterial taxa classified as putative ACC deaminase producers. A) Inferred from the sequencing subset “set1” (variable regions V4 and V5 of the 16S) or subset “set2” (variable regions V7 and V8 of the 16S), see Table S6 for details.

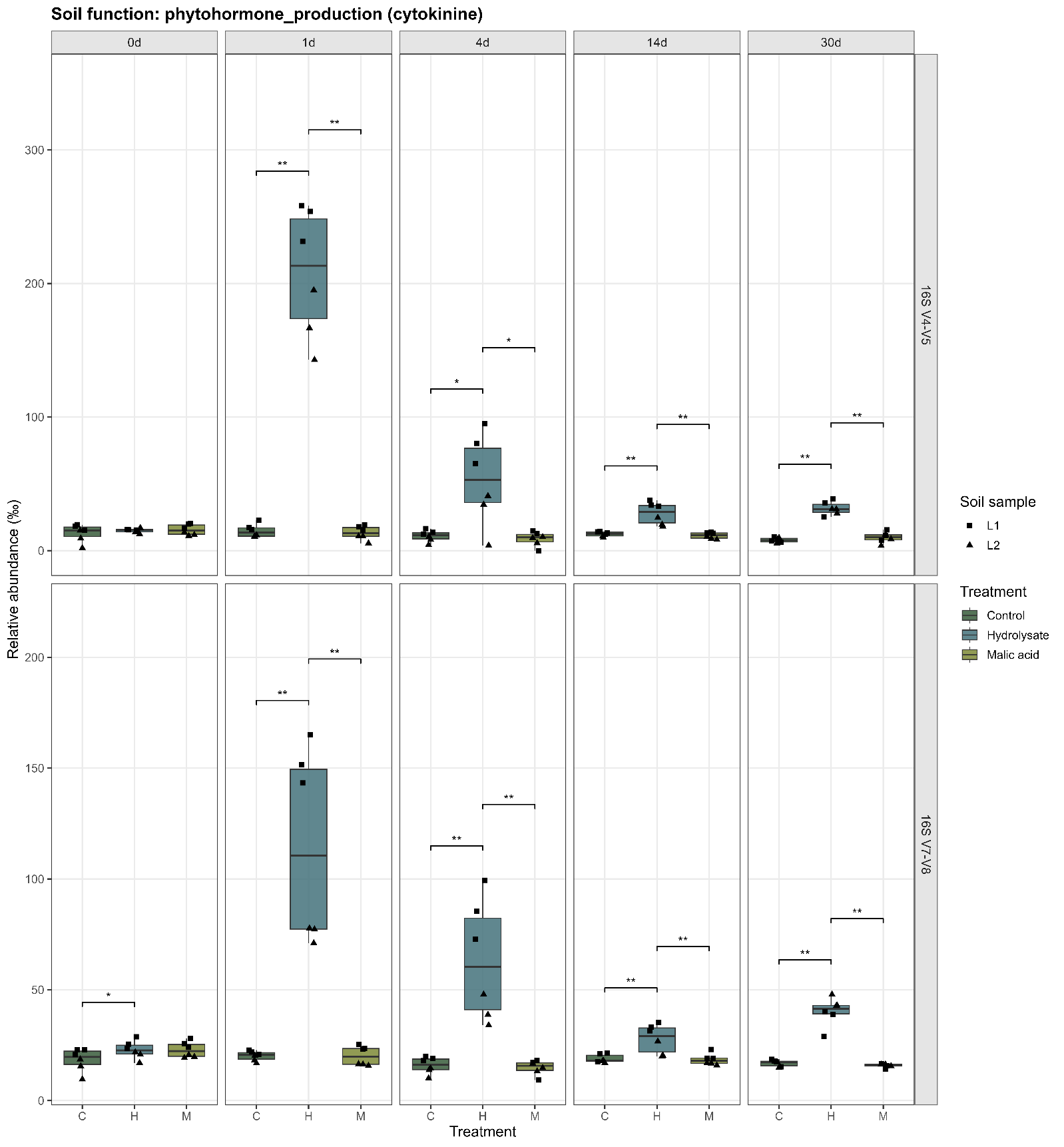

**Figure S30.** Temporal dynamics of the abundance of bacterial taxa classified as putative cytokinine producers. A) Inferred from the sequencing subset “set1” (variable regions V4 and V5 of the 16S) or subset “set2” (variable regions V7 and V8 of the 16S), see Table S6 for details.

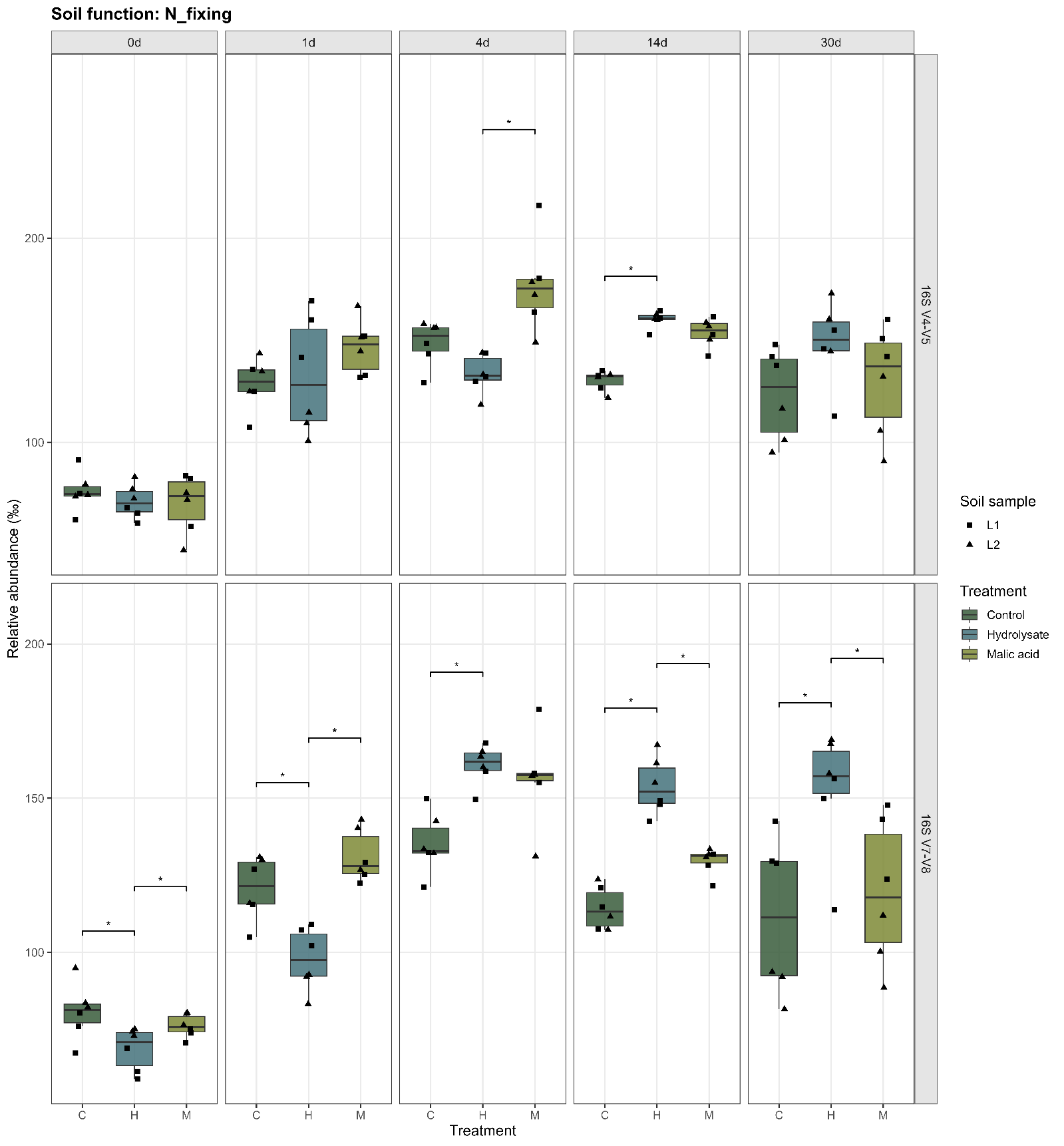

**Figure S31.** Temporal dynamics of the abundance of bacterial taxa classified as putative nitrogen fixers. A) Inferred from the sequencing subset “set1” (variable regions V4 and V5 of the 16S) or subset “set2” (variable regions V7 and V8 of the 16S), see Table S6 for details.

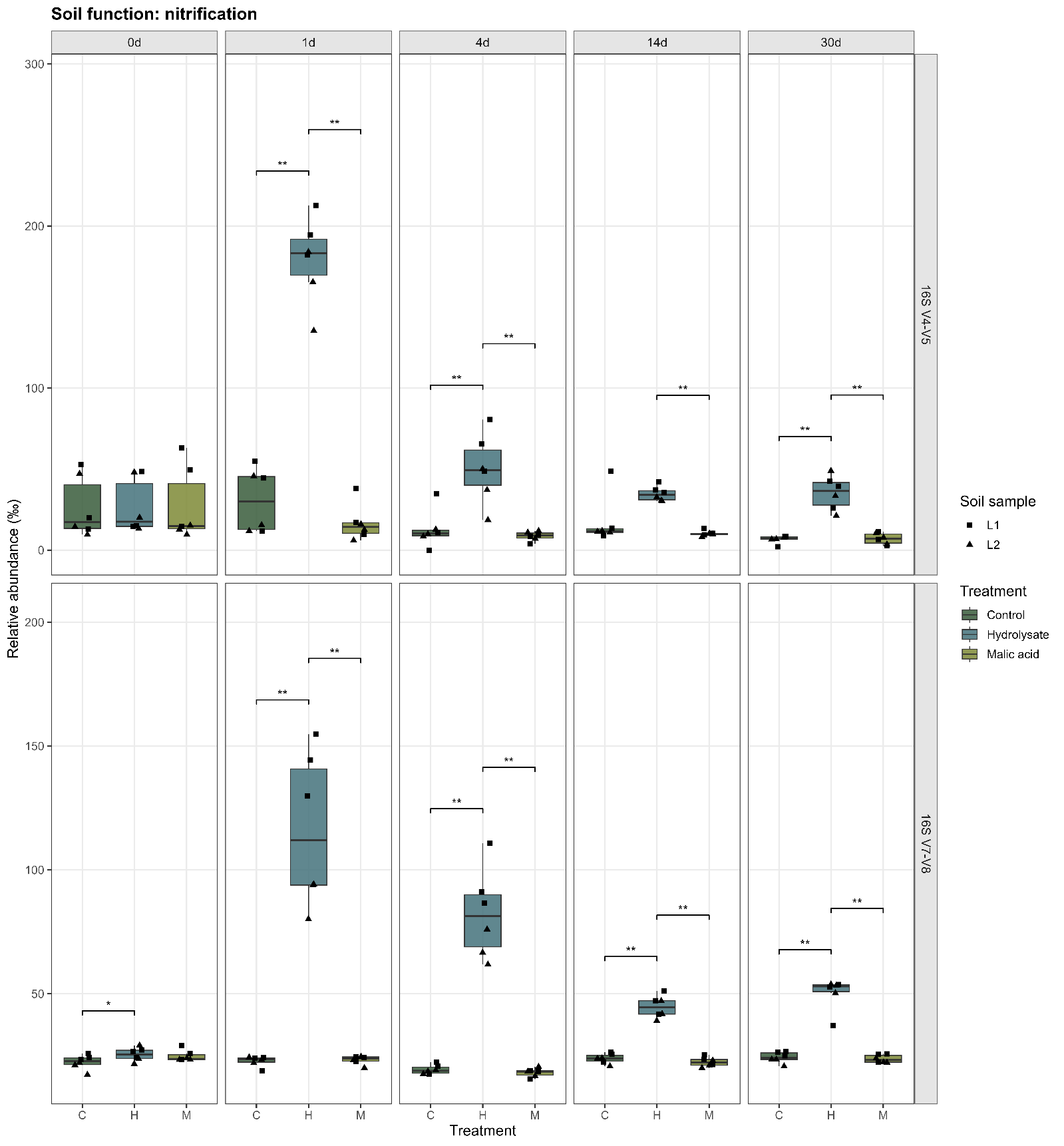

**Figure S32.** Temporal dynamics of the abundance of bacterial taxa classified as putative nitrifiers. A) Inferred from the sequencing subset “set1” (variable regions V4 and V5 of the 16S) or subset “set2” (variable regions V7 and V8 of the 16S), see Table S6 for details.

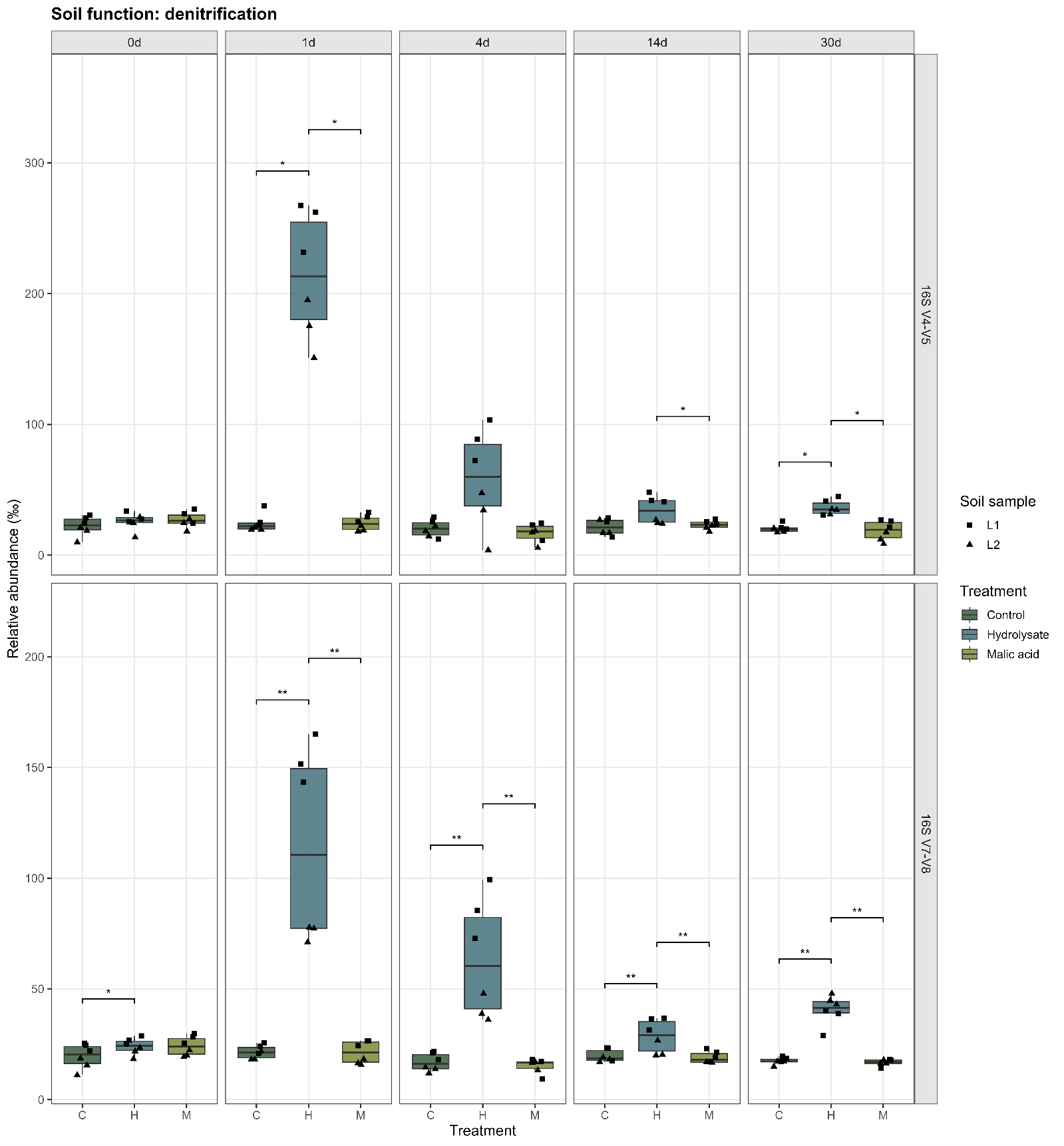

**Figure S33.** Temporal dynamics of the abundance of bacterial taxa classified as putative denitrifiers. A) Inferred from the sequencing subset “set1” (variable regions V4 and V5 of the 16S) or subset “set2” (variable regions V7 and V8 of the 16S), see Table S6 for details.

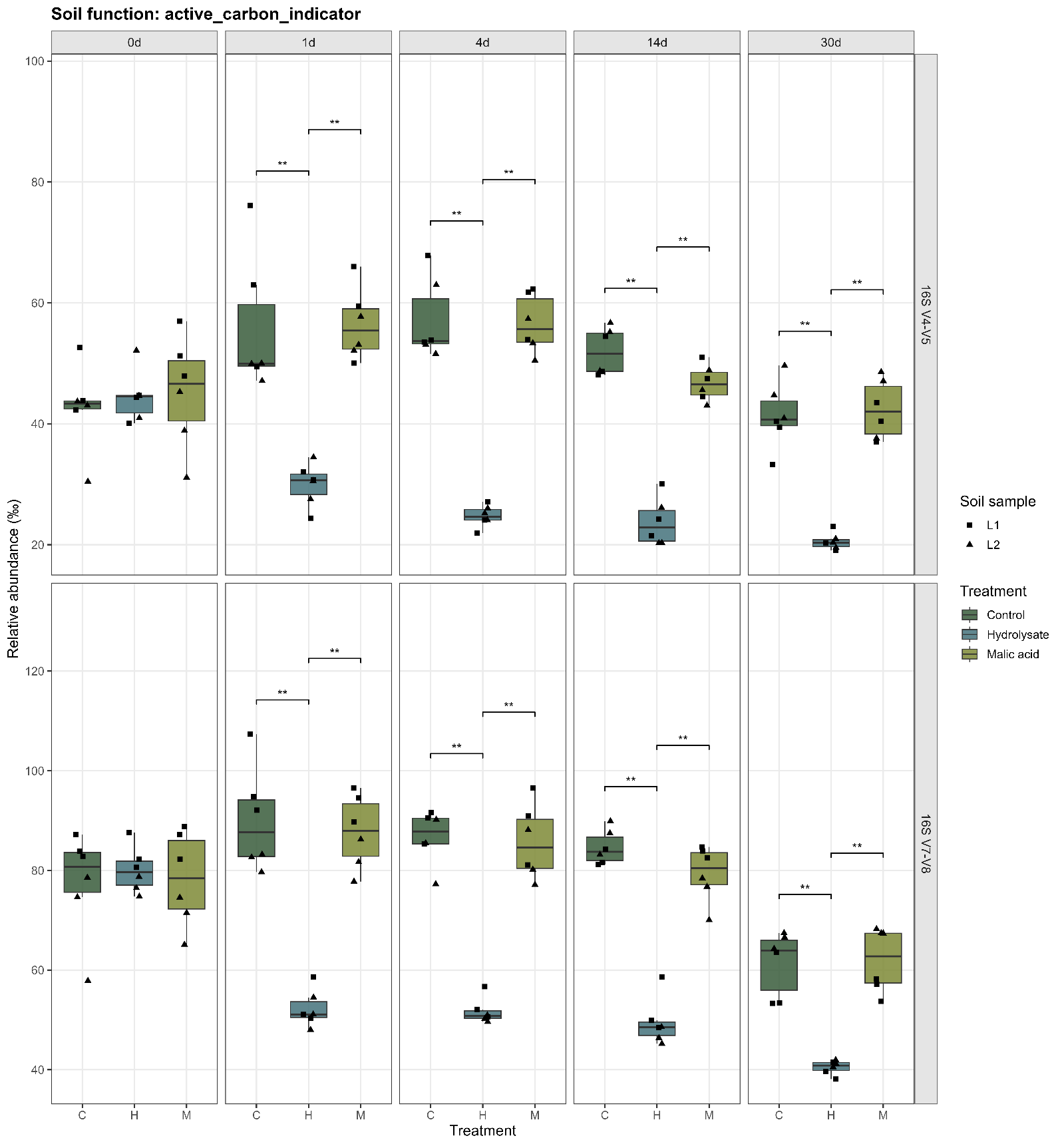

**Figure S34.** Temporal dynamics of the abundance of bacterial taxa classified as active carbon indicators. A) Inferred from the sequencing subset “set1” (variable regions V4 and V5 of the 16S) or subset “set2” (variable regions V7 and V8 of the 16S), see Table S6 for details.
